# Effects of germline and somatic events in candidate BRCAness genes on breast-tumor signatures

**DOI:** 10.1101/662874

**Authors:** Weston R. Bodily, Brian H. Shirts, Tom Walsh, Suleyman Gulsuner, Mary-Claire King, Alyssa Parker, Moom Roosan, Stephen R. Piccolo

## Abstract

**Background:** Mutations in *BRCA1* and *BRCA2* cause deficiencies in homologous recombination repair (HR), resulting in repair of DNA double-strand breaks by the alternative non-homologous end-joining pathway, which is more error prone. HR deficiency of breast tumors is important because it is associated with better responses to platinum salt therapies and PARP inhibitors. Among other consequences of HR deficiency are characteristic somatic-mutation signatures and gene-expression patterns. The term “BRCAness” describes tumors that harbor an HR defect but have no detectable germline mutation in *BRCA1* or *BRCA2*. A better understanding of the genes and molecular events associated with BRCAness could provide mechanistic insights and guide development of targeted treatments.

**Methods:** Using data from The Cancer Genome Atlas (TCGA) for 1101 breast-cancer patients, we identified individuals with a germline mutation, somatic mutation, homozygous deletion, and/or hypermethylation event in *BRCA1*, *BRCA2*, and 59 other cancer-predisposition genes. Based on the assumption that BRCAness events would have similar downstream effects on tumor biology as *BRCA1/BRCA2* germline mutations, we quantified these effects based on somatic-mutation signatures and gene-expression profiles. We reduced the dimensionality of the somatic-mutation signatures and expression data and used a statistical resampling approach to quantify similarities between patients who had a *BRCA1/BRCA2* germline mutation, another type of aberration in *BRCA1* or *BRCA2*, or any type of aberration in one of the other genes.

**Results:** Somatic-mutation signatures of tumors having a somatic mutation, homozygous deletion, or hypermethylation event in *BRCA1/BRCA2* (n = 80) were generally similar to each other and to tumors from *BRCA1/BRCA2* germline carriers (n = 44). Additionally, somatic-mutation signatures of tumors with germline or somatic events in *ATR* (n = 16) and *BARD1* (n = 8) showed high similarity to tumors from *BRCA1/BRCA2* carriers. Other genes also showed high similarity but only for a small number of events or for a single event type. Tumors with germline mutations or hypermethylation of *BRCA1* had relatively similar gene-expression profiles and overlapped considerably with the Basal-like subtype; but the transcriptional effects of the other events lacked consistency.

**Conclusions:** Our findings confirm previously known relationships between molecular signatures and germline or somatic events in *BRCA1/BRCA2* and suggest additional genes that may be considered for inclusion in the definition of BRCAness.

## Introduction

Approximately 1-5% of breast-cancer patients carry a pathogenic germline variant in either *BRCA1* or *BRCA2*^1–5^. These genes play important roles in homologous recombination repair (HR) of double-stranded breaks and stalled or damaged replication forks^6,7^. When the *BRCA1* or *BRCA2* gene products are unable to perform HR, cells may resort to non-homologous end-joining, a less effective means of repairing double-stranded breaks, potentially leading to an increased rate of DNA mutations^8–11^. Patients who carry biallelic loss of *BRCA1* and *BRCA2* due to germline variants and/or somatic events often respond well to *poly ADP ribose polymerase* (PARP) inhibitors and platinum-salt therapies, which increase the rate of DNA damage, typically causing the cells to enter programmed cell death^12–16^.

The downstream effects of BRCA mutations are distinctive. For example, BRCA-mutant tumors exhibit an abundance of C-to-T transitions across the genome, potentially reflecting tumor cells’ impaired ability to repair specific types of DNA damage^18^. In large-scale sequencing projects, such mutational patterns (termed *somatic-mutation signatures*) have been observed in association with other types of molecular events, as well as environmental and endogenous exposures, across many cancer types^19,20^. Among these signatures, the so-called “Signature 3” has been particularly associated with BRCA mutations and HR^19,21^.

Other downstream effects of BRCA mutations include characteristic transcriptional responses. For example, it has been shown that the “Basal” gene-expression subtype is enriched for tumors with *BRCA1* mutations^22–25^, that *BRCA1* mutations are commonly found in triple-negative breast tumors^26,27^, and that gene-expression profiles may predict PARP inhibitor responses^28^. These patterns are consistently observable, even in the presence of hundreds of other mutations^25,29^.

In 2004, Turner, et al. coined the term *BRCAness* to describe patients who do not have a pathogenic germline variant in *BRCA1* or *BRCA2* but who have developed a tumor with an impaired ability to perform HR^30^. This category may be useful for clinical management of patients and especially for predicting treatment responses^30,31^. Recent estimates suggest that the proportion of breast-cancer patients who fall into this category may be as high as 20%^32^. Davies, et al. demonstrated an ability to categorize patients into this category with high accuracy based on high-level mutational patterns^32^. Polak, et al. confirmed that somatic mutations, large deletions, and DNA hypermethylation of *BRCA1* and *BRCA2* are reliable indicators of BRCAness^21,33–35^. They also showed a relationship between BRCAness and germline mutations in *PALB2* and hypermethylation of *RAD51C*^21^. However, a considerable portion of breast tumors with HR deficiency lack a known driver. Furthermore, less is known about whether the downstream effects of germline variants, somatic variants, large deletions, and hypermethylation are similar to each other or whether these effects are similar for different genes.

An underlying assumption of the BRCAness concept is that the effects of HR deficiency are similar across tumors, regardless of the genes that drive those deficiencies and despite considerable variation in genetic backgrounds, environmental factors, and the presence of other driver mutations. Based on this assumption—and in a quest to identify additional genes that contribute to BRCAness—we performed a systematic evaluation of multiomic and clinical data from 1101 patients in The Cancer Genome Atlas (TCGA)^25^. In performing these evaluations, we characterized each tumor using two types of molecular signature: 1) weights that represent the tumor’s somatic-mutation profile and 2) the tumor’s mRNA expression profile. To evaluate similarities among tumors based on these molecular profiles, we used a statistical-resampling approach designed to quantify similarities among patient subgroups, even when those subgroups are small, thus helping to account for rare events. We use *aberration* as a general term to describe germline mutations, somatic mutations, copy-number deletions, and hypermethylation events.

## Methods

### Data preparation and filtering

We obtained breast-cancer data from TCGA for 1101 patients in total. To determine germline-mutation status, we downloaded raw sequencing data from CGHub^36^ for normal (blood) samples. We limited our analysis to whole-exome sequencing samples that had been sequenced using Illumina Genome Analyzer or HiSeq equipment. Because the sequencing data were stored in BAM format, we used Picard Tools (SamToFastq module, version 1.131, http://broadinstitute.github.io/picard) to convert the files to FASTQ format. We used the Burrows-Wheeler Alignment (BWA) tool (version 0.7.12)^37^ to align the sequencing reads to version 19 of the GENCODE reference genome (hg19 compatible)^38^. We used sambamba (version 0.5.4)^39^ to sort, index, mark duplicates, and flag statistics for the aligned BAM files. In cases where multiple BAM files were available for a single patient, we used bamUtil (version 1.0.13, https://github.com/statgen/bamUtil) to merge the BAM files. When searching for relevant germline variants, we examined 61 genes from a previous version of the BROCA Cancer Risk Panel (http://tests.labmed.washington.edu/BROCA)^40,41^. We extracted genomic data for these genes using bedtools (intersectBed module, version 2)^42^.

We used Picard Tools (CalculateHsMetrics module) to calculate alignment metrics. For exome-capture regions across all germline samples, the average sequencing coverage was 44.4. The average percentage of target bases that achieved at least 30X coverage was 33.7%. The average percentage of target bases that achieved at least 100X coverage was 12.3%.

To call DNA variants, we used freebayes (version v0.9.21-18-gc15a283)^43^ and Pindel (https://github.com/genome/pindel). We used freebayes to identify single-nucleotide variants (SNVs) and small insertions or deletions (indels); we used Pindel to identify medium-sized insertions and deletions. Having called these variants, we used snpEff (version 4.1)^44^ to annotate the variants and GEMINI (version 0.16.3)^45^ to query the variant data. To expedite execution of these steps, we used the GNU Parallel software^46^. The scripts and code that we used to process the germline data can be found in an open-access repository: https://bitbucket.org/srp33/tcga_germline.

Geneticists experienced in variant interpretation (BHS, TW, SG, MCK) further filtered the germline variants for pathogenicity using available sources of information on variants, following accepted guidelines for variant classification as previously described^47^. Accordingly, these germline calls were independent of variant-classification calls used in prior studies of TCGA data^25,48^. To assess loss of heterozygosity (LOH), we used data from Riaz, et al.^49^. They had made LOH calls for a large proportion of the breast-cancer patients in our study.

We identified somatic SNVs and indels for each patient by examining variant calls that had been made using Mutect^50^; these variants had been made available via the Genomic Data Commons^51^. We used the following criteria to exclude somatic variants: 1) synonymous variants 2) variants that snpEff classified as having a “LOW” or “MODIFIER” effect on protein sequence, 3) variants that SIFT^52^ and Polyphen2^53^ both suggested to be benign^54^, and 4) variants that were observed at greater than 1% frequency across all populations in ExAC^55^. For *BRCA1* and *BRCA2*, we examined candidate variants based on all available sources of evidence and the University of Washington, Department of Laboratory Medicine clinical database as described previously^56^. We compared our classifications to those publicly reported in the ClinVar database^57^ when available and found complete concordance. Based on these criteria, we categorized each variant as pathogenic, likely pathogenic, variant of uncertain significance (VUS), likely benign, or benign. Then we examined the ClinVar database^58^ for evidence that VUS or likely benign variants had been classified by others as pathogenic; however, none met this criterion. To err on the side of sensitivity, we considered any *BRCA1* and *BRCA2* mutation to be “aberrant” if it fell into our pathogenic, likely pathogenic, or VUS categories.

Using the somatic-mutation data for each patient, we derived mutation-signature profiles using the deconstructSigs (version 1.8.0) R package^59^. As input to this process, we used somatic-variant calls that had not been filtered for pathogenicity, as a way to ensure adequate representation of each signature. The output of this process was a vector for each tumor that indicated a weight for each signature^19^. Figures S1-S2 illustrate these weights for two tumors that we analyzed.

We downloaded DNA methylation data via the Xena Functional Genomics Explorer^60^. These data were generated using the Illumina HumanMethylation27 and HumanMethylation450 BeadChip platforms. For the HumanMethylation27 arrays, we mapped probes to genes using a file provided by the manufacturer (https://www.ncbi.nlm.nih.gov/geo/query/acc.cgi?acc=GPL8490). For the HumanMethylation450 arrays, we mapped probes to genes using an annotation file created by Price, et al.^61^ (see http://www.ncbi.nlm.nih.gov/geo/query/acc.cgi?acc=GPL16304). Typically, multiple probes mapped to a given gene. Using probe-level data from *BRCA1*, *BRCA2*, *PTEN*, and *RAD51C*, we performed a preliminary analysis to determine criteria for selecting and summarizing these probe-level values. We started with the assumption that in most cases, the genes would be methylated at low levels. We also assumed that probes nearest the transcription start site would be most informative. Upon plotting the data (Figure S3), we decided to limit our analysis to probes that mapped to the genome within 300 nucleotides of each gene’s transcription start site. In some cases, probes appeared to be faulty because they showed considerably different methylation levels (“beta” values) than other probes in the region (Figures S3A, S3C). To mitigate the effects of these outliers, we calculated gene-level methylation values as the median beta value across any remaining probes for that gene. To identify tumors that exhibited relatively high beta values—and thus could be considered to be *hypermethylated*—we used a univariate, outlier-detection algorithm, implemented in the extremevalues R packages (version 2.3.2)^62^. This enabled us to look for extreme values using one side of a specified distribution. This package supports five options for the distribution: normal (default), lognormal, exponential, pareto, and weibull. None of these distributions was a consistently good fit for the methylation (beta) values, in part because the shape of the data differed considerably across the genes (Figures S4-S7). We used the exponential distribution because it identified hypermethylated genes at rates that were largely consistent with prior work^21^. We used the *getOutliersII* function with default parameter values.

We downloaded copy-number-variation data from the Xena Functional Genomics Explorer^60^. These data had been generated using Affymetrix SNP 6.0 arrays; CNV calls had been made using the GISTIC2 method^63^. The CNV calls had also been summarized to gene-level values using integer-based discretization. We focused on tumors with a gene count of “-2”, which indicates a homozygous deletion.

We used RNA-Sequencing data that had been aligned and summarized to gene-level values using the original TCGA pipeline^25^. To facilitate biological and clinical interpretation, we examined relationships between germline and somatic events and the Prosigna™ Breast Cancer Prognostic Gene Signature (PAM50) subtypes^64^. Netanely, et al. had previously published PAM50 subtypes for TCGA breast cancer samples; we reused this information in our study^65^. We also sought to identify tumors with unusually low expression levels. To do this, we used the *getOutliersI* function in the extremevalues package. We used the following non-default parameter values: `alpha = c(0.000001, 0.000001), distribution = “lognormal”, FLim = c(0.1, 0.9)`. RNA-Sequencing data have been shown to fit the log-normal distribution^66^.

We parsed demographic, histopathological, and surgical variables for TCGA samples from the repository prepared by Rahman, et al.^67^. We obtained drug-response data from the TCGA legacy archive (https://portal.gdc.cancer.gov/legacy-archive) and standardized drug names using synonyms from the National Cancer Institute Thesaurus^68^.

### Quantitative analysis and visualization

To prepare, analyze, and visualize the data, we wrote computer scripts in the R programming language^69^. In writing these scripts, we used the following packages: readr^70^, dplyr^71^, ggplot2^72^, tidyr^73^, reshape2^74^, ggrepel^75^, cowplot^76^, data.table^77^, UpSetR^78^, BSgenome.Hsapiens.UCSC.hg38^79,80^, and Rtsne^81^.

To reduce data dimensionality for visualization purposes, we applied multidimensional scaling (MDS)^82^ to the somatic-mutation signatures and gene-expression profiles. This reduced the data to two dimensions. To quantify homogeneity within a group of tumors that harbored a particular aberration, we calculated the pairwise Euclidean distance between each patient pair in the group and then calculated the median pairwise distance, based on the two-dimensional values^83^. As an additional measure of homogeneity, we used logistic regression to predict BRCA mutation status using the dimensionally reduced coordinates. In evaluating this approach, we used two-fold cross validation and configured the model to weight the minority class in inverse proportion to the frequency of the minority class.

When comparing two given groups, we used a similar resampling approach but instead calculated the median distance between each pair of individuals in either group. To determine whether the similarity within or between groups was statistically significant, we used a permutation approach. We randomized the patient identifiers, calculated the median pairwise distance within (or between) groups, and repeated these steps 100,000 times. This process resulted in an empirical null distribution against which we compared the actual median distance. We then derived empirical p-values by calculating the proportion of randomized median distances that were larger than the actual median distance. We adjusted the empirical p-values for multiple testing using Holm’s method^84^; we applied this correction to the p-values across all aberration types.

For visualization, we plotted the MDS values as Cartesian coordinates; we also used Barnes-Hut *t*-distributed Stochastic Neighbor Embedding (*t*-SNE)^85,86^ as an alternative method for visualizing the data. We created a series of R scripts that execute all steps of our analysis and that generate the figures in this paper; these scripts are available at https://osf.io/9jhr2.

## Results

Initially, we used clinical and molecular data from breast-cancer patients in TCGA to evaluate the downstream effects of *BRCA1* and *BRCA2* germline mutations. We evaluated two types of downstream effect: 1) signatures that reflect a tumor’s overall somatic-mutation profile in a trinucleotide context and 2) tumor gene-expression levels. We used somatic-mutation signatures because they reflect the genomic effects of HR defects and have been associated with *BRCA1*/*BRCA2* mutation status^18,19^. We used gene-expression data because they are used to classify breast tumors into subtypes^87,88^ and often reflect genomic variation^54,89,90^. We assessed whether either of these profiles were more homogeneous in *BRCA1*/*BRCA2* germline carriers than in randomly selected patients. Next we evaluated potential criteria for classifying tumors into the “BRCAness” category. These criteria included somatic mutations, homozygous deletions, and DNA hypermethylation of *BRCA1* and *BRCA2*. Similarly, we assessed whether these types of aberration in 59 other cancer-predisposition genes have similar effects to *BRCA1*/*BRCA2* aberrations.

Of 993 breast-cancer patients with available germline data, 22 harbored a pathogenic SNV or indel in *BRCA1*; 22 harbored a *BRCA2* variant (Figure 1A). We were able to identify loss of heterozygosity (LOH) for all but 3 *BRCA1* carriers and 7 *BRCA2* carriers (Figures S8-S9). *BRCA1* carriers fell into the Basal (n = 17); Her2-enriched (n = 1), Luminal A (n = 2), and Luminal B (n = 1) gene-expression subtypes (Figure S10)^22,87,88^; we were unable to assign a gene-expression subtype to one BRCA1 carrier due to missing data. Most *BRCA2* carriers fell into the Luminal A subtype (n = 13); the remaining individuals were dispersed across the other subtypes. As demonstrated previously^19^, the primary somatic-mutation signature for most *BRCA1* and *BRCA2* carriers was “Signature 3”; however, other signatures (especially 1A) were also common (Figure S11). Figure S12 shows the overlap between these two types of molecular profile.

**Figure 1:**
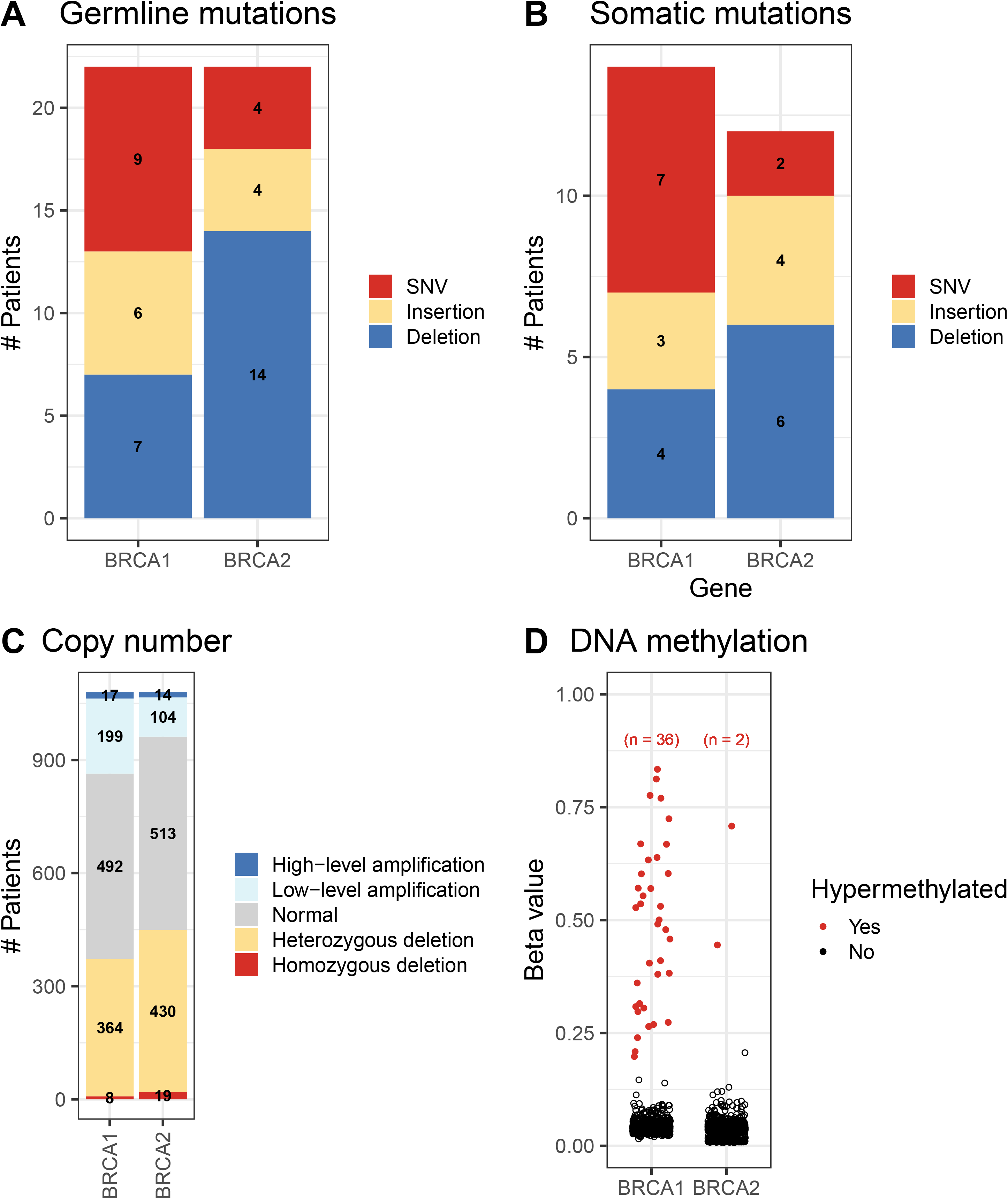
Molecular aberrations in *BRCA1* and *BRCA2* across all breast-cancer patients. A) Germline mutations, B) Somatic mutations, C) copy-number variations, D) DNA methylation levels. SNV = single nucleotide variation.

Although it is useful to evaluate breast-cancer patients based on the *primary* somatic-mutation signature or PAM50 subtype associated with each tumor, tumors are aggregates of multiple signatures and subtypes. To account for this diversity, we characterized tumors based on 1) all 27 somatic-mutation signatures or 2) expression levels for all available genes. To enable easier interpretation of these profiles, we reduced dimensionality of the data using the MDS and t-SNE techniques (see Methods). Generally, tumors with the same somatic-mutation signature or *primary* PAM50 subtype clustered together in these visualizations (Figures 2-3, S13-S14); however, in some cases, this did not happen. For example, the dimensionally reduced gene-expression profiles for Basal-like tumors formed a cluster that was mostly separate from the other tumors; but some Basal-like tumors were modestly distant from this cluster, and some Normal-like tumors clustered closely with the Basal-like tumors (Figures 3, S14). Tumors assigned to somatic-mutation “Signature 3” formed a cohesive cluster (Figures 2, S13), but some “Signature 3” tumors were modestly distant from this cluster. These observations highlight the importance of evaluating molecular profiles as a whole, not just using the primary category for each tumor.

**Figure 2:**
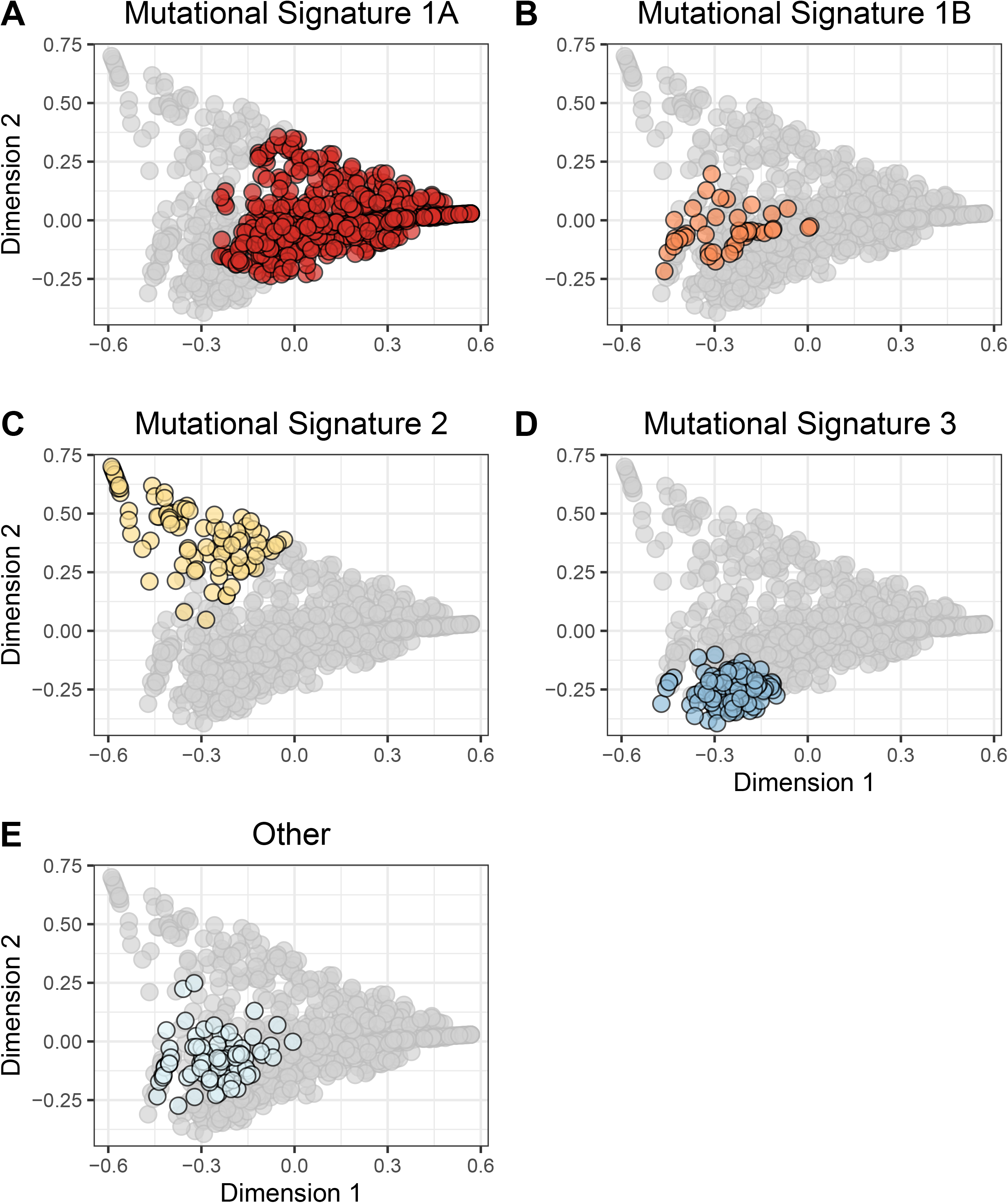
Two-dimensional representation of somatic-mutation signatures using multidimensional scaling. We summarized each tumor based on their somatic-mutation signatures, which represent overall mutational patterns in a trinucleotide context. We used multidimensional scaling (MDS) to reduce the data to two dimensions. Each point represents a single tumor, overlaid with colors that represent the tumor’s primary somatic-mutation signature. Mutational Signature 1A (A) was the most prevalent; these tumors were widely dispersed across the signature landscape. Signatures 1B (B), 2 (C), and 3 (D) were relatively small and formed cohesive clusters. The remaining 23 clusters were rare individually and were dispersed broadly.

**Figure 3:**
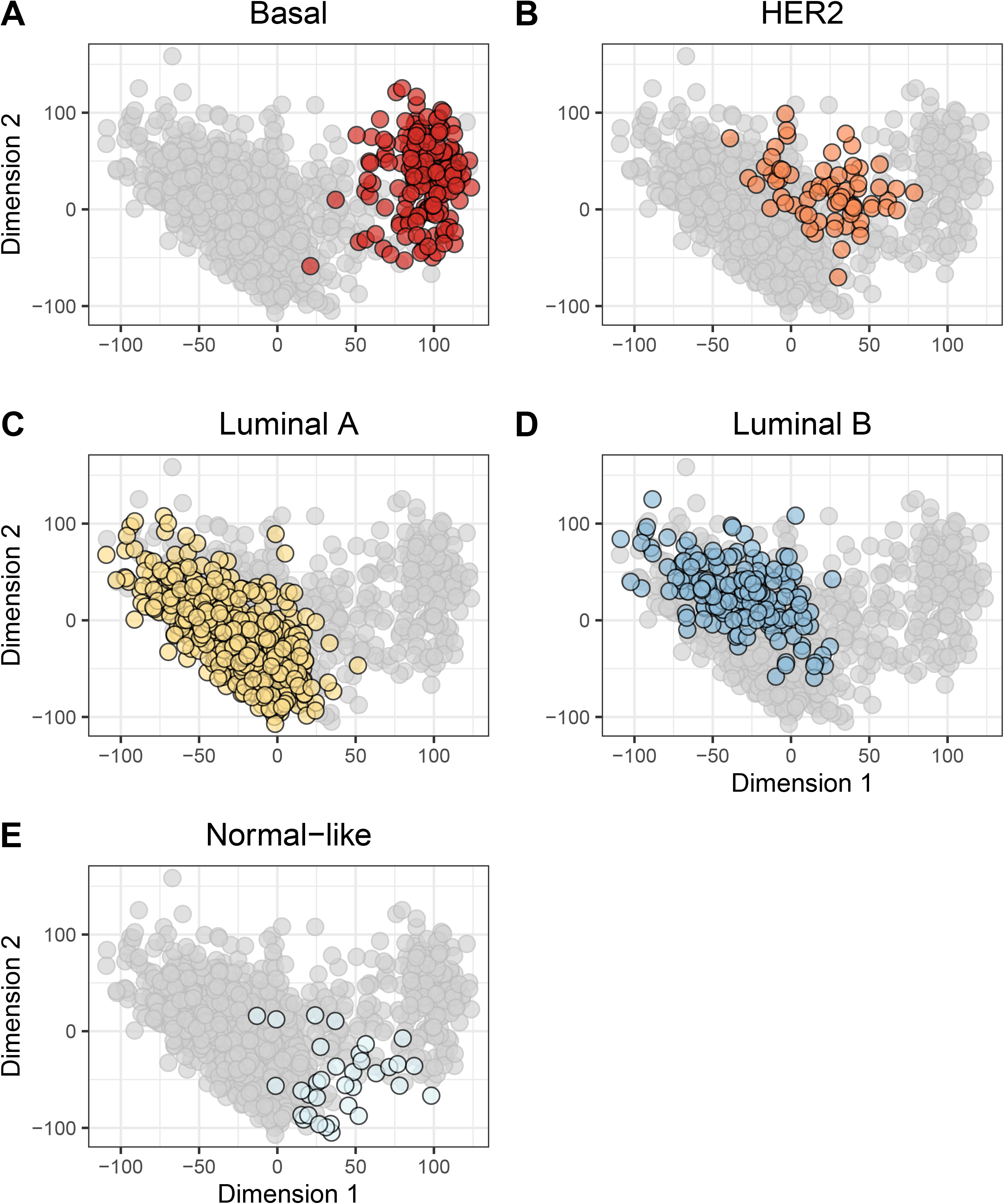
Two-dimensional representation of gene-expression levels using multidimensional scaling. We used multidimensional scaling (MDS) to reduce the gene-expression profiles to two dimensions. Each point represents a single tumor, overlaid with colors that represent the tumor’s primary PAM50 subtype. Generally, the PAM50 subtypes clustered cohesively, but there were exceptions. For example, some Basal-like tumors (A) exhbited expression patterns that differed considerably from the remaining Basal-like tumors. The normal-like tumors (E) showed the most variability in expression. This graph represents patients for whom we could identify a PAM50 subtype.

The somatic-mutation signatures of *BRCA1* germline carriers were *more* homogeneous than expected by chance (p = 0.00056; Figures S15A, S16A, S17A), as were those from *BRCA2* carriers (p = 0.0003; Figures S15B, S16B, S17B). As an additional measure of homogeneity, we used logistic regression to predict BRCA aberration status based on the dimensionally reduced data. Using somatic-mutation signatures, we could predict the presence of *BRCA1* germline mutations with a sensitivity of 0.72 and a specificity of 0.82. Additional classification results are shown in Table S1.

None of the three *BRCA1* carriers who lacked LOH events clustered closely with the remaining *BRCA1* tumors (Figures S15A, S16A). Of the 7 *BRCA2* tumors without detected LOH events, 3 clustered closely with the remaining *BRCA2* tumors, while 4 did not (Figures S15B, S16B). It has been shown previously that germline *BRCA1*/*BRCA2* mutations leave a recognizable imprint on a tumor’s mutational landscape^19^. This effect may be more consistent when a LOH event has occurred as a second “hit” within the same gene^91^, but this was difficult to confirm with the available sample sizes.

Under the assumption that *BRCA1*/*BRCA2* germline variants exhibit recognizable effects on tumor transcription, we assessed whether tumors from *BRCA1*/*BRCA2* carriers have homogeneous gene-expression profiles. As expected based on the tumors’ primary PAM50 classification, 17 of 21 *BRCA1* carriers (for whom we had gene-expression data) overlapped closely with the Basal-like subtype (Figure S10). As a whole, the expression profiles for this group were more homogeneous than expected by chance (p = 0.0318; Figures S18A, S19A, S20A). However, expression values for *BRCA2* carriers were *not* significantly homogeneous (p = 1.0; Figure S20B). Tumors from these individuals were dispersed across the gene-expression topography (Figures S18B, S19B).

Next we evaluated similarities between *BRCA1* and *BRCA2* germline carriers. Somatic-mutation signatures for *BRCA1* and *BRCA2* carriers were highly similar to each other (p = 0.00014; Figures S15A-S15B, S16A-S16B, S21A). However, this pattern did not hold for gene-expression profiles. Although some *BRCA2* carriers fell into the Basal-like gene-expression subtype, overall profiles for these patients were dissimilar to those from *BRCA1* carriers (p = 1.0; Figures S18A-S18B, S19A-S19B, S22A).

A somatic mutation, homozygous deletion, or hypermethylation event occurred in *BRCA1* and *BRCA2* for 80 patients (Figures 1B-1D). Most of these events were mutually exclusive with each other and with germline variants (Figure S23). Whether for somatic-mutation signatures or gene-expression profiles, tumors with *BRCA1* hypermethylation were relatively homogeneous and highly similar to tumors from *BRCA1* germline carriers (Table 1; Figures S15G, S16G, S17G, S18G, S19G, S20G). For gene-expression data, no other aberration type showed significant similarity to *BRCA1* germline mutations. Somatic-mutation signatures from tumors with *BRCA1* somatic mutations were significantly similar to those from *BRCA1* germline mutations (Table 1). Only 2 tumors had *BRCA2* hypermethylation, but the mutational signatures for these samples were significantly similar to tumors from *BRCA2* germline carriers (p = 0.00054; Figure S16H). Likewise, tumors with a *BRCA2* somatic mutation or homozygous deletion had mutational signatures that were similar to germline *BRCA2* carriers (Table 1; Figures S15D, S15F, S16D, S16F). Aberrations in *BRCA1* and *BRCA2* appear to exert similar effects on somatic-mutation signatures—but not necessarily gene expression—whether those disruptions originate in the germline or via somatic events.

**Table 1:** Results of similarity comparisons among BRCA aberration groups. We compared somatic-mutation signatures or gene-expression profiles between groups of patients who harbored aberrations in *BRCA1* or *BRCA2*. We evaluated whether patients in one group (e.g., those who harbored a *BRCA1* germline mutation) were more similar to patients in a second group (e.g., those with *BRCA2* germline mutation) than random patient subsets of the same sizes. The numbers in this table represent empirical p-values from our resampling approach. In cases where an individual harbored an aberration in both comparison groups, we excluded that patient from the comparison. We used Holm’s method to correct for testing multiple hypotheses^84^.

| **Aberration Type 1** | **Aberration Type 2** | **PAM50 Subtypes** | **Mutational Signatures** |
| --- | --- | --- | --- |
| BRCA1 germline mutation (n = 22) | BRCA2 germline mutation (n = 22) | 1.0 | 1.4e-04 |
| BRCA1 germline mutation (n = 22) | BRCA1 somatic mutation (n = 14) | 1.0 | 1.4e-04 |
| BRCA1 germline mutation (n = 22) | BRCA1 homozygous deletion (n = 8) | 1.0 | 0.30 |
| BRCA1 germline mutation (n = 22) | BRCA1 hypermethylation (n = 36) | 0.013 | 1.4e-04 |
| BRCA2 germline mutation (n = 22) | BRCA2 somatic mutation (n = 12) | 1.0 | 1.4e-04 |
| BRCA2 germline mutation (n = 22) | BRCA2 homozygous deletion (n = 19) | 1.0 | 1.4e-04 |
| BRCA2 germline mutation (n = 22) | BRCA2 hypermethylation (n = 2) | 1.0 | 5.4e-04 |

Next we aggregated all patients who had any type of *BRCA1* or *BRCA2* aberration into a *BRCAness reference group*. As a whole, mutational signatures for this group were more homogeneous than expected by chance (p = 0.00001; Figure S24). We used this reference group to evaluate 59 other cancer-predisposition genes that might be associated with BRCAness. For the remaining evaluations, we used somatic-mutation signatures only.

We evaluated whether molecular aberrations in the cancer-predisposition genes resulted in mutational signatures that were similar to our BRCAness reference group. We found pathogenic and likely pathogenic germline mutations in 13 genes (ATM, BARD1, BRCA1, BRCA2, BRIP1, CDH1, CHEK2, NBN, PALB2, PTEN, RAD51B, RAD51C, SLX4, TP53, XRCC2). The most frequently mutated were *CHEK2*, *ATM*, and *NBN* (Figures 4, S25-S26). We found potentially pathogenic somatic mutations in 55 genes, most frequently in *TP53*, *PIK3CA*, *CDH1*, and *PTEN* (Figures 5, S27-S28). Homozygous deletions occurred most frequently in *PTEN*, *CDKN2A*, *RB1*, and *CDH1* (Figures 6, S29-S30). Hypermethylation occurred in 22 genes, most commonly *GALNT12*, *PTCH1*, *CDKN2A*, and *RAD51C* (Figures 7, S31-S32). Using our resampling approach, we compared each aberration type in each gene against the BRCAness reference group. In cases where an aberration overlapped between the reference and comparison groups, we excluded individuals who harbored both aberrations. For 11 genes, at least one type of aberration attained statistical significance after multiple-testing correction (Table 2). A total of 8 aberrations occurred in *BARD1*: a germline mutation, 2 somatic mutations, and 5 homozygous deletions; as a group, these tumors were statistically similar to the BRCAness reference group (p = 0.0018). *ATR* was mutated in 15 tumors and hypermethylated in 1 tumor; together, these tumors were statistically similar to the BRCAness reference group (p = 0.0035). Tumors with an aberration in *PRSS1* were also statistically similar to the BRCAness reference group (p = 0.0069), but we only observed 2 aberrations in this gene. Other genes showed high similarity to the BRCAness group for one type of aberration only; these included homozygous deletions in *CDKN2A* (n = 47; p = 0.00001), homozygous deletions in *CTNNA1* (n = 6; p = 0.00004), germline mutations in *PALB2* (n = 3; p = 0.00001), and homozygous deletions in *PALLD* (n = 9; p = 0.00001).

**Table 2:**
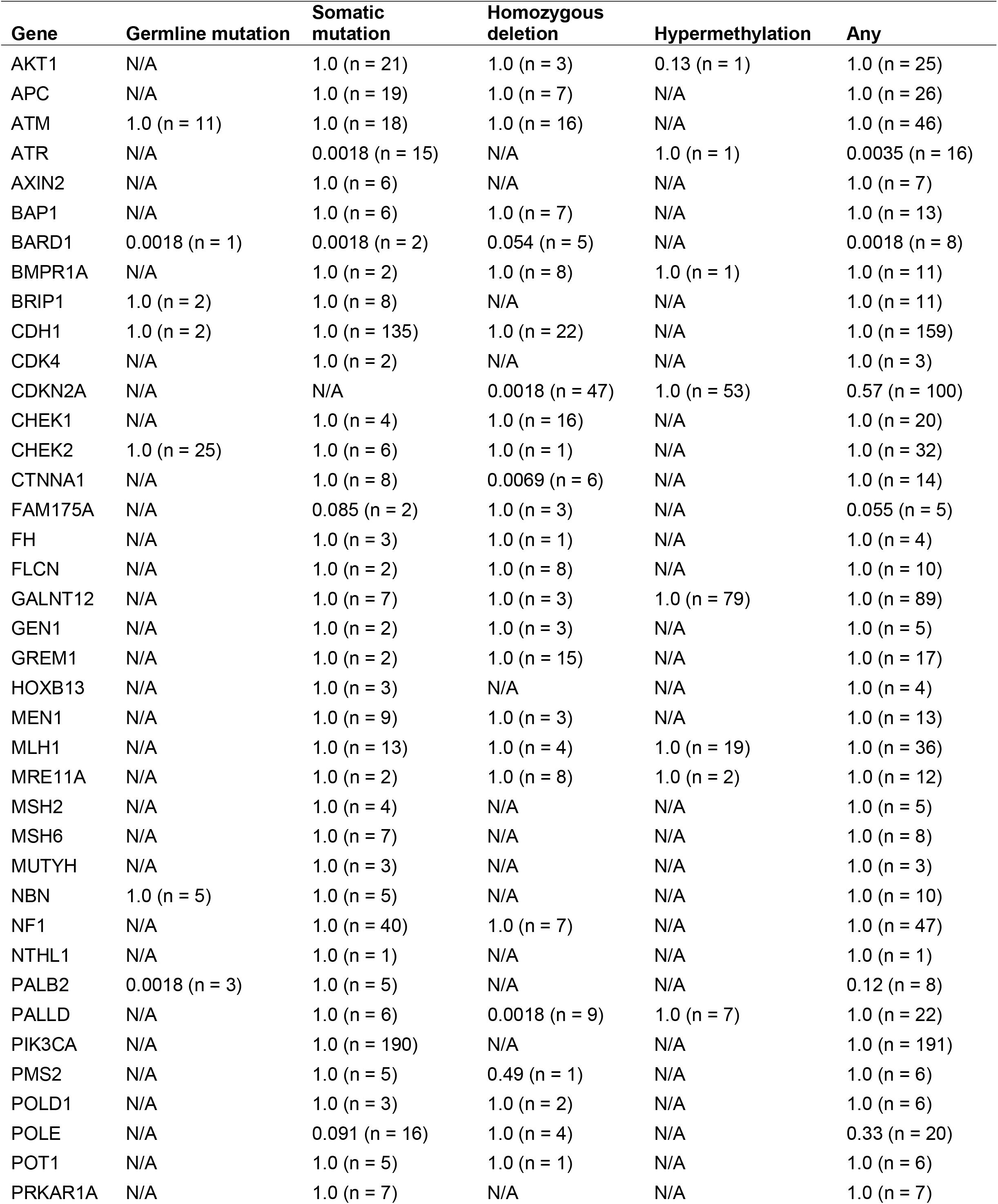
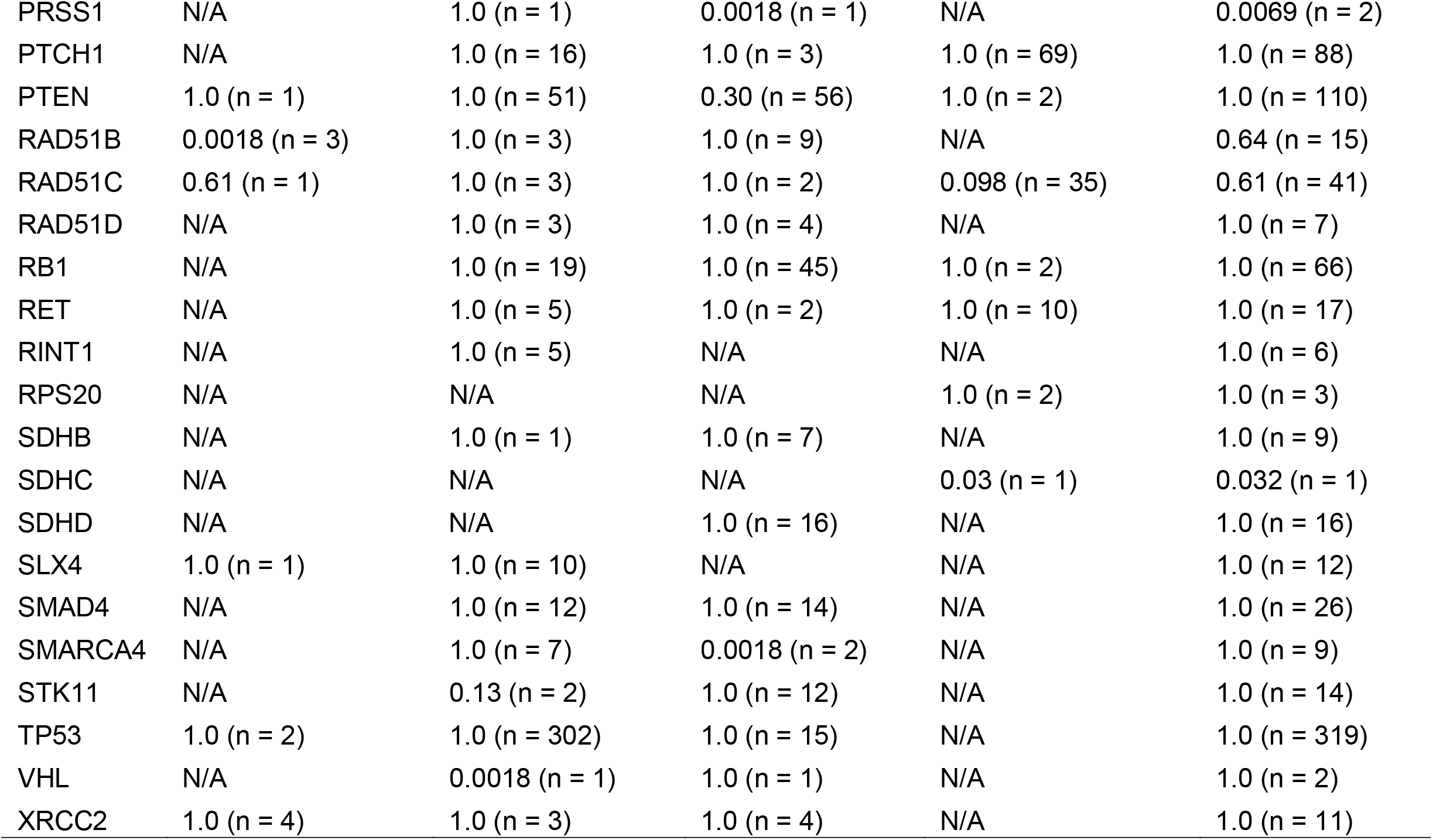
Summary of comparisons between the BRCAness reference group and groups of patients who harbored a specific type of aberration in a candidate BRCAness gene. We evaluated whether somatic-mutation signatures from patients who harbored a given type of aberration (e.g., BARD1 germline mutation) were more similar to the BRCAness reference group than expected by random chance. The numbers in this table represent empirical p-values from our resampling approach. In cases where no patient had a given type of aberration in a given gene, we list “N/A”. The “Any” group represents individuals who harbored any type of aberration in a given gene. We used Holm’s method to correct for testing multiple hypotheses^84^.

| Gene | Germline mutation | Somatic mutation | Homozygous deletion | Hypermethylation | Any |
| --- | --- | --- | --- | --- | --- |
| AKT1 | N/A | 1.0 (n = 21) | 1.0 (n = 3) | 0.13 (n = 1) | 1.0 (n = 25) |
| APC | N/A | 1.0 (n = 19) | 1.0 (n = 7) | N/A | 1.0 (n = 26) |
| ATM | 1.0 (n = 11) | 1.0 (n = 18) | 1.0 (n = 16) | N/A | 1.0 (n = 46) |
| ATR | N/A | 0.0018 (n = 15) | N/A | 1.0 (n = 1) | 0.0035 (n = 16) |
| AXIN2 | N/A | 1.0 (n = 6) | N/A | N/A | 1.0 (n = 7) |
| BAP1 | N/A | 1.0 (n = 6) | 1.0 (n = 7) | N/A | 1.0 (n = 13) |
| BARD1 | 0.0018 (n = 1) | 0.0018 (n = 2) | 0.054 (n = 5) | N/A | 0.0018 (n = 8) |
| BMPR1A | N/A | 1.0 (n = 2) | 1.0 (n = 8) | 1.0 (n = 1) | 1.0 (n = 11) |
| BRIP1 | 1.0 (n = 2) | 1.0 (n = 8) | N/A | N/A | 1.0 (n = 11) |
| CDH1 | 1.0 (n = 2) | 1.0 (n = 135) | 1.0 (n = 22) | N/A | 1.0 (n = 159) |
| CDK4 | N/A | 1.0 (n = 2) | N/A | N/A | 1.0 (n = 3) |
| CDKN2A | N/A | N/A | 0.0018 (n = 47) | 1.0 (n = 53) | 0.57 (n = 100) |
| CHEK1 | N/A | 1.0 (n = 4) | 1.0 (n = 16) | N/A | 1.0 (n = 20) |
| CHEK2 | 1.0 (n = 25) | 1.0 (n = 6) | 1.0 (n = 1) | N/A | 1.0 (n = 32) |
| CTNNA1 | N/A | 1.0 (n = 8) | 0.0069 (n = 6) | N/A | 1.0 (n = 14) |
| FAM175A | N/A | 0.085 (n = 2) | 1.0 (n = 3) | N/A | 0.055 (n = 5) |
| FH | N/A | 1.0 (n = 3) | 1.0 (n = 1) | N/A | 1.0 (n = 4) |
| FLCN | N/A | 1.0 (n = 2) | 1.0 (n = 8) | N/A | 1.0 (n = 10) |
| GALNT12 | N/A | 1.0 (n = 7) | 1.0 (n = 3) | 1.0 (n = 79) | 1.0 (n = 89) |
| GEN1 | N/A | 1.0 (n = 2) | 1.0 (n = 3) | N/A | 1.0 (n = 5) |
| GREM1 | N/A | 1.0 (n = 2) | 1.0 (n = 15) | N/A | 1.0 (n = 17) |
| HOXB13 | N/A | 1.0 (n = 3) | N/A | N/A | 1.0 (n = 4) |
| MEN1 | N/A | 1.0 (n = 9) | 1.0 (n = 3) | N/A | 1.0 (n = 13) |
| MLH1 | N/A | 1.0 (n = 13) | 1.0 (n = 4) | 1.0 (n = 19) | 1.0 (n = 36) |
| MRE11A | N/A | 1.0 (n = 2) | 1.0 (n = 8) | 1.0 (n = 2) | 1.0 (n = 12) |
| MSH2 | N/A | 1.0 (n = 4) | N/A | N/A | 1.0 (n = 5) |
| MSH6 | N/A | 1.0 (n = 7) | N/A | N/A | 1.0 (n = 8) |
| MUTYH | N/A | 1.0 (n = 3) | N/A | N/A | 1.0 (n = 3) |
| NBN | 1.0 (n = 5) | 1.0 (n = 5) | N/A | N/A | 1.0 (n = 10) |
| NF1 | N/A | 1.0 (n = 40) | 1.0 (n = 7) | N/A | 1.0 (n = 47) |
| NTHL1 | N/A | 1.0 (n = 1) | N/A | N/A | 1.0 (n = 1) |
| PALB2 | 0.0018 (n = 3) | 1.0 (n = 5) | N/A | N/A | 0.12 (n = 8) |
| PALLD | N/A | 1.0 (n = 6) | 0.0018 (n = 9) | 1.0 (n = 7) | 1.0 (n = 22) |
| PIK3CA | N/A | 1.0 (n = 190) | N/A | N/A | 1.0 (n = 191) |
| PMS2 | N/A | 1.0 (n = 5) | 0.49 (n = 1) | N/A | 1.0 (n = 6) |
| POLD1 | N/A | 1.0 (n = 3) | 1.0 (n = 2) | N/A | 1.0 (n = 6) |
| POLE | N/A | 0.091 (n = 16) | 1.0 (n = 4) | N/A | 0.33 (n = 20) |
| POT1 | N/A | 1.0 (n = 5) | 1.0 (n = 1) | N/A | 1.0 (n = 6) |
| PRKAR1A | N/A | 1.0 (n = 7) | N/A | N/A | 1.0 (n = 7) |
| PRSS1 | N/A | 1.0 (n = 1) | 0.0018 (n = 1) | N/A | 0.0069 (n = 2) |
| PTCH1 | N/A | 1.0 (n = 16) | 1.0 (n = 3) | 1.0 (n = 69) | 1.0 (n = 88) |
| PTEN | 1.0 (n = 1) | 1.0 (n = 51) | 0.30 (n = 56) | 1.0 (n = 2) | 1.0 (n = 110) |
| RAD51B | 0.0018 (n = 3) | 1.0 (n = 3) | 1.0 (n = 9) | N/A | 0.64 (n = 15) |
| RAD51C | 0.61 (n = 1) | 1.0 (n = 3) | 1.0 (n = 2) | 0.098 (n = 35) | 0.61 (n = 41) |
| RAD51D | N/A | 1.0 (n = 3) | 1.0 (n = 4) | N/A | 1.0 (n = 7) |
| RB1 | N/A | 1.0 (n = 19) | 1.0 (n = 45) | 1.0 (n = 2) | 1.0 (n = 66) |
| RET | N/A | 1.0 (n = 5) | 1.0 (n = 2) | 1.0 (n = 10) | 1.0 (n = 17) |
| RINT1 | N/A | 1.0 (n = 5) | N/A | N/A | 1.0 (n = 6) |
| RPS20 | N/A | N/A | N/A | 1.0 (n = 2) | 1.0 (n = 3) |
| SDHB | N/A | 1.0 (n = 1) | 1.0 (n = 7) | N/A | 1.0 (n = 9) |
| SDHC | N/A | N/A | N/A | 0.03 (n = 1) | 0.032 (n = 1) |
| SDHD | N/A | N/A | 1.0 (n = 16) | N/A | 1.0 (n = 16) |
| SLX4 | 1.0 (n = 1) | 1.0 (n = 10) | N/A | N/A | 1.0 (n = 12) |
| SMAD4 | N/A | 1.0 (n = 12) | 1.0 (n = 14) | N/A | 1.0 (n = 26) |
| SMARCA4 | N/A | 1.0 (n = 7) | 0.0018 (n = 2) | N/A | 1.0 (n = 9) |
| STK11 | N/A | 0.13 (n = 2) | 1.0 (n = 12) | N/A | 1.0 (n = 14) |
| TP53 | 1.0 (n = 2) | 1.0 (n = 302) | 1.0 (n = 15) | N/A | 1.0 (n = 319) |
| VHL | N/A | 0.0018 (n = 1) | 1.0 (n = 1) | N/A | 1.0 (n = 2) |
| XRCC2 | 1.0 (n = 4) | 1.0 (n = 3) | 1.0 (n = 4) | N/A | 1.0 (n = 11) |

**Figure 4:**
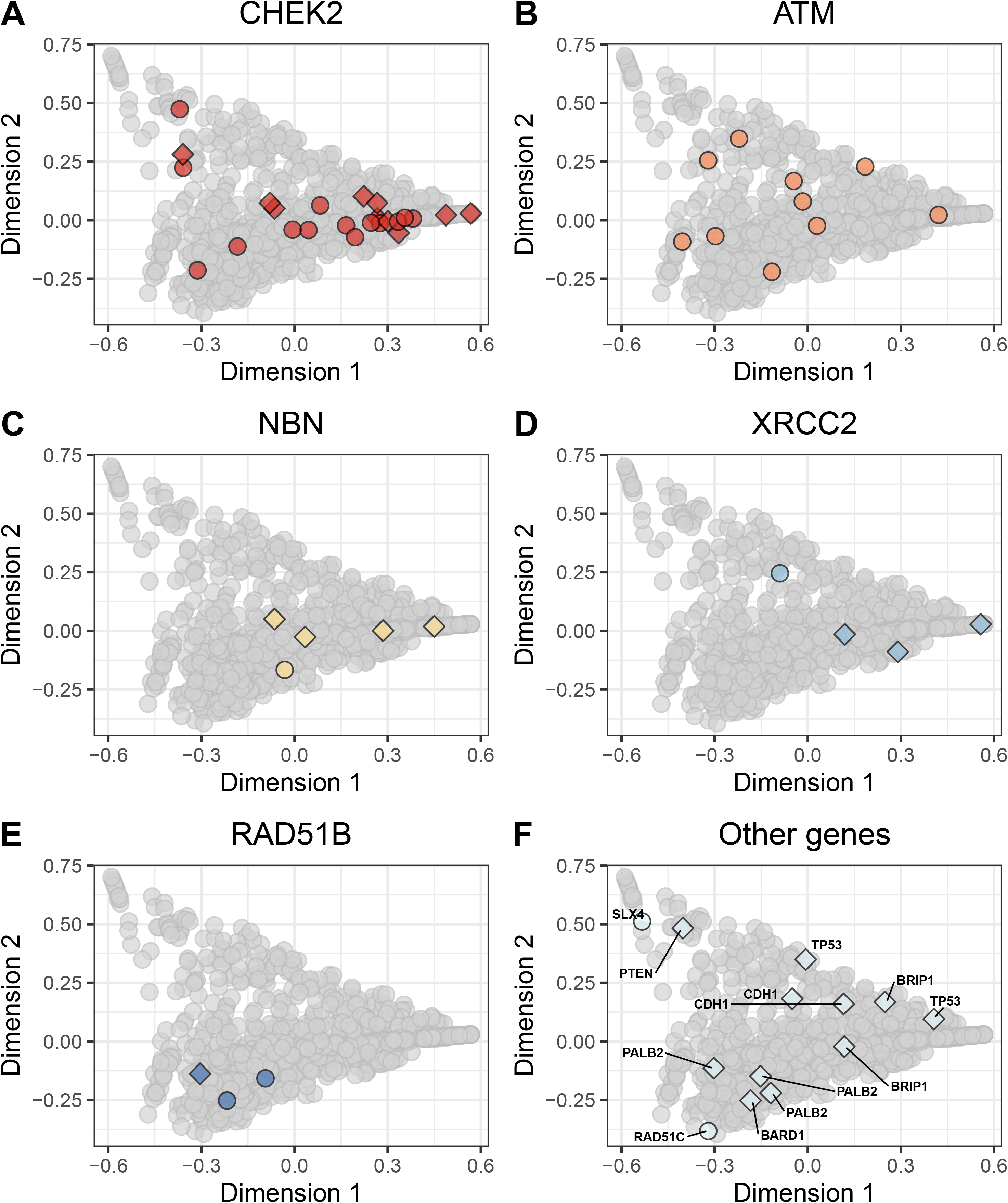
Non-BRCA germline mutations on the somatic-mutation signature landscape using multidimensional scaling. Using the same two-dimensional representation of mutational signatures shown in Figure 2, this plot indicates which patients had germline mutations in non-BRCA cancer-predisposition genes. Diamond shapes indicate patients for whom *no* loss-of-heterozygosity was observed.

**Figure 5:**
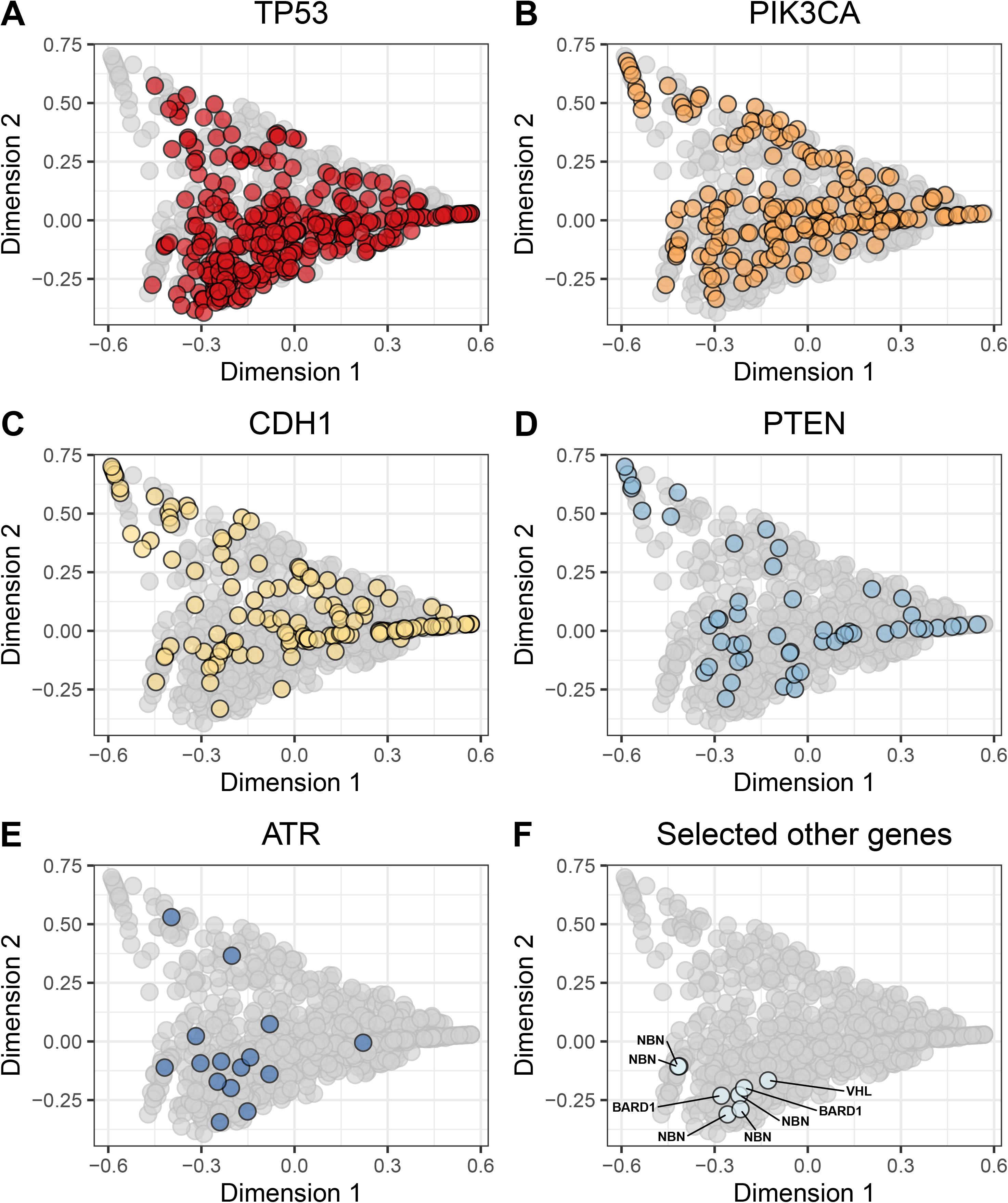
Non-BRCA somatic mutations on the somatic-mutation signature landscape using multidimensional scaling. Using the same two-dimensional representation of mutational signatures shown in Figure 2, this plot indicates which patients had somatic mutations in non-BRCA cancer-predisposition genes. Diamond shapes indicate patients for whom *no* loss-of-heterozygosity was observed.

**Figure 6:**
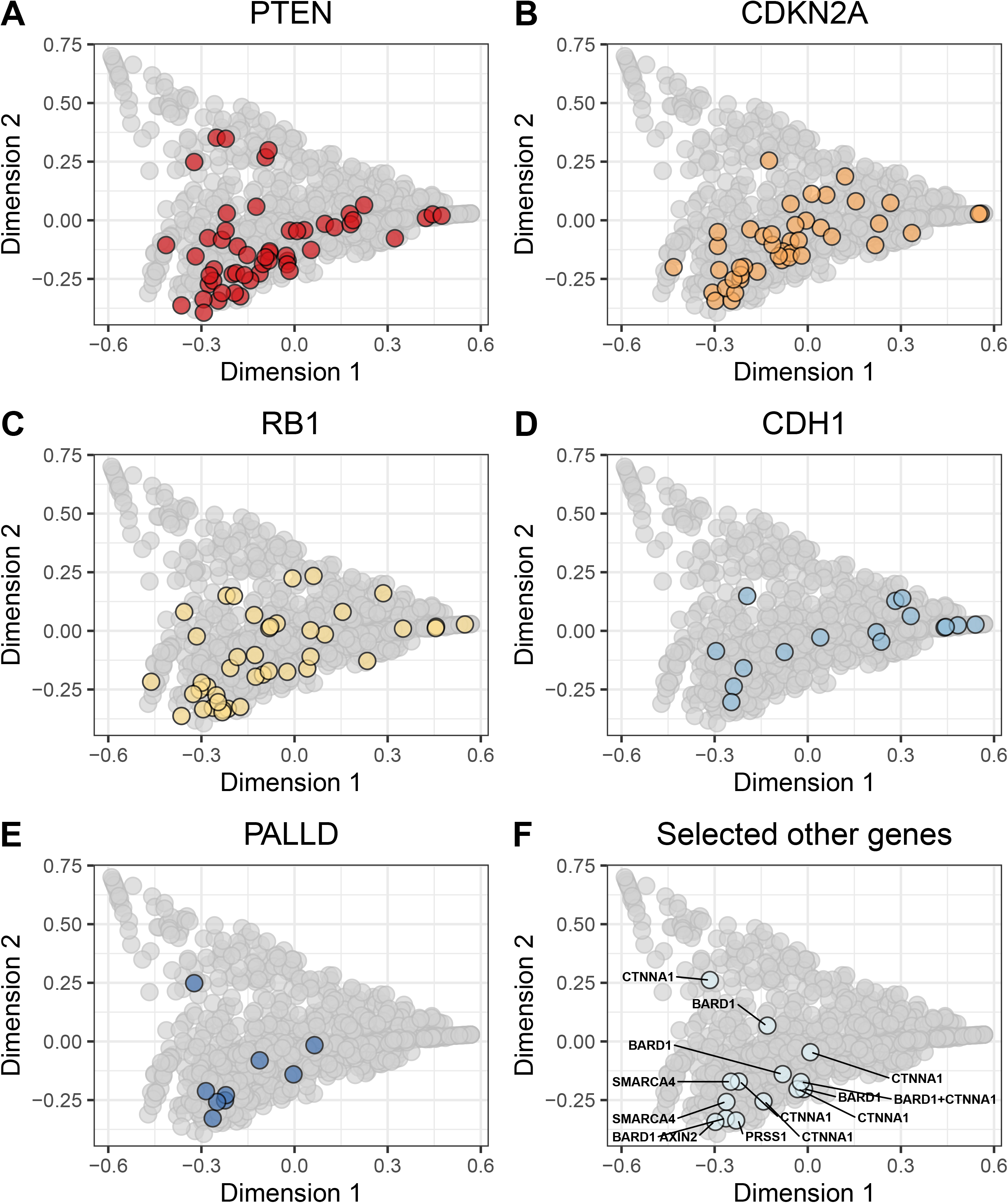
Non-BRCA homozygous deletions on the somatic-mutation signature landscape using multidimensional scaling. Using the same two-dimensional representation of mutational signatures shown in Figure 2, this plot indicates which patients had homozygous deletions in non-BRCA cancer-predisposition genes. Diamond shapes indicate patients for whom *no* loss-of-heterozygosity was observed.

**Figure 7:**
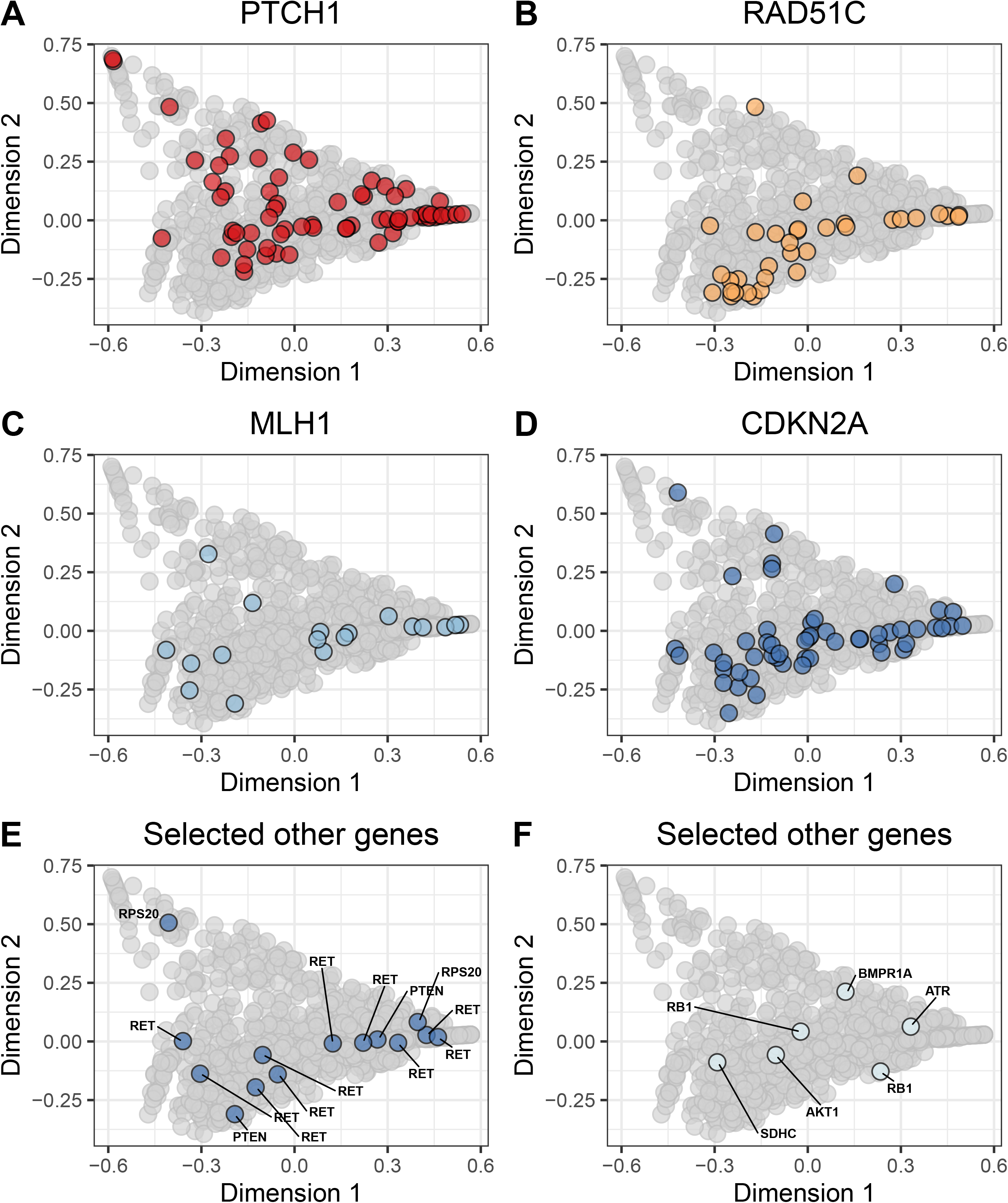
Non-BRCA hypermethylation events on the somatic-mutation signature landscape using multidimensional scaling. Using the same two-dimensional representation of mutational signatures shown in Figure 2, this plot indicates which patients had hypermethylation events in non-BRCA cancer-predisposition genes. Diamond shapes indicate patients for whom *no* loss-of-heterozygosity was observed.

Lastly, we evaluated whether the following data types were correlated with BRCAness: 1) unusually low mRNA expression in a given gene, 2) demographic, histopathological, and surgical observations, and 3) patient drug responses. First, we calculated the median Euclidean distance—based on somatic-mutation signatures—between each patient and the BRCAness reference group. Then we used a two-sided Pearson correlation test to assess the relationship between these median distances and each candidate variable. In determining whether a tumor exhibited unusually low mRNA expression for a given gene, we used an outlier-detection technique (see Methods). Unusually low expression of *BRCA1* (rho = 0.22, p = 0.0024) and *RAD51C* (rho = 0.20, p = 0.016) showed the strongest positive correlation with the reference group, whereas *CDH1* (rho = −0.19, p = 0.023), *PIK3CA* (rho = −0.19, p = 0.025) and *BARD1* (rho = −0.19, p = 0.028) showed the strongest negative correlation (Figures S33-S34). Triple-negative status and infiltrating ductal carcinoma histology were the most positively correlated clinical variables (Figure S35). No chemotherapy treatment was significantly associated with BRCAness, though availability was limited for the drug data (n = 211; Figure S36). Relationships among low mRNA expression and these factors are likely interwoven. For example, CDH1 expression has been associated with molecular subtypes such as triple-negative status^92^. Larger studies will be necessary to disentangle these effects.

## Discussion

Various criteria have been proposed as indicators of BRCAness, including triple-negative hormone-receptor status^93^, somatic mutations in *BRCA1*, hypermethylation of *BRCA1*, germline mutations in *PALB2*, and hypermethylation of *RAD51C*^21^. We evaluated these factors, as well as a wide range of other clinical and molecular factors, that could be indicators of BRCAness. We used a publicly available, multiomic dataset and robust, quantitative methods to evaluate the downstream effects of these aberrations on gene-expression and somatic-mutation signatures. Our permutation approach takes multiple variables (e.g. the full profile of signature weights) into account simultaneously, not just the primary subtype, for example. Although we observed a clear relationship between *BRCA1* aberrations and the “Basal-like” gene-expression subtype—which overlaps considerably with triple-negative status—we otherwise observed few consistent patterns in the gene-expression data. In contrast, we observed multiple significant results for the somatic-mutation signatures, many of which agree with findings from prior studies.

We found that germline *BRCA1* mutations affected somatic-mutation signatures similarly to germline *BRCA2* mutations. Furthermore, somatic-mutations, homozygous deletions, and hypermethylation of *BRCA1* and *BRCA2* had downstream effects that were overall similar to those of germline mutations in these genes. As a whole, tumors with any *BRCA1*/*BRCA2* aberration formed a cohesive group, against which we compared other tumors. For a gene to be considered a strong candidate to be included in the BRCAness definition, we required evidence of similarity across all available molecular data types. Two genes met these criteria: *BARD1* and *ATR*. Both genes interact directly with BRCA1 to help repair double-stranded breaks and control G1/S cell-cycle arrest^94^. Other genes—such as *CDKN2A*, *CTNNA1*, and *PALB2*—attained statistical significance for a single aberration type across a relatively large number of samples (Table 2) but failed to attain significance across all aberration types. Some genes known to play a role in DNA damage repair^95^-including *ATM*, *CHEK2*, and *RAD51C*—did not attain statistical significance for any aberration type. Different types of aberration may result in different downstream effects; however, these differences may result from technical challenges in identifying and filtering variants. Determining which aberrations are pathogenic remains a challenging task, so it is likely that more- or less-stringent filtering of candidate aberrations would lead to more accurate results. In addition, we could not always determine whether mono- or bi-allelic inactivation of a given gene had occurred in a given tumor; mono-allelic inactivation may be insufficient to impair HR function^49^.

To enable easier visualization, we reduced the molecular signatures to two dimensions. We also used the dimensionally reduced data as input to our statistical resampling approach. Accordingly, even though the number of original input variables was much larger for the gene-expression data, we could perform a more consistent comparison between the two data types. However, reducing the data dimensionality to this extent likely failed to capture much of the biological signal in the data for either data type. Further refining of this approach help to strike a better balance between dimensionality reduction and data interpretability.

The family-wise method we used to correct for multiple testing is generally conservative^84^. Accordingly, our results likely erred on the side of specificity rather than sensitivity for estimating whether a given gene should be considered a candidate for the BRCAness category. This may explain why *RAD51C* hypermethylation, for example, did not reach statistical significance in our analysis, even though it has been highlighted in other studies^21,32^.

By definition, BRCAness tumors have HR defects^30^. As with germline mutations in *BRCA1* and *BRCA2*, these deficiencies could be exploited therapeutically^15–17,96,97^. Although the patterns we observed were highly consistent in many cases, it remains to be determined whether these observations are clinically relevant. Clinical trials are currently underway to identify biomarkers for carboplatin, a platinum-salt agent. Tutt, et al. concluded that *BRCA1*/*BRCA2* mutations and triple-negative hormone status are reliable biomarkers of objective treatment responses but that *BRCA1* hypermethylation is not^93^. It may be that other BRCAness genes or different types of aberration will become useful markers of treatment response.

## Conclusions

Altogether our findings shed new light on factors that may be useful to classify genes into the BRCAness category and illustrate a quantitative methodology for evaluating tumor subtypes in general.

## Declarations

### Ethics approval and consent to participate

Brigham Young University’s Institutional Review Board approved this study under exemption status. This study uses data collected from public repositories only. We played no part in patient recruiting or in obtaining consent. We have adhered to guidelines from TCGA on handling data.

### Consent for publication

Not applicable.

### Availability of data and material

The datasets generated and analyzed during the current study are available in the Open Science Framework repository (https://osf.io/9jhr2). (We are not permitted to share the germline-mutation data.)

### Author’s contributions

WRB, BHS, and SRP conceived the study design, prepared and analyzed data, and interpreted results. BHS, TW, SG, and MCK evaluated variant pathogenicity and contributed intellectual insights regarding study design and data interpretation. AP and MR parsed and evaluated the pharmacological data. WRB and SRP wrote the manuscript. BHS, TW, MCK, AP and MR edited the manuscript.

## Acknowledgements

Results from this study are in part based upon data generated by TCGA and managed by the United States National Cancer Institute and National Human Genome Research Institute (see http://cancergenome.nih.gov). We thank the patients who participated in this study and shared their data publicly. We thank the Fulton Supercomputing Laboratory at Brigham Young University for providing computational facilities.

