## Supplementary Material for "Effects of germline and somatic events in candidate BRCAness genes on breast-tumor signatures"

### Supplementary Figures

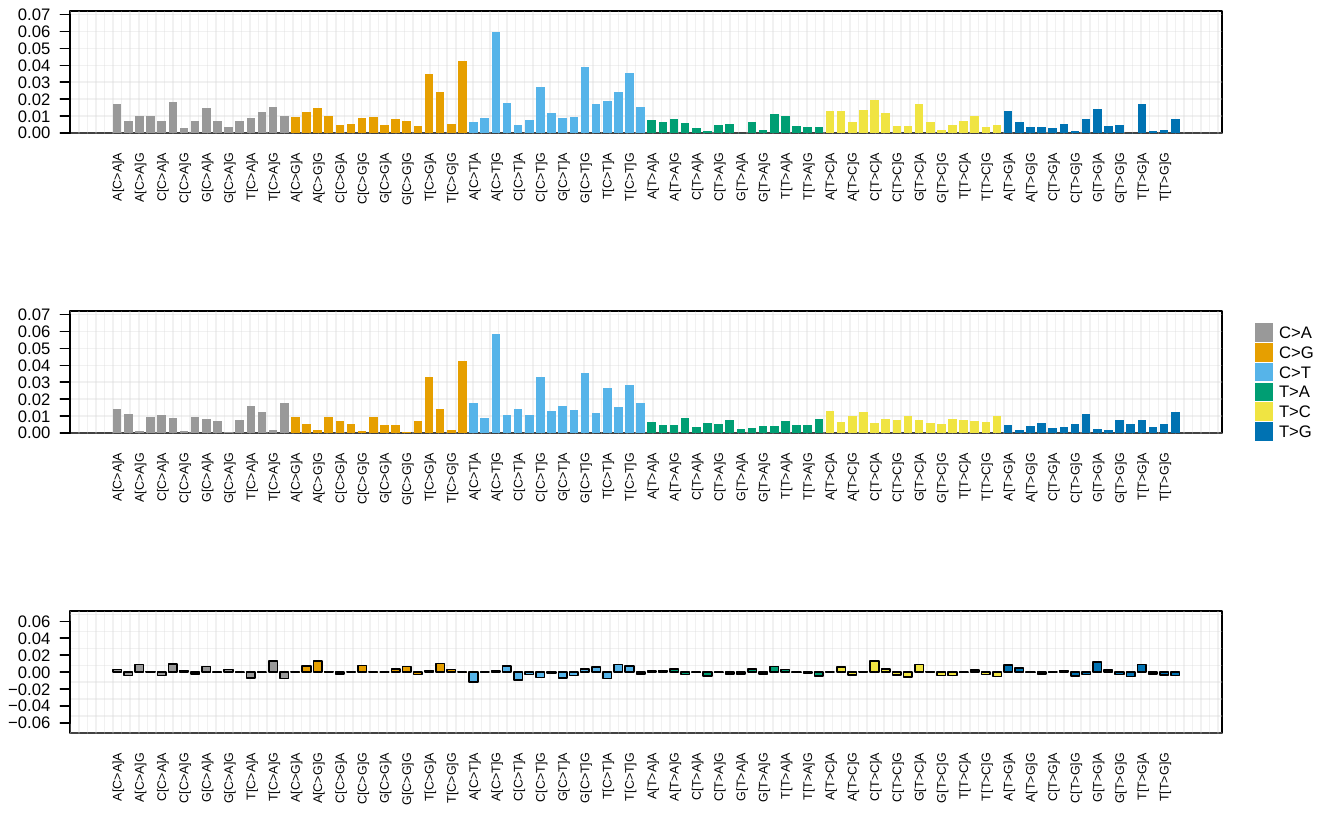
 **Figure S1: Somatic-mutation signature weights for one Signature 3 tumor.**  This tumor had a large proportion of C>T mutations, which are representative of Signature 3.

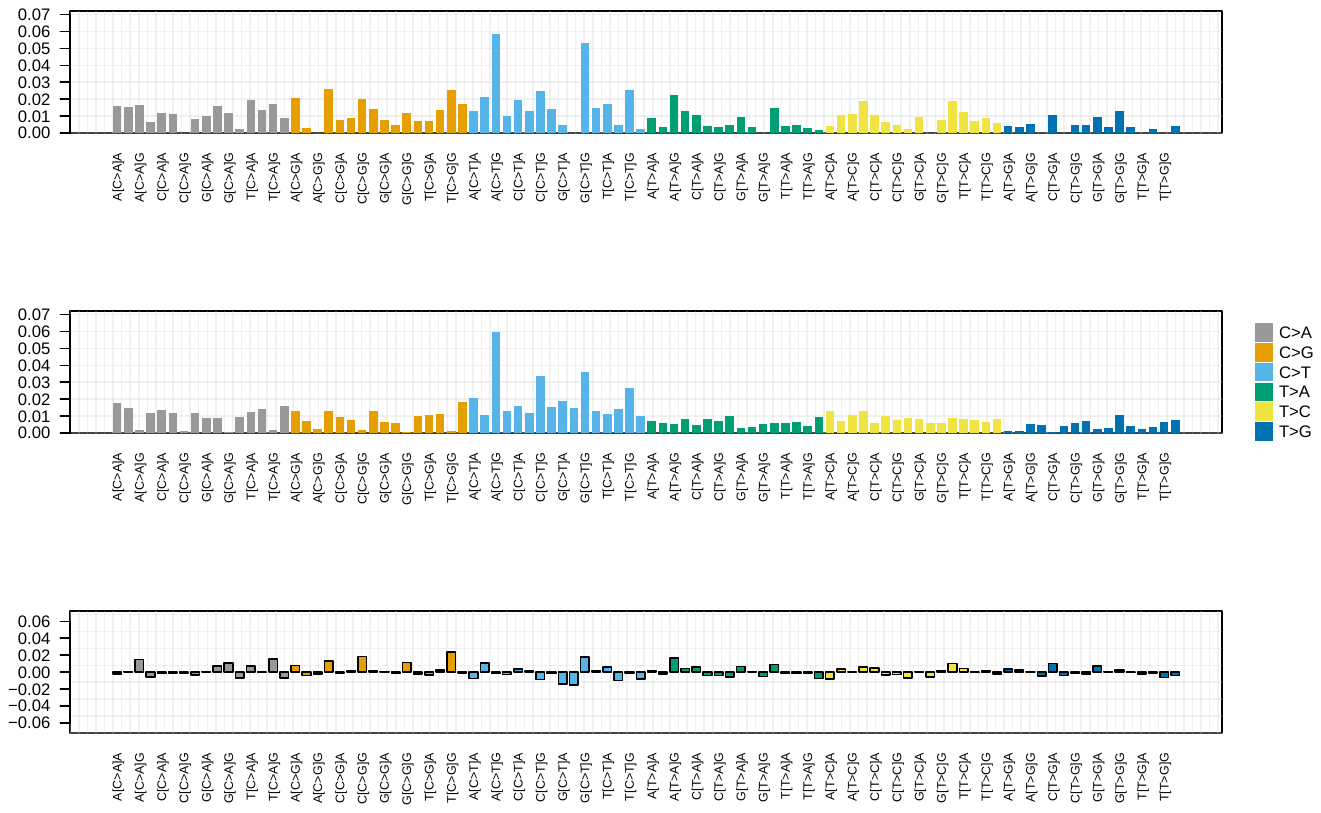
 **Figure S2: Somatic-mutation signature weights for a second Signature 3 tumor.**  This tumor had a large proportion of C>T mutations, which are representative of Signature 3.

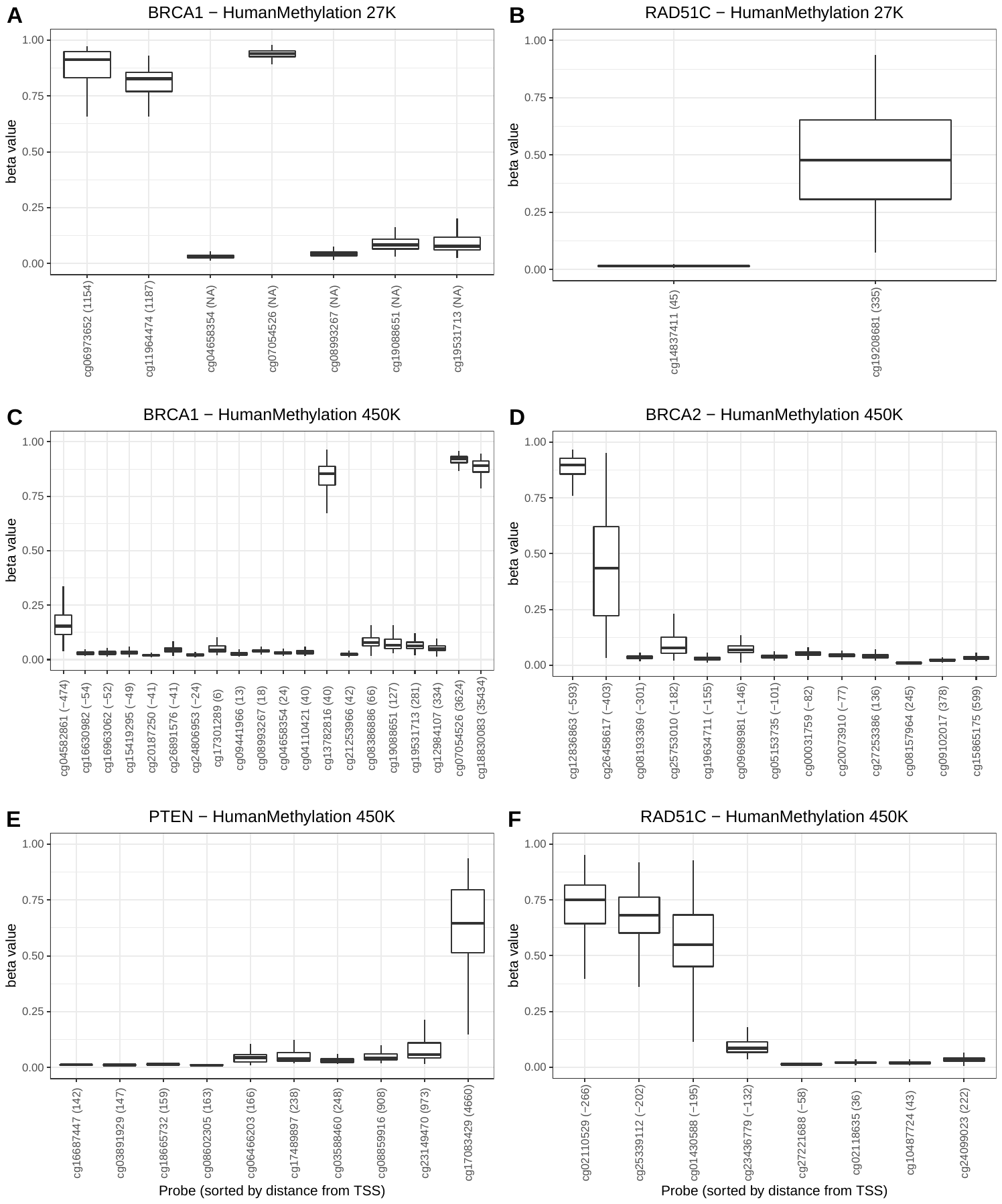
 **Figure S3: Probe-level summarization of DNA methylation probes.**  We extracted probe-level methylation (beta) values for all available breast-cancer samples in TCGA and plotted them relative to the transcription start site of each gene. These graphs illustrate beta values for four genes (*BRCA1*, *BRCA2*, *PTEN*, and *RAD51C*) and two microarray platforms (Illumina HumanMethylation 27K and 450K). Values in parenthesis indicate distance from the transcription start site (TSS). TSS distances marked as “NA” were unavailable. The 27K arrays have fewer probes per gene. In general, probes near the TSS exhibited relatively low methylation levels for these genes, whereas probes further from the TSS were more highly methylated. These observations are consistent with the assumption that most genes would be “on” by default. Some exceptions to this pattern are apparent (for example, cg13782816 on panel C); these exceptions may be caused by mismapped probes, cross hybridization, or misannotations. We calculated gene-level values as the median across all probes that were within 300 nucleotides of the TSS.

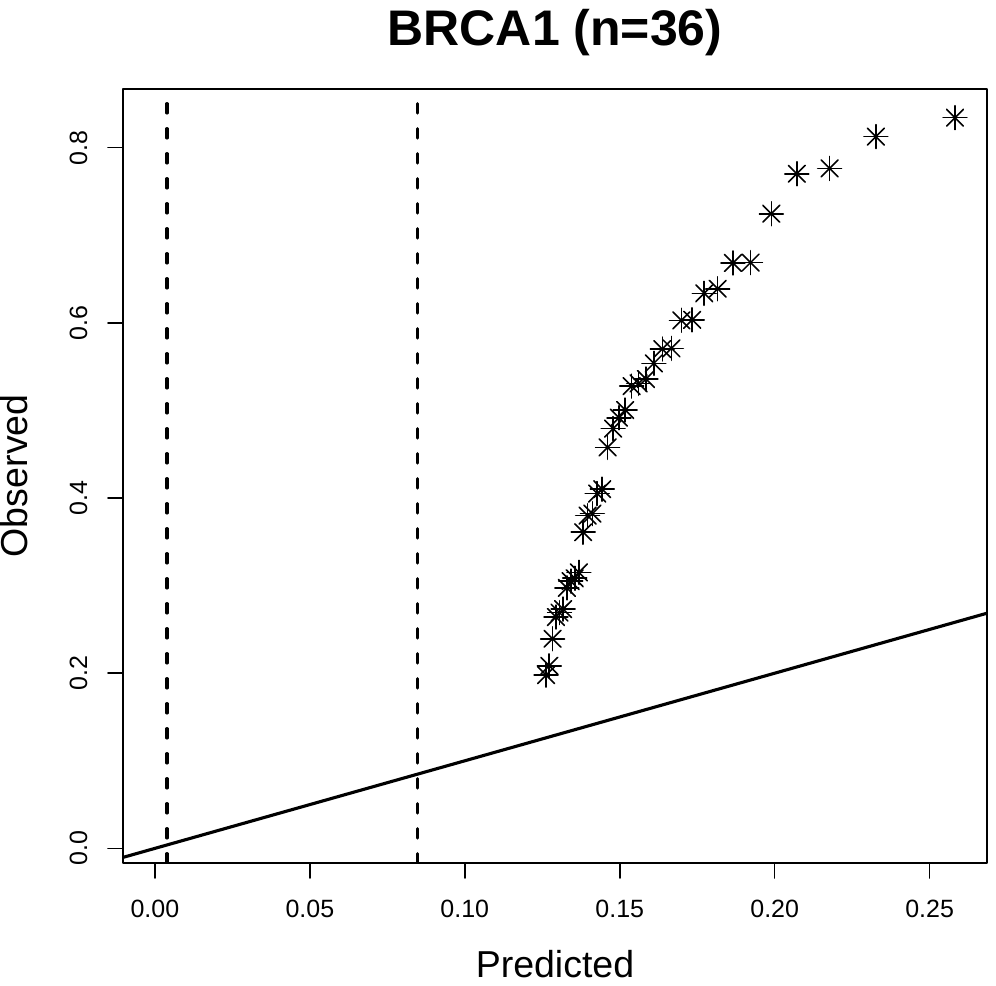
 **Figure S4: Fit of outlier-detection model to DNA methylation data for BRCA1.**  We used an outlier-detection methodology to estimate which tumors were hypermethylated for a given gene. This scatter plot illustrates the model fit for BRCA1. Asterisks represent tumors considered to be outliers.

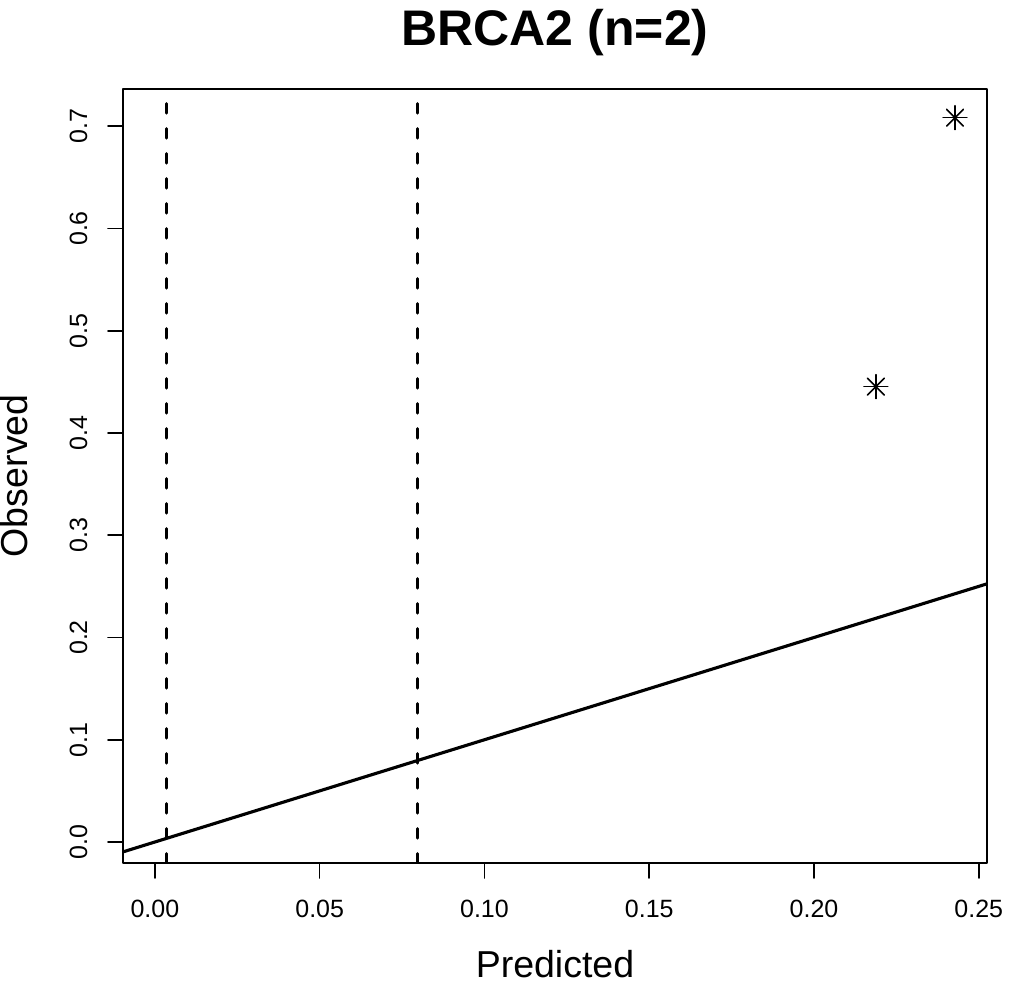
 **Figure S5: Fit of outlier-detection model to DNA methylation data for BRCA2.**  We used an outlier-detection methodology to estimate which tumors were hypermethylated for a given gene. This scatter plot illustrates the model fit for BRCA2. Asterisks represent tumors considered to be outliers.

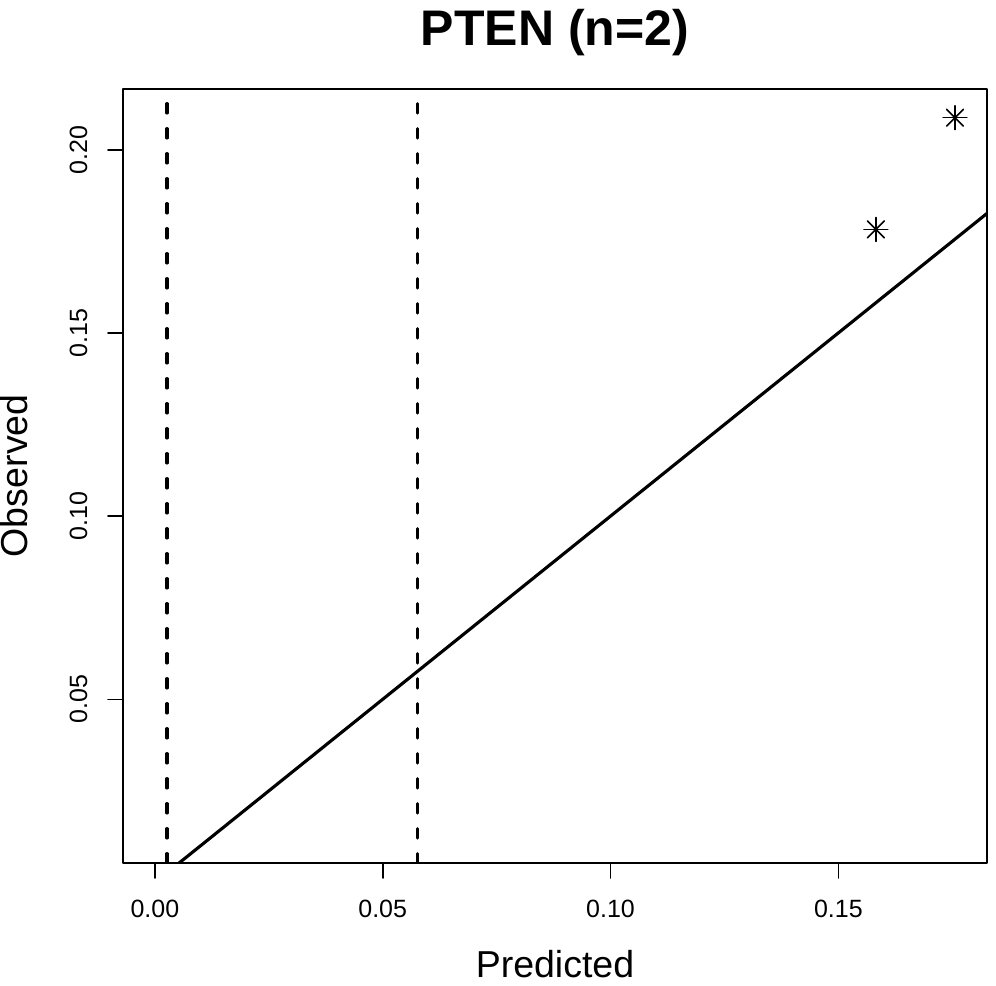
 **Figure S6: Fit of outlier-detection model to DNA methylation data for PTEN.**  We used an outlier-detection methodology to estimate which tumors were hypermethylated for a given gene. This scatter plot illustrates the model fit for PTEN. Asterisks represent tumors considered to be outliers.

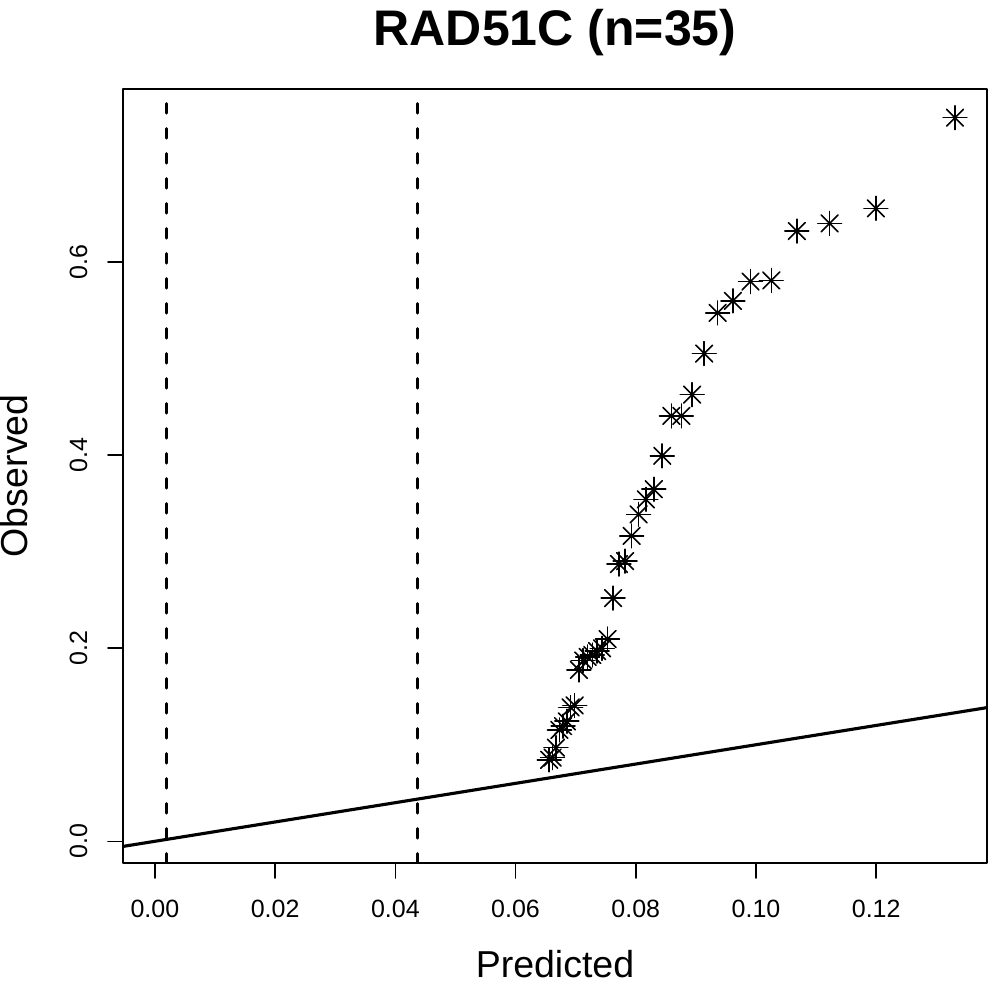
 **Figure S7: Fit of outlier-detection model to DNA methylation data for RAD51C.**  We used an outlier-detection methodology to estimate which tumors were hypermethylated for a given gene. This scatter plot illustrates the model fit for RAD51C. Asterisks represent tumors considered to be outliers.

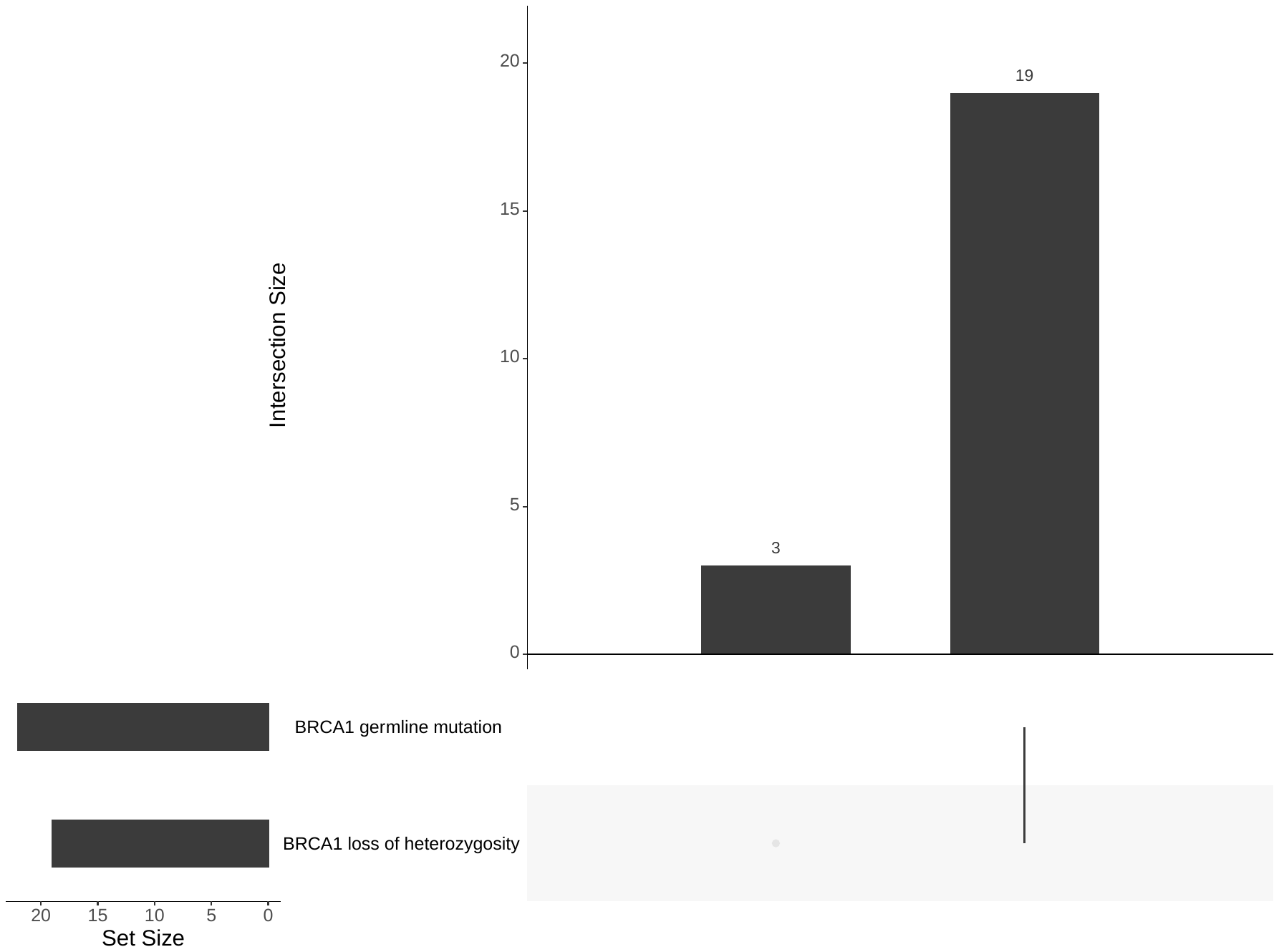
 **Figure S8: Intersection between germline-mutation status and loss of heterozygosity for *BRCA1*.**  A total of 22 patients carried a germline mutation in *BRCA1*. We detected loss-of-heterozygosity events in tumors for all but 3 of these patients. Data are only shown for patients for whom we had both types of data.

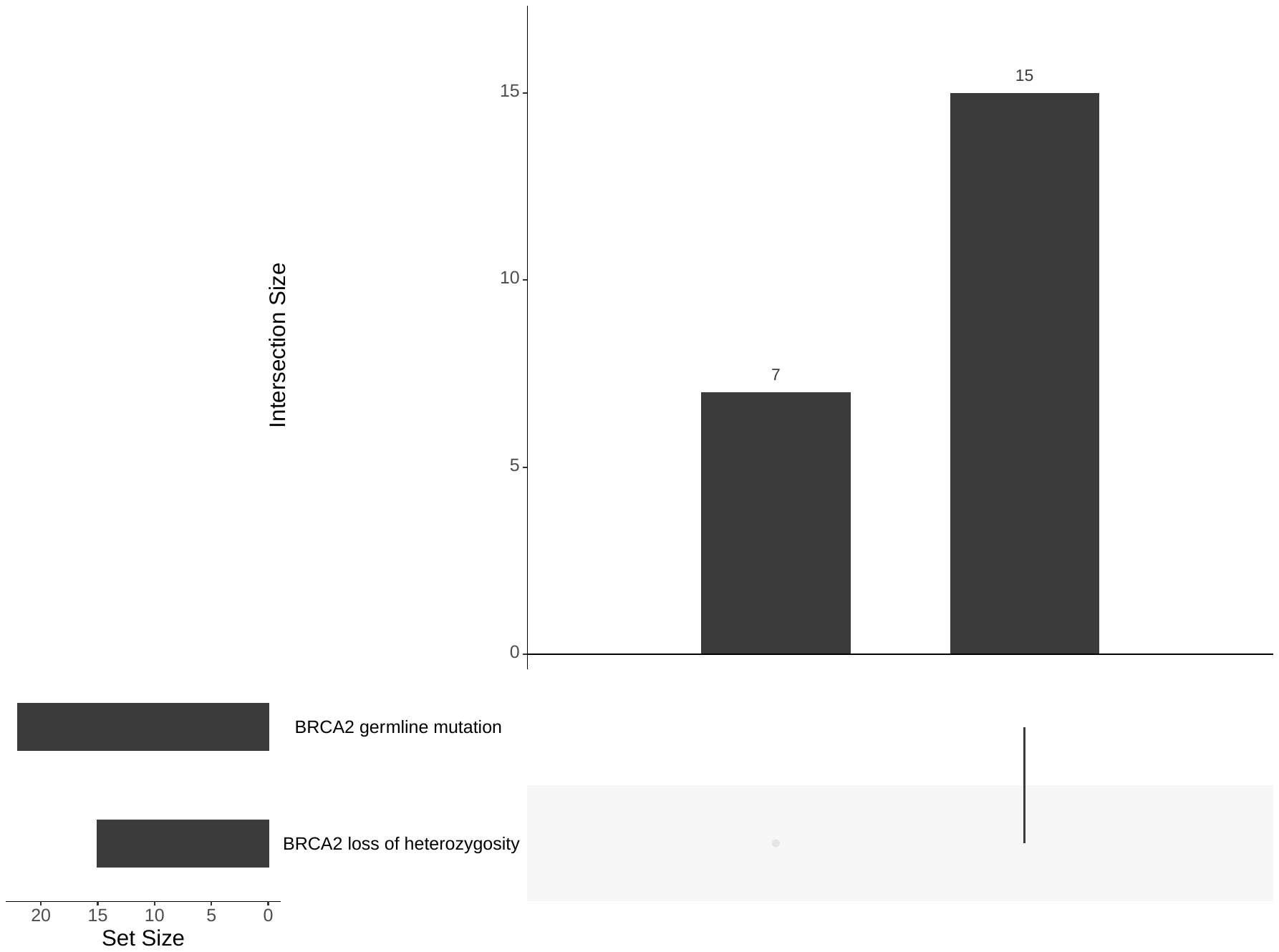
 **Figure S9: Intersection between germline-mutation status and loss of heterozygosity for *BRCA2*.**  A total of 22 patients carried a germline mutation in *BRCA2*. We detected loss-of-heterozygosity events in tumors from all but 7 of these patients. Data are only shown for patients for whom we had both types of data.

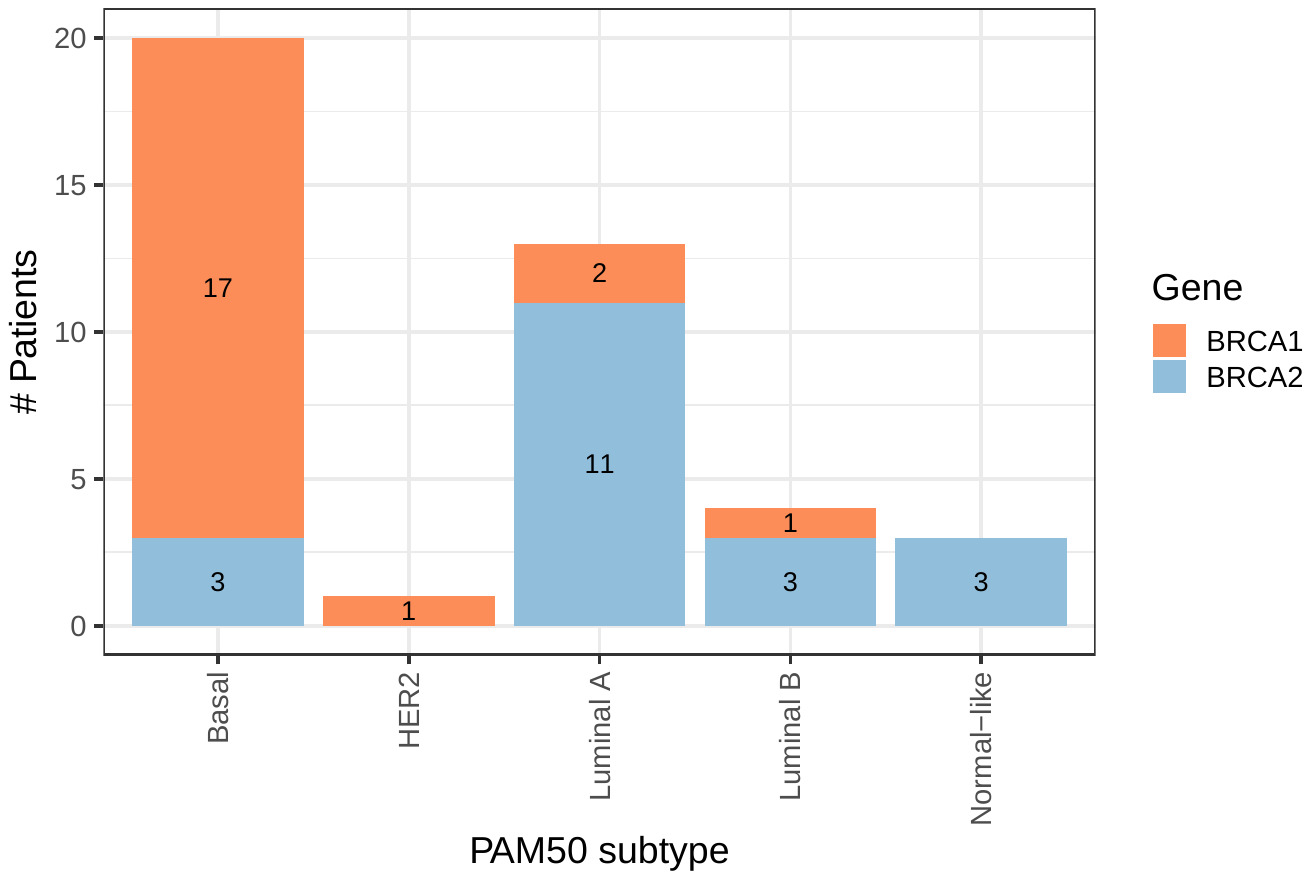
 **Figure S10: Overlap between *BRCA1*/*BRCA2* germline-mutation status and PAM50 subtype.**  Gene-expression subtypes were unavailable for some patients; One BRCA1 carrier is not represented in this figure; we could not assign a gene-expression subtype to this individual due to missing data.

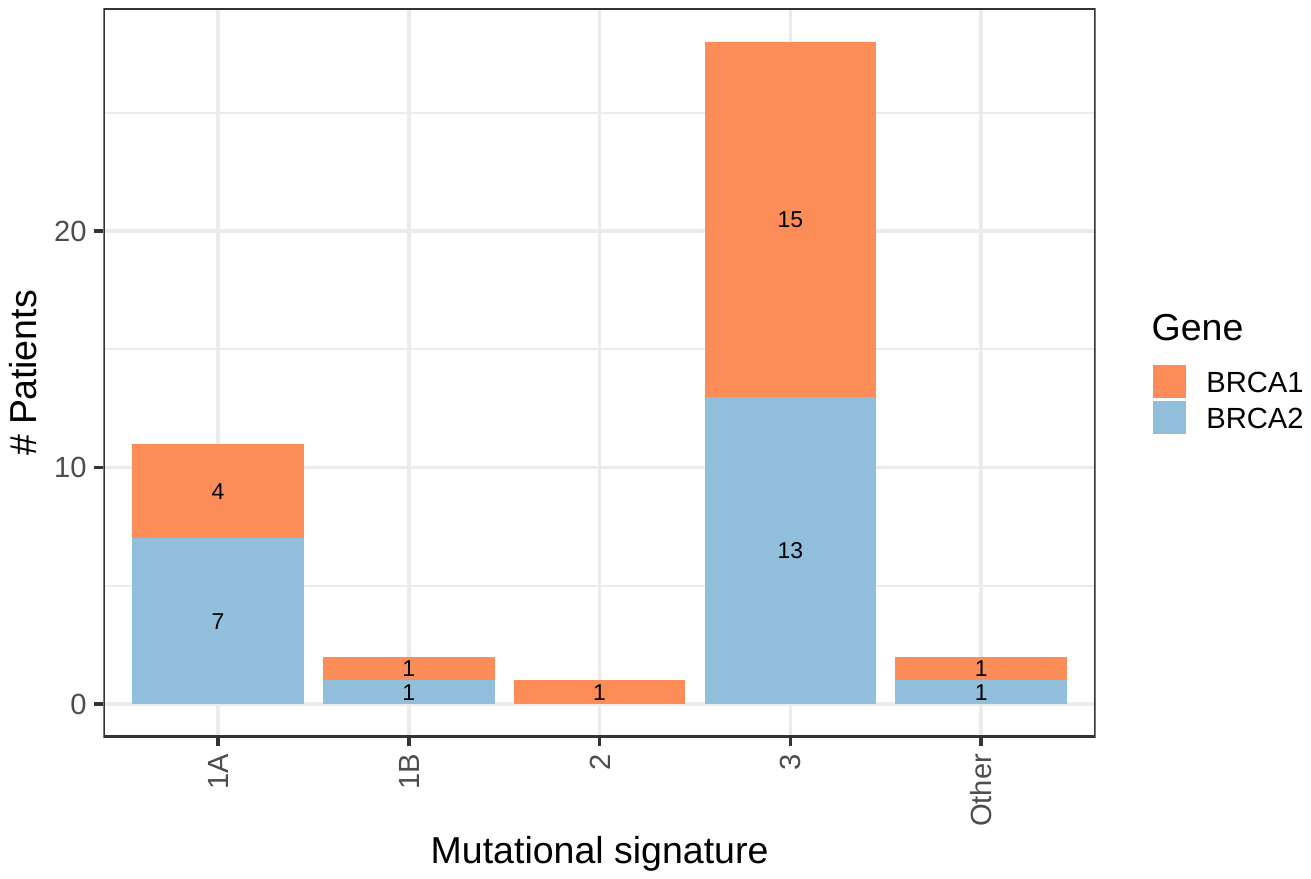
 **Figure S11: Overlap between *BRCA1*/*BRCA2* germline-mutation status and primary somatic-mutation signature.**  This graph represents patients for whom we had both germline- and somatic-mutation data.

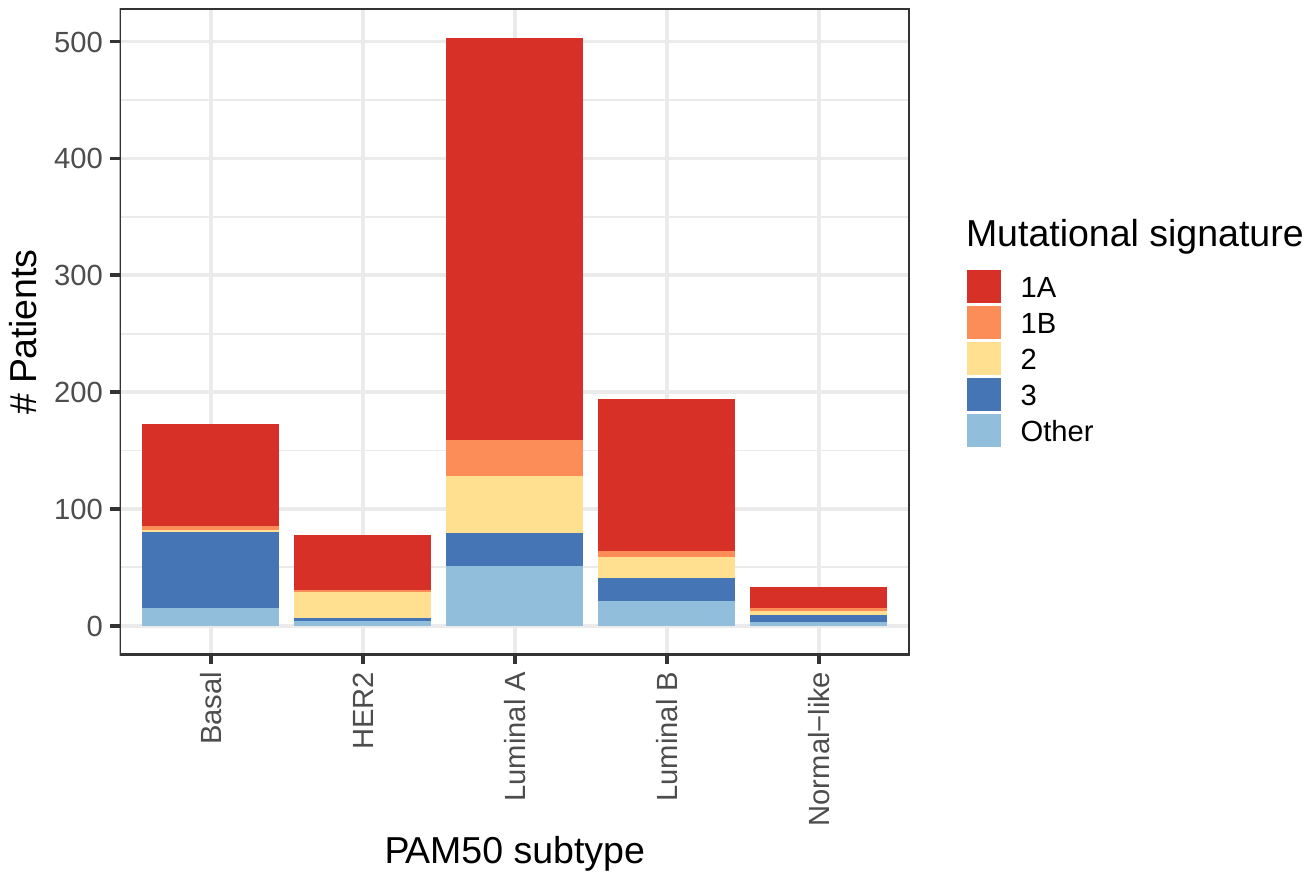
 **Figure S12: Overlap between PAM50 subtype and primary somatic-mutation signature.**  This graph represents patients for whom we could evaluate the status of both PAM50 subtype and somatic-mutation signatures.

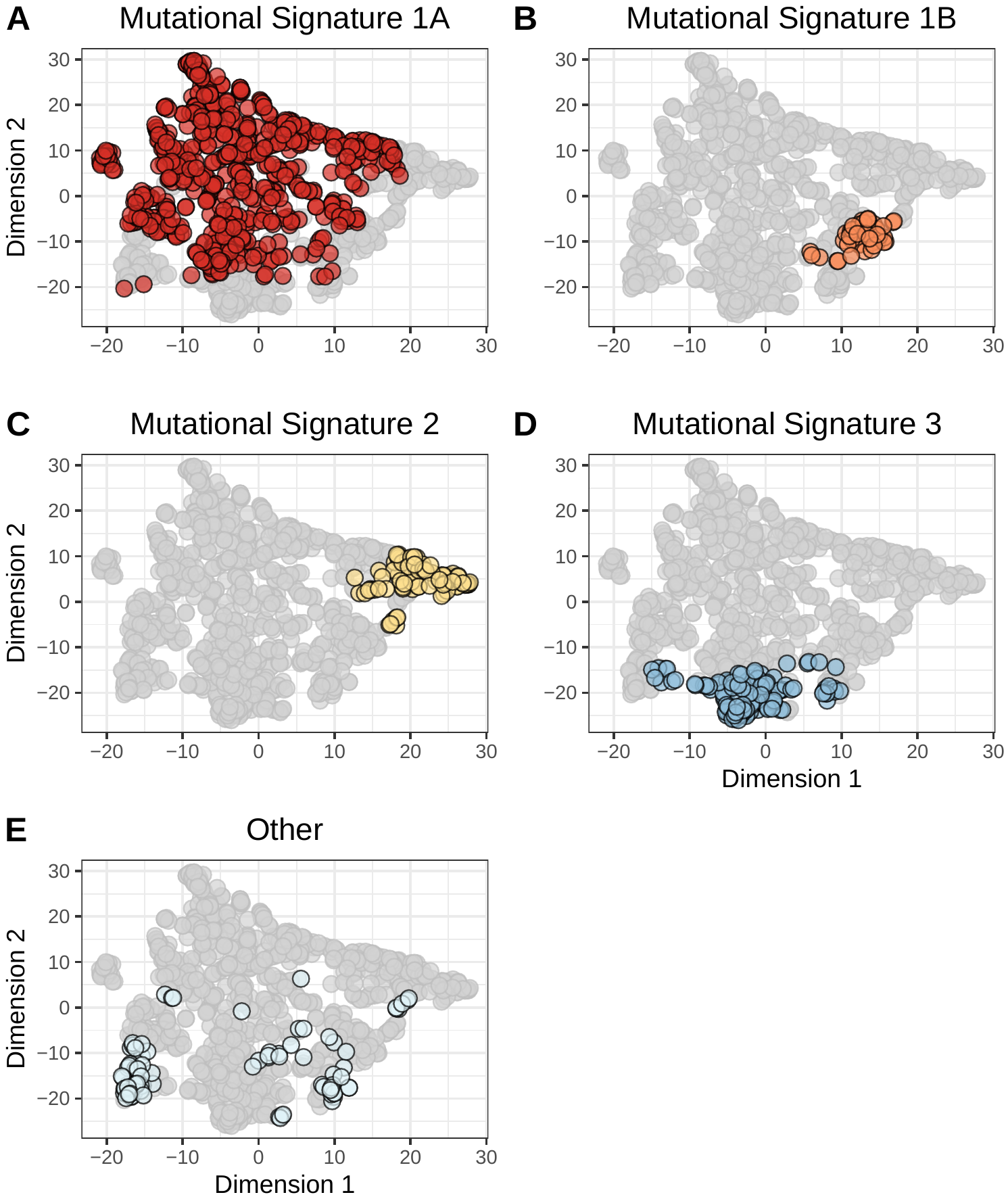
 **Figure S13: Two-dimensional representation of somatic-mutation signatures using the *t*-SNE method.**  We summarized each tumor based on their somatic-mutation signatures, which represent overall mutational patterns in a trinucleotide context. We used the *t*-distributed Stochastic Neighbor Embedding (*t*-SNE) method to reduce the data to two dimensions. Each point represents a single tumor, overlaid with colors that represent the tumor’s primary somatic-mutation signature. Mutational Signature 1A (A) was the most prevalent; these tumors were widely dispersed across the signature landscape. Signatures 1B (B), 2 (C), and 3 (D) were relatively small and formed cohesive clusters. The remaining 23 clusters were rare individually and were dispersed broadly.

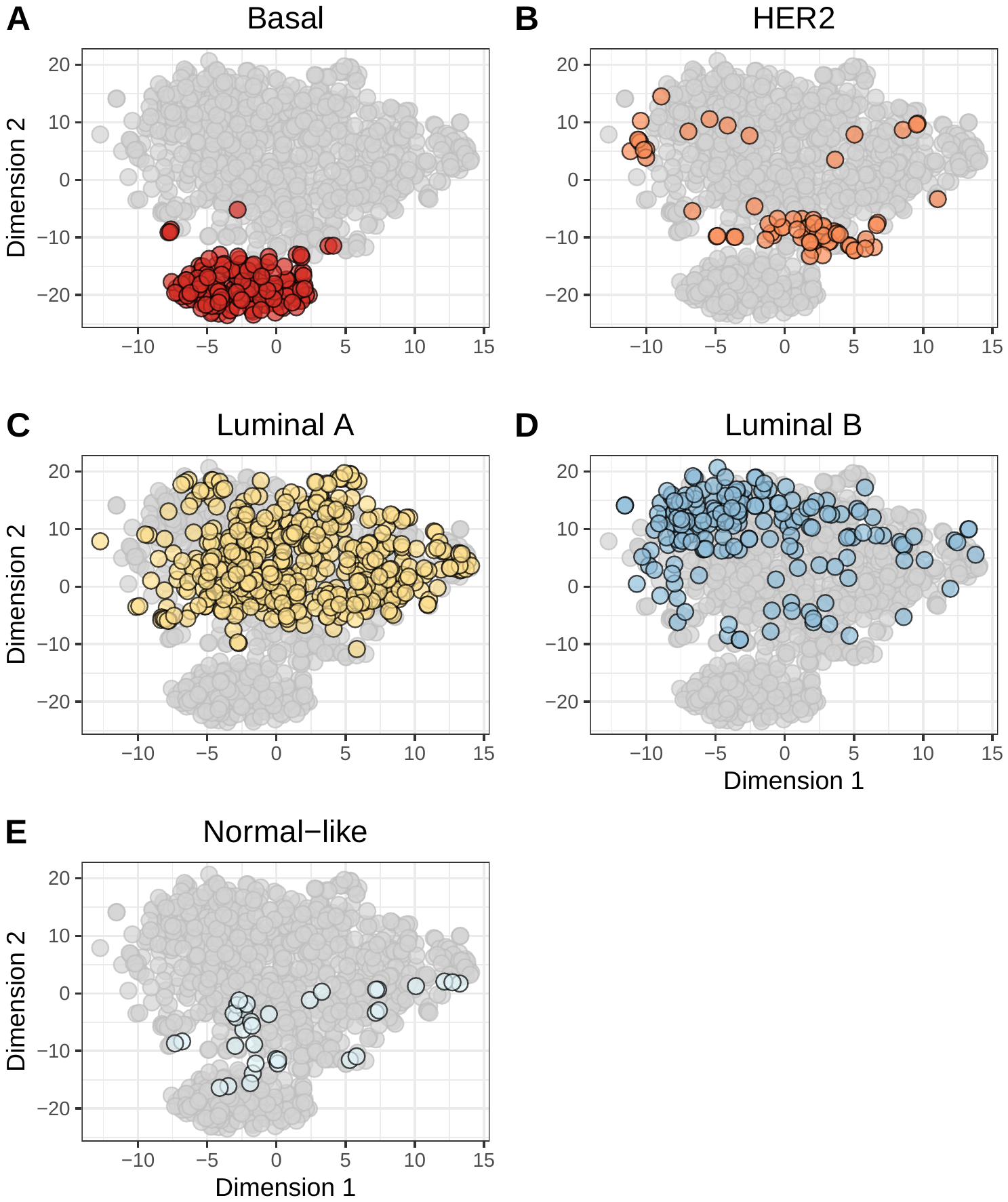
 **Figure S14: Two-dimensional representation of gene-expression levels using the *t*-SNE method.**  We used the *t*-distributed Stochastic Neighbor Embedding (*t*-SNE) method to reduce the gene-expression profiles to two dimensions. Each point represents a single tumor, overlaid with colors that represent the tumor’s primary PAM50 subtype. Generally, the PAM50 subtypes clustered cohesively, but there were exceptions. For example, some Basal-like tumors (A) exhbited expression patterns that differed considerably from the remaining Basal-like tumors. The normal-like tumors (E) showed the most variability in expression. This graph represents patients for whom we could identify a PAM50 subtype.

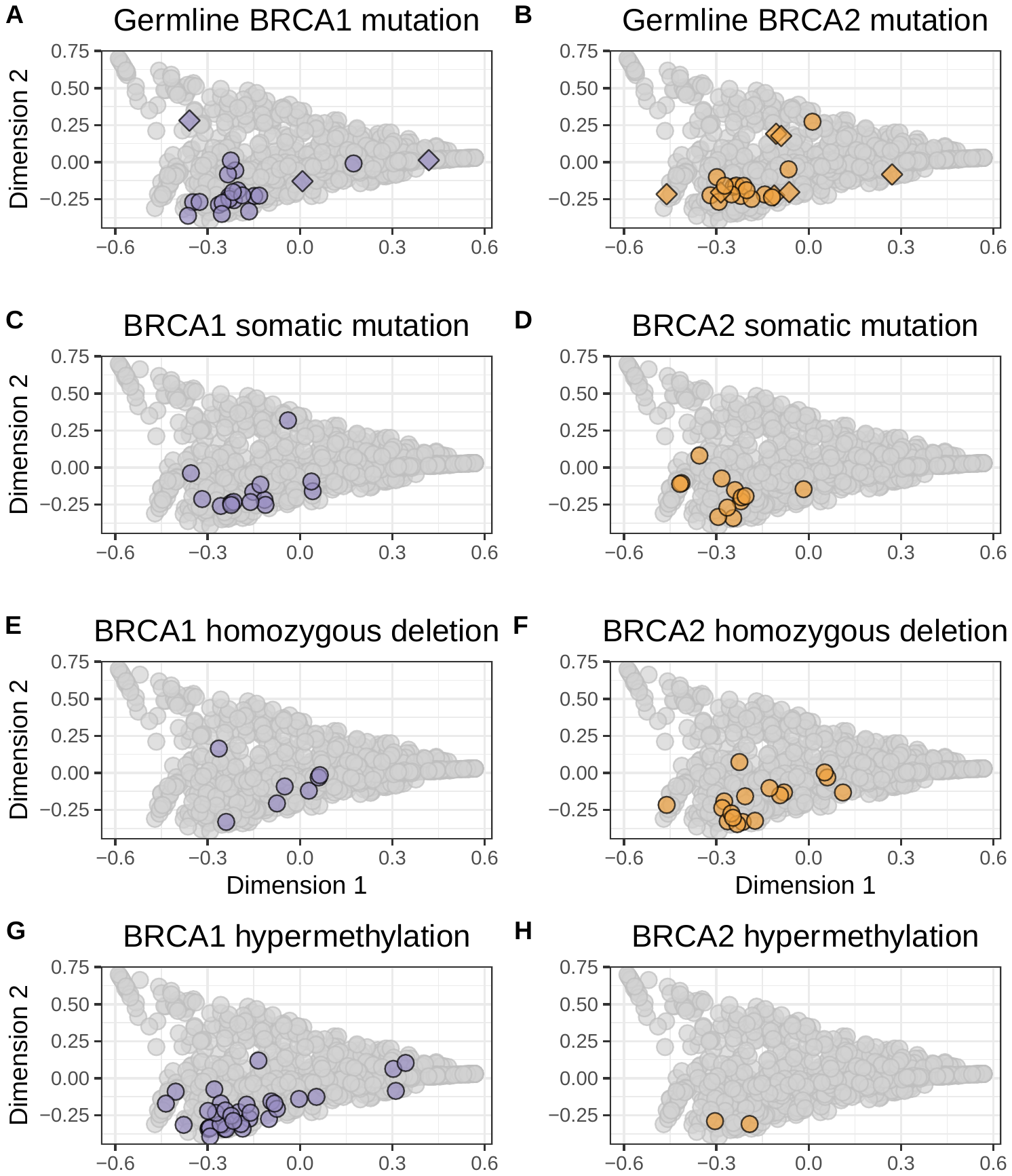
 **Figure S15: *BRCA1* and *BRCA2* aberrations on the somatic-mutation signature landscape using multidimensional scaling.**  Using the same two-dimensional representation of mutational signatures shown in Figure 2, this plot indicates which patients had germline mutations (A, B), somatic mutations (C, D), homozygous deletions (E, F), or hypermethylation events (G, H) in *BRCA1* and *BRCA2*, respectively. Largely, these tumors had similar somatic-mutation signatures. Diamond shapes indicate patients for whom *no* loss-of-heterozygosity was observed. Data are shown for all patients, even those for whom we did not have all types of data.

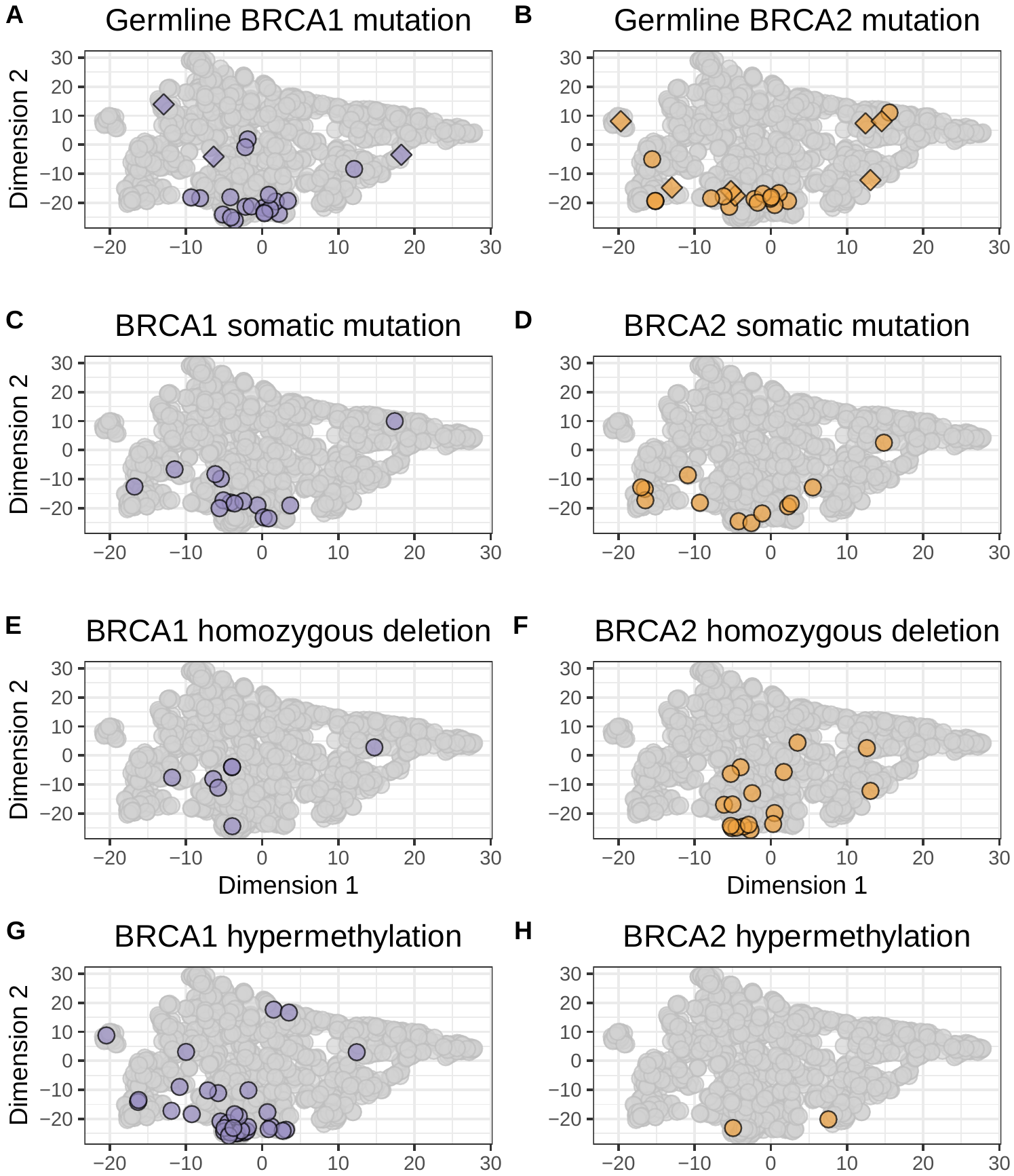
 **Figure S16: *BRCA1* and *BRCA2* aberrations on the somatic-mutation signature landscape using the *t*-SNE method.**  Using the same two-dimensional representation of mutational signatures shown in Figure S13, this plot indicates which patients had germline mutations (A, B), somatic mutations (C, D), homozygous deletions (E, F), or hypermethylation events (G, H) in *BRCA1* and *BRCA2, respectively. Largely, these tumors had similar somatic-mutation signatures. Diamond shapes indicate patients for whom* no* loss-of-heterozygosity was observed. Data are shown for all patients, even those for whom we did not have all types of data.

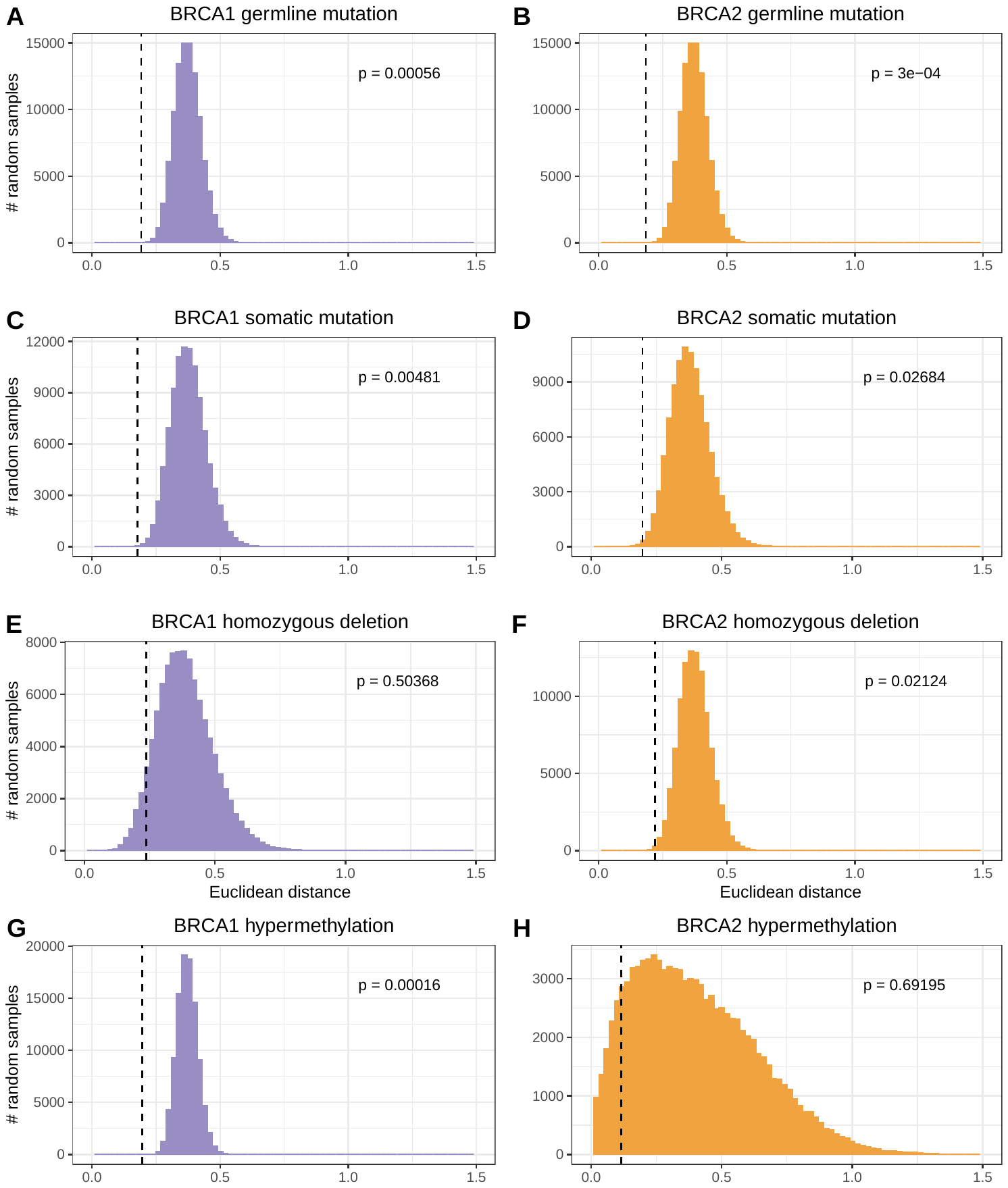
 **Figure S17: Euclidean distances for randomly selected patients compared to actual distances within *BRCA1*/*BRCA2* patient groups based on somatic-mutation signatures.**  We calculated the Euclidean distance between each pair of individuals who had germline mutations (A, B), somatic mutations (C, D), homozygous deletions (E, F), or hypermethylation events (G, H) in *BRCA1* or *BRCA2*; the medians of these distances are illustrated using vertical, dashed lines. We then randomized the patient identifiers and calculated pairwise distances for the same number of randomly selected patients, which resulted in an empirical null distribution. We calculated p-values by comparing the actual distances against the randomized distances and then adjusted for multiple tests using Holm’s method.

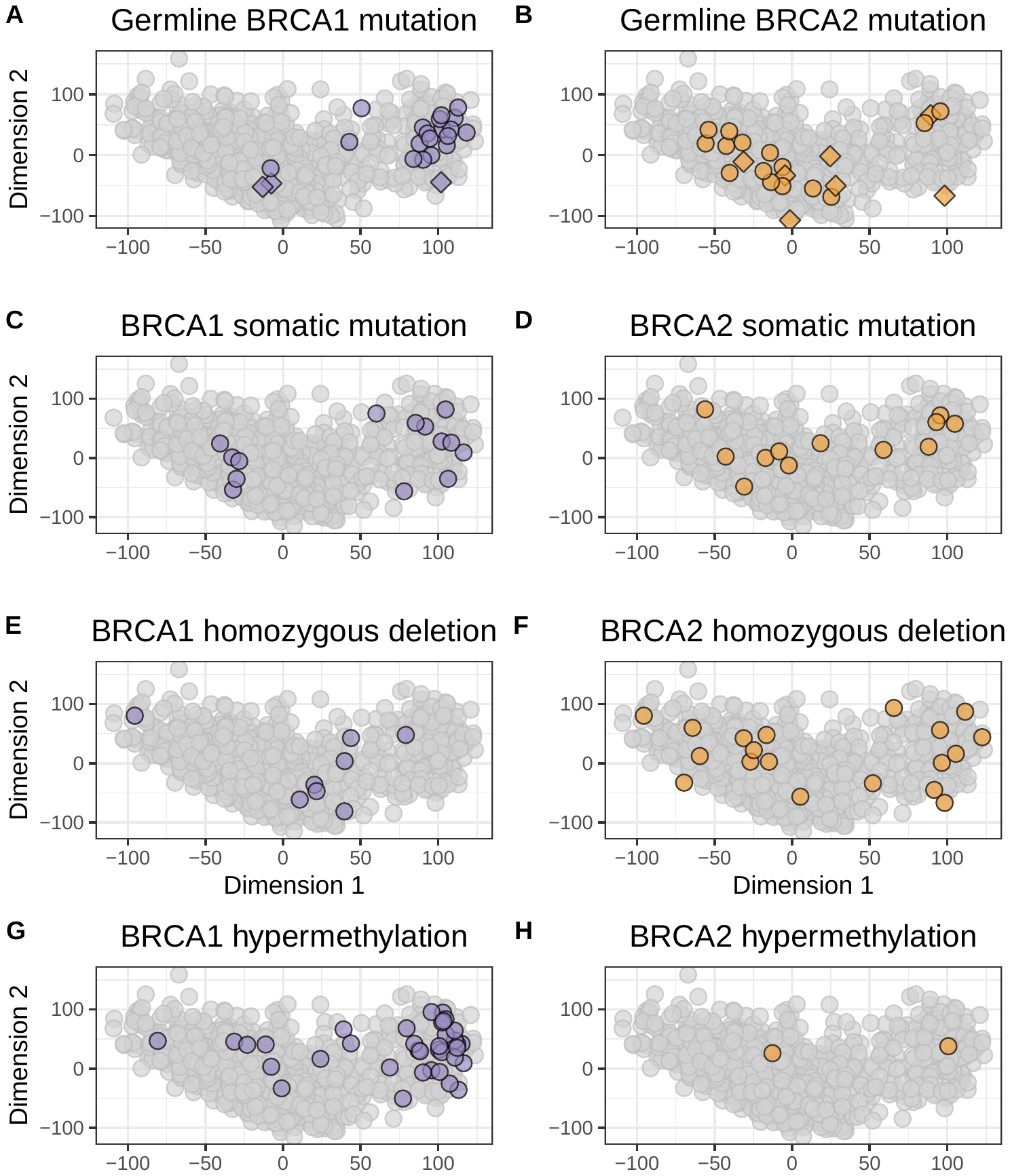
 **Figure S18: *BRCA1* and *BRCA2* aberrations on the gene-expression landscape using multidimensional scaling.**  Using the same two-dimensional representation of gene-expression profiles shown in Figure 3, this plot indicates which patients had germline mutations (A, B), somatic mutations (C, D), homozygous deletions (E, F), or hypermethylation events (G, H) in *BRCA1* and *BRCA2*, respectively. Many of these tumors overlapped with the Basal-like subtype, but other tumors were dispersed broadly across the gene-expression landscape. Diamond shapes indicate patients for whom *no* loss-of-heterozygosity was observed. Data are shown for all patients, even those for whom we did not have all types of data.

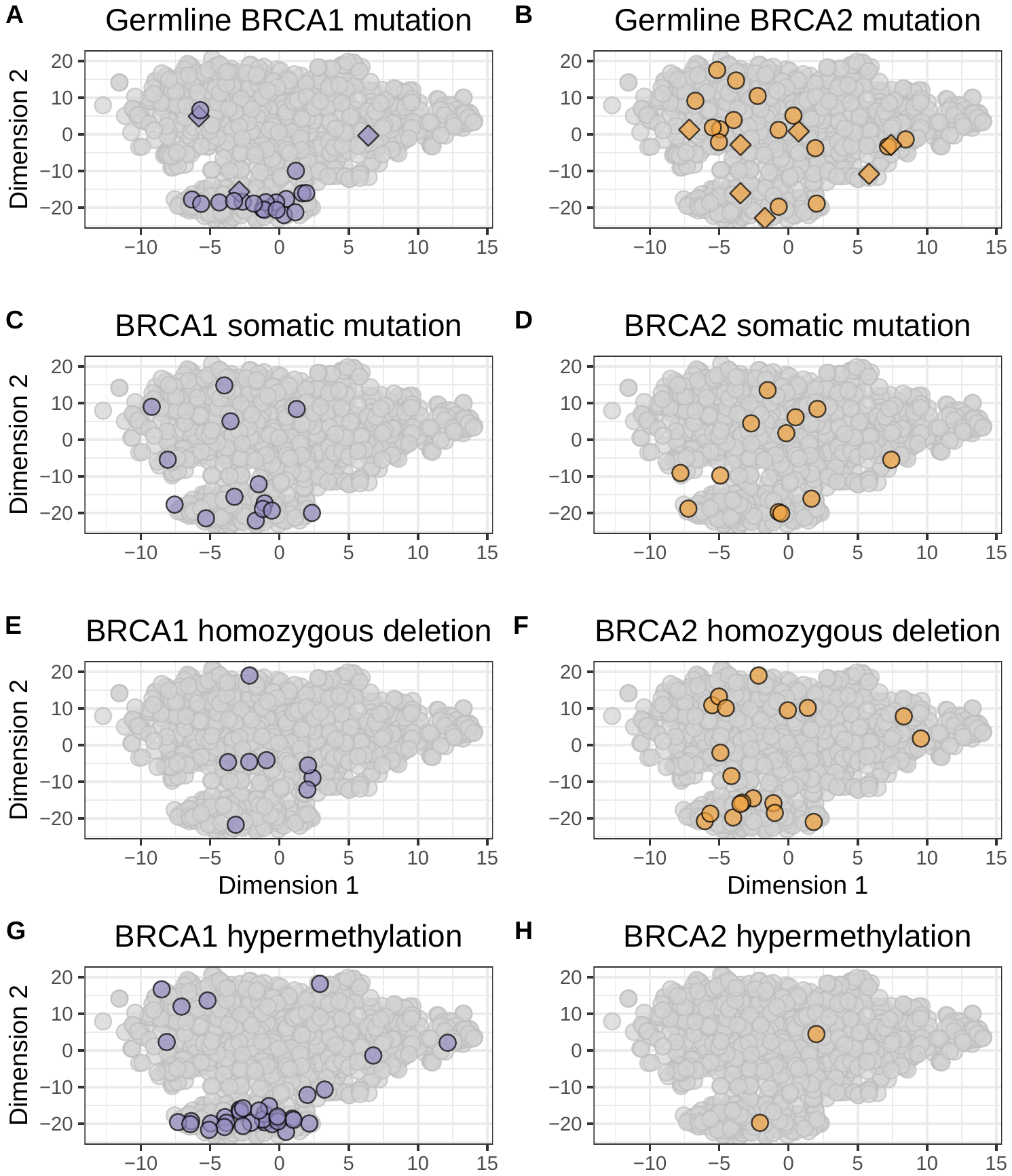
 **Figure S19: *BRCA1* and *BRCA2* aberrations on the gene-expression landscape using the *t*-SNE method.**  Using the same two-dimensional representation of gene-expression profiles shown in Figure S14, this plot indicates which patients had germline mutations (A, B), somatic mutations (C, D), homozygous deletions (E, F), or hypermethylation events (G, H) in *BRCA1* and *BRCA2*, respectively. Many of these tumors overlapped with the Basal-like subtype, but other tumors were dispersed broadly across the gene-expression landscape. Diamond shapes indicate patients for whom *no* loss-of-heterozygosity was observed. Data are shown for all patients, even those for whom we did not have all types of data.

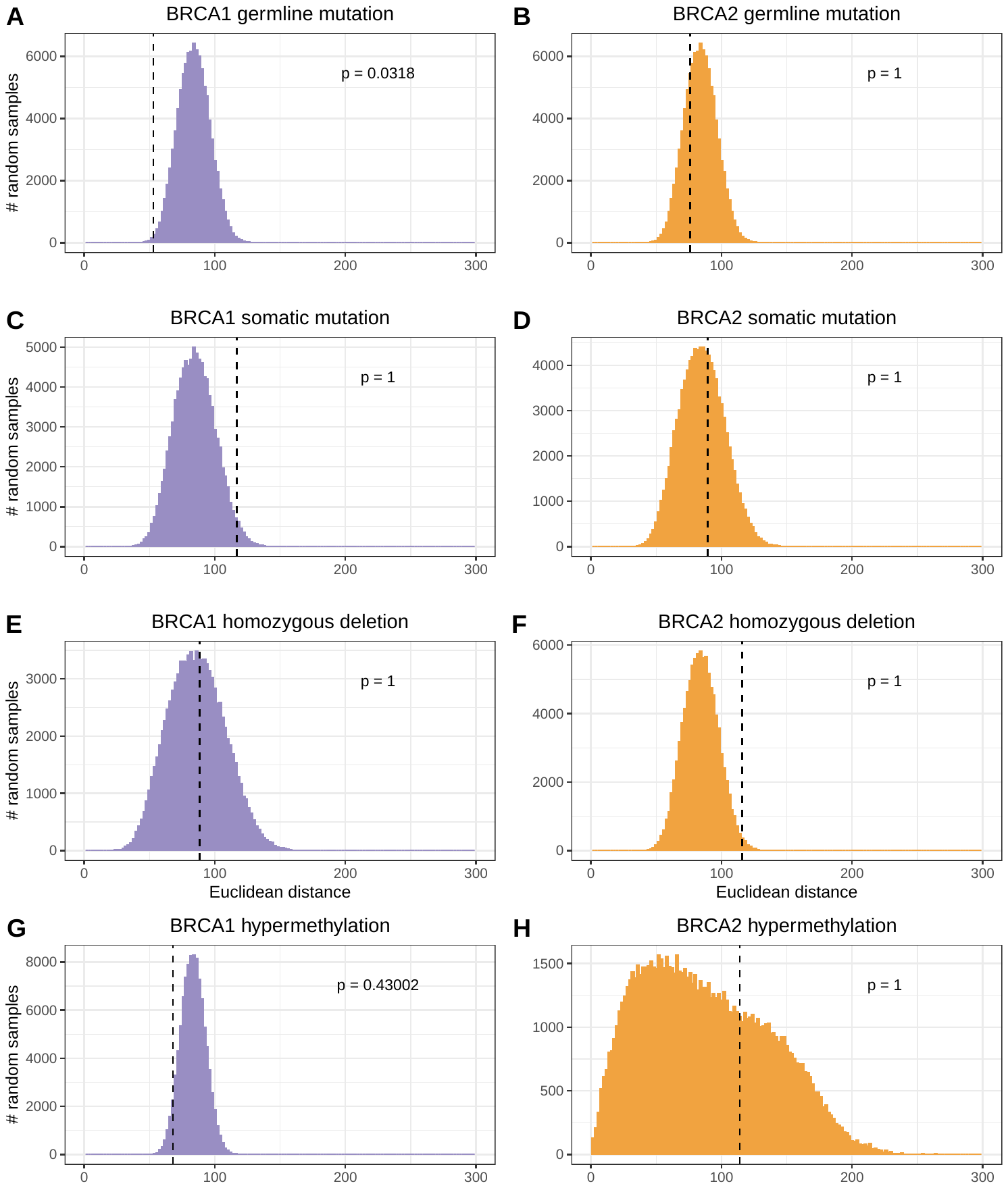
 **Figure S20: Euclidean distances for randomly selected patients compared to actual distances within *BRCA1*/*BRCA2* patient groups based on gene-expression profiles.**  We calculated the Euclidean distance between each pair of individuals who had germline mutations (A, B), somatic mutations (C, D), homozygous deletions (E, F), or hypermethylation events (G, H) in *BRCA1* or *BRCA2*; the medians of these distances are illustrated using vertical, dashed lines. We then randomized the patient identifiers and calculated pairwise distances for the same number of randomly selected patients, which resulted in an empirical null distribution. We calculated p-values by comparing the actual distances against the randomized distances and then adjusted for multiple tests using Holm’s method.

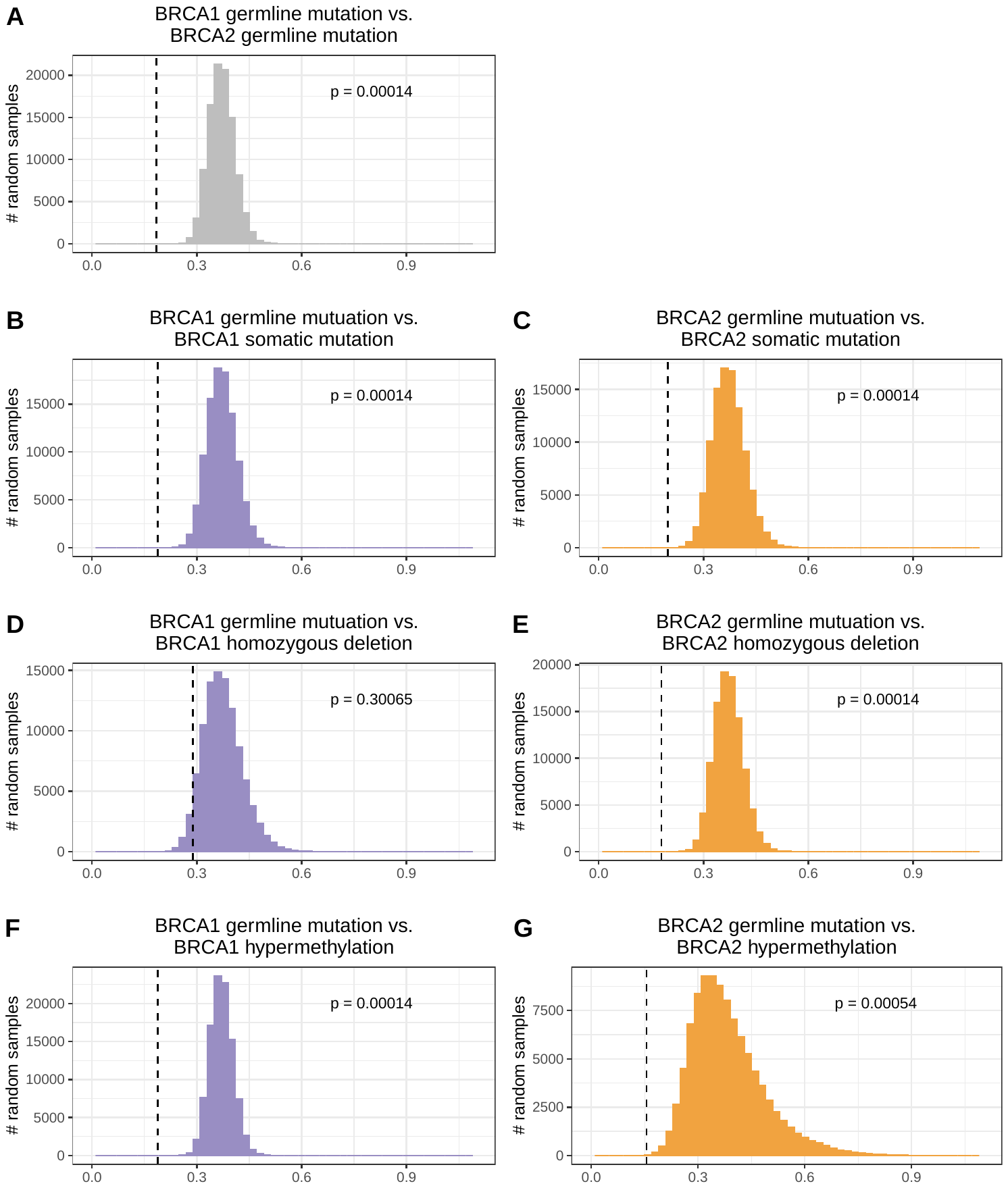
 **Figure S21: Somatic-mutation signature-based Euclidean distances for randomly selected patient pairs compared to actual distances between patient pairs for individuals with *BRCA1*/*BRCA2* aberrations.**  We identified patients who had a germline mutation in *BRCA1* or *BRCA2* and compared them against each other (A), those with a somatic mutation in the same gene (B-C), those with a homozygous deletion in the same gene (D-E) and those with DNA hypermethylation of the same gene (F-G). We calculated the Euclidean distance between each pair of individuals in these groups; these distances are illustrated using vertical, dashed lines. We then randomized the patient identifiers and calculated pairwise distances for groups of randomly selected patients, which resulted in an empirical null distribution. We calculated p-values by comparing the actual distances against the randomized distances.

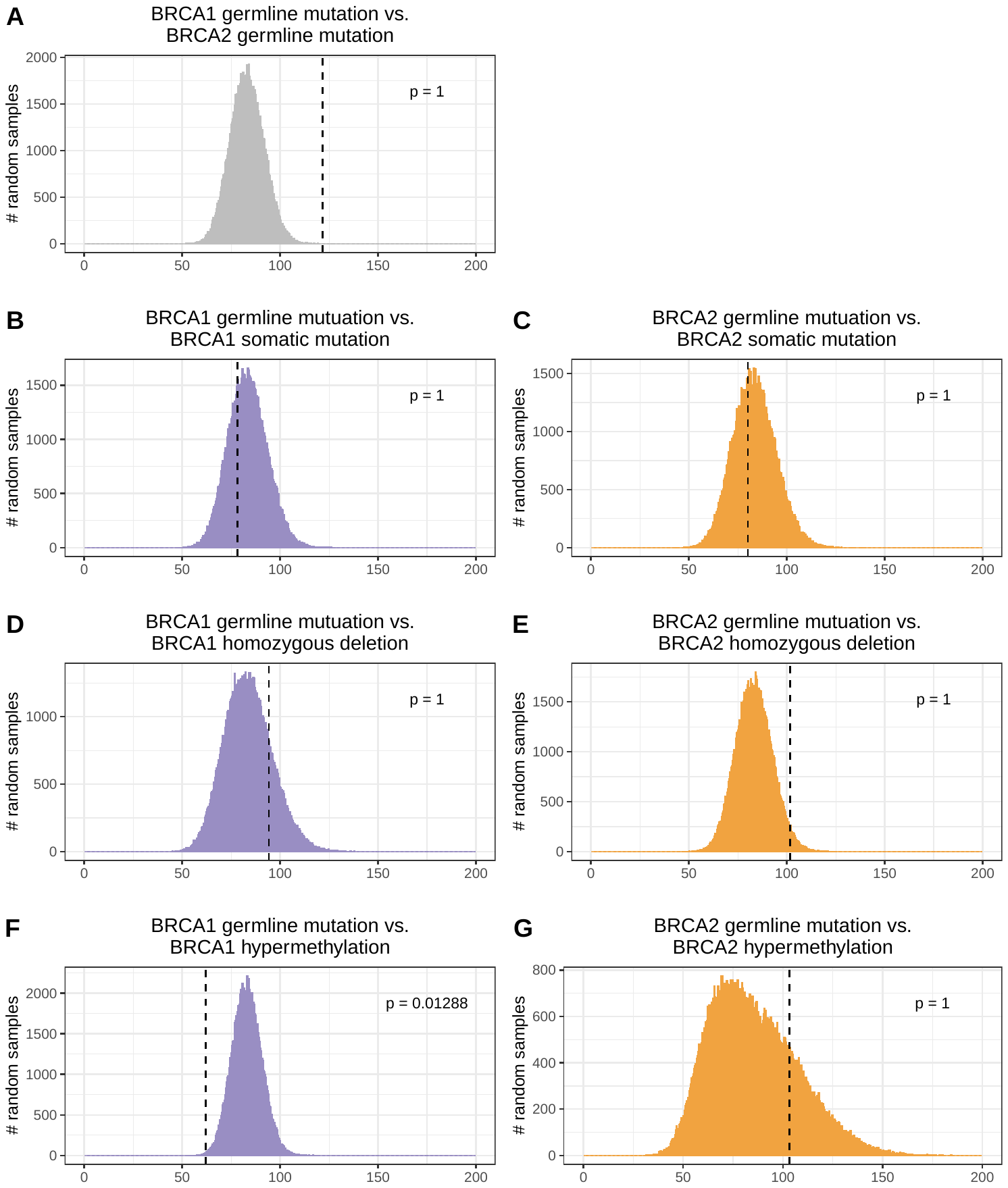
 **Figure S22: Gene-expression based Euclidean distances for randomly selected patient pairs compared to actual distances between patient pairs for individuals with *BRCA1*/*BRCA2* aberrations.**  We identified patients who had a germline mutation in *BRCA1* or *BRCA2* and compared them against each other (A), those with a somatic mutation in the same gene (B-C), those with a homozygous deletion in the same gene (D-E) and those with DNA hypermethylation of the same gene (F-G). We calculated the Euclidean distance between each pair of individuals in these groups; these distances are illustrated using vertical, dashed lines. We then randomized the patient identifiers and calculated pairwise distances for groups of randomly selected patients, which resulted in an empirical null distribution. We calculated p-values by comparing the actual distances against the randomized distances.

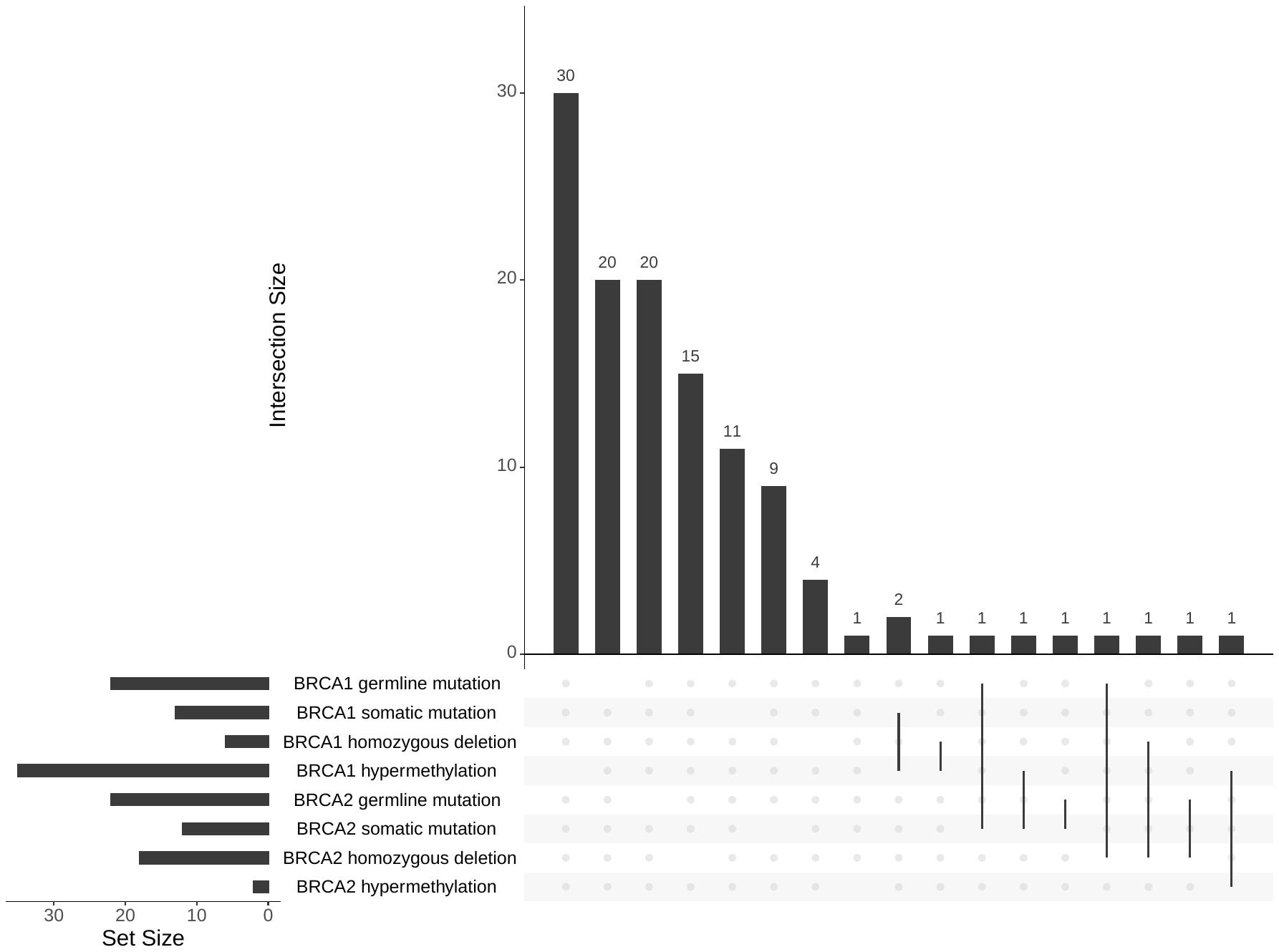
 **Figure S23: Intersection between different types of molecular aberration in *BRCA1* and *BRCA2*.**  This graph indicates how many patients had each type of molecular aberration and the level of overlap among these aberrations within a given patient. In most cases, these aberrations were mutually exclusive from each other; however, some overlap did occur. For example, one patient had a somatic mutation in *BRCA1* and hypermethylation of the same gene. This graph only depicts patients for whom all four types of molecular data were available.

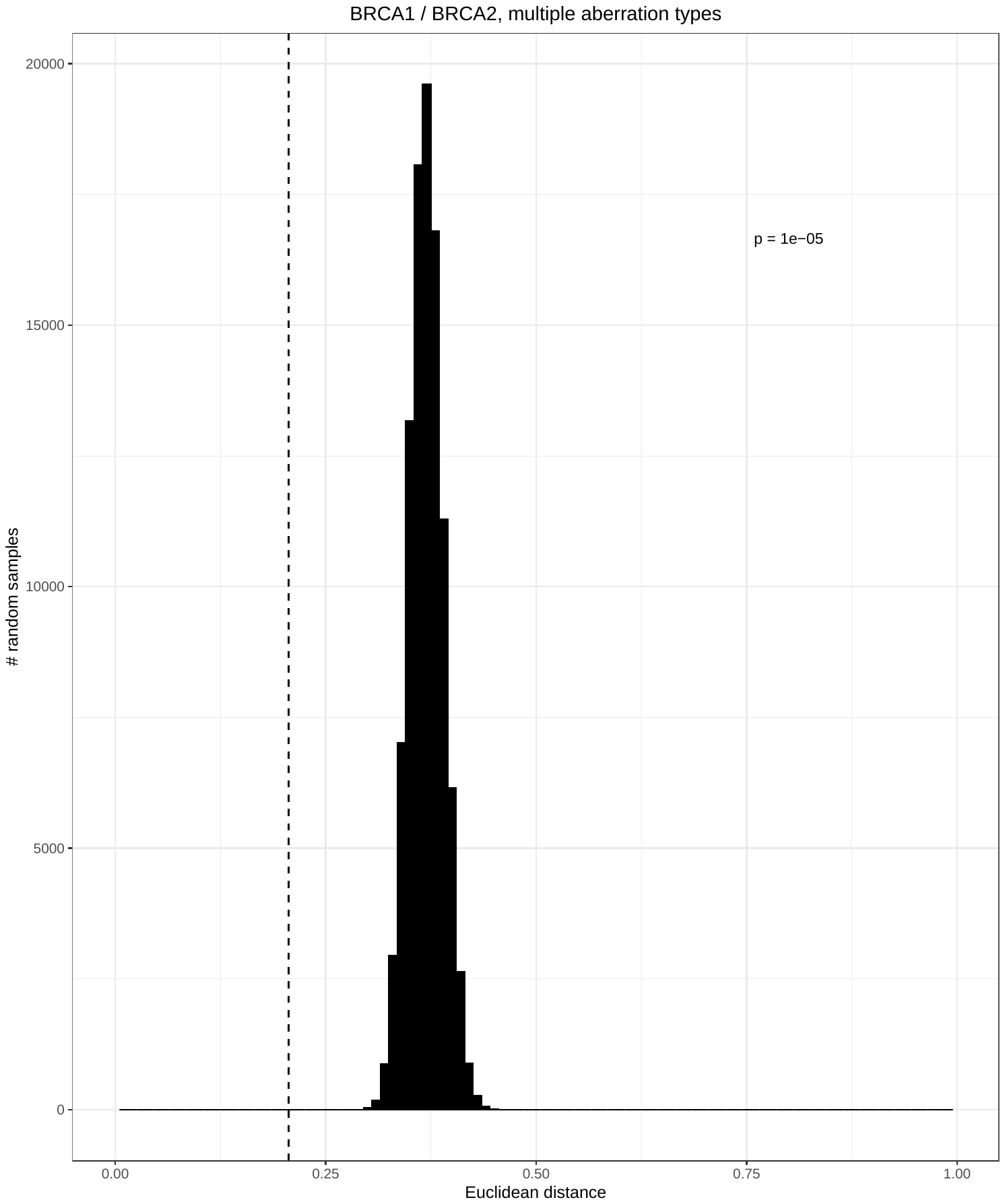
 **Figure S24: Euclidean distances for randomly selected patients compared to actual distances across all patients with a *BRCA1* or *BRCA2* aberration based on somatic-mutation signatures.**  We calculated the Euclidean distance between each pair of individuals who had a germline mutation, somatic mutation, homozygous deletion, and/or hypermethylation event in *BRCA1* and/or *BRCA2*; the median of these distances is illustrated using a vertical, dashed line. We then randomized the patient identifiers and calculated pairwise distances for the same number of randomly selected patients, which resulted in an empirical null distribution. We calculated a p-value by comparing the actual distance against the randomized distances.

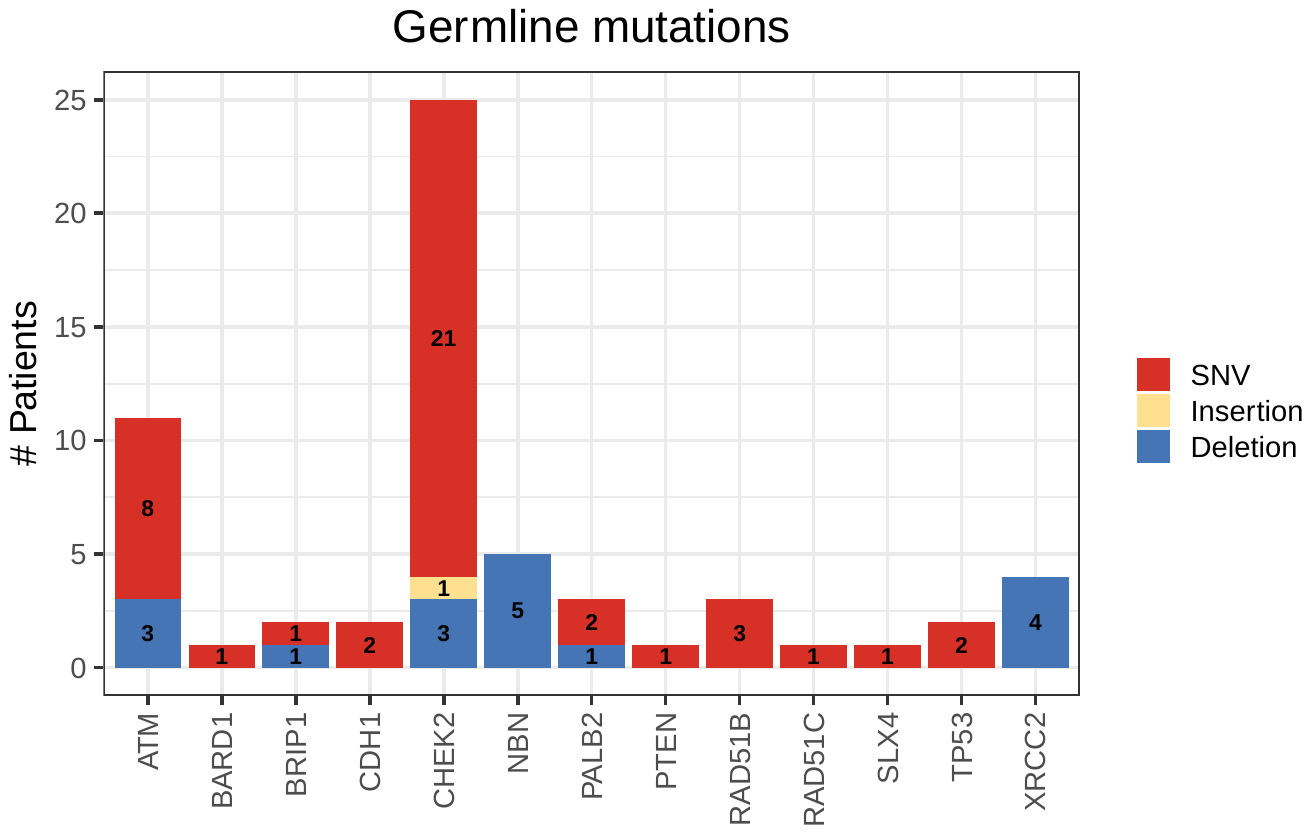
 **Figure S25: Number of patients with germline mutations in non-BRCA cancer-predisposition genes.**  This graph omits genes in which we observed no germline mutations. SNV = single-nucleotide variant.

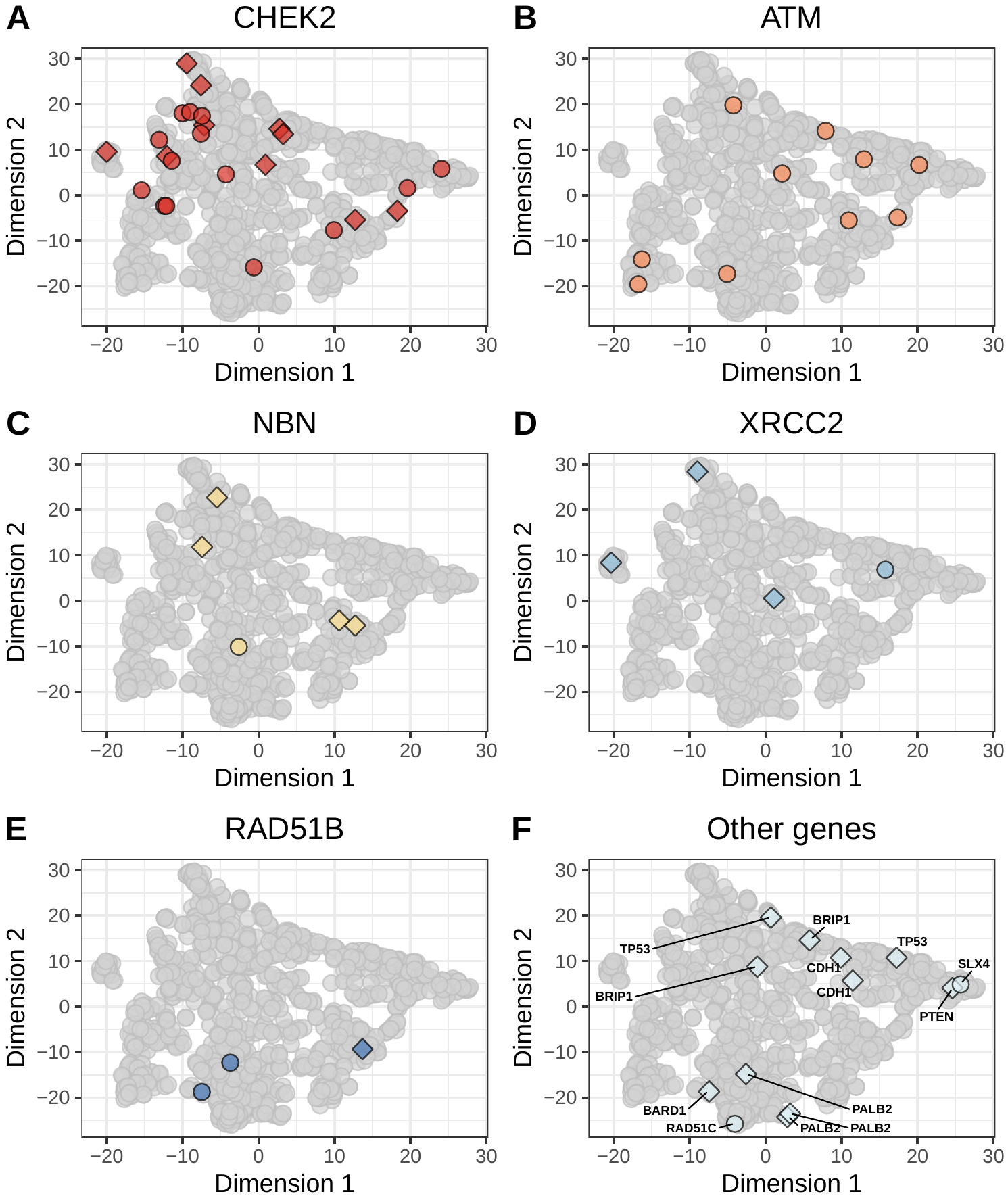
 **Figure S26: Non-BRCA germline mutations on the somatic-mutation signature landscape using the *t*-SNE method.**  Using the same two-dimensional representation of mutational signatures shown in Figure S13, this plot indicates which patients had germline mutations in non-BRCA cancer-predisposition genes. Diamond shapes indicate patients for whom *no* loss-of-heterozygosity was observed.

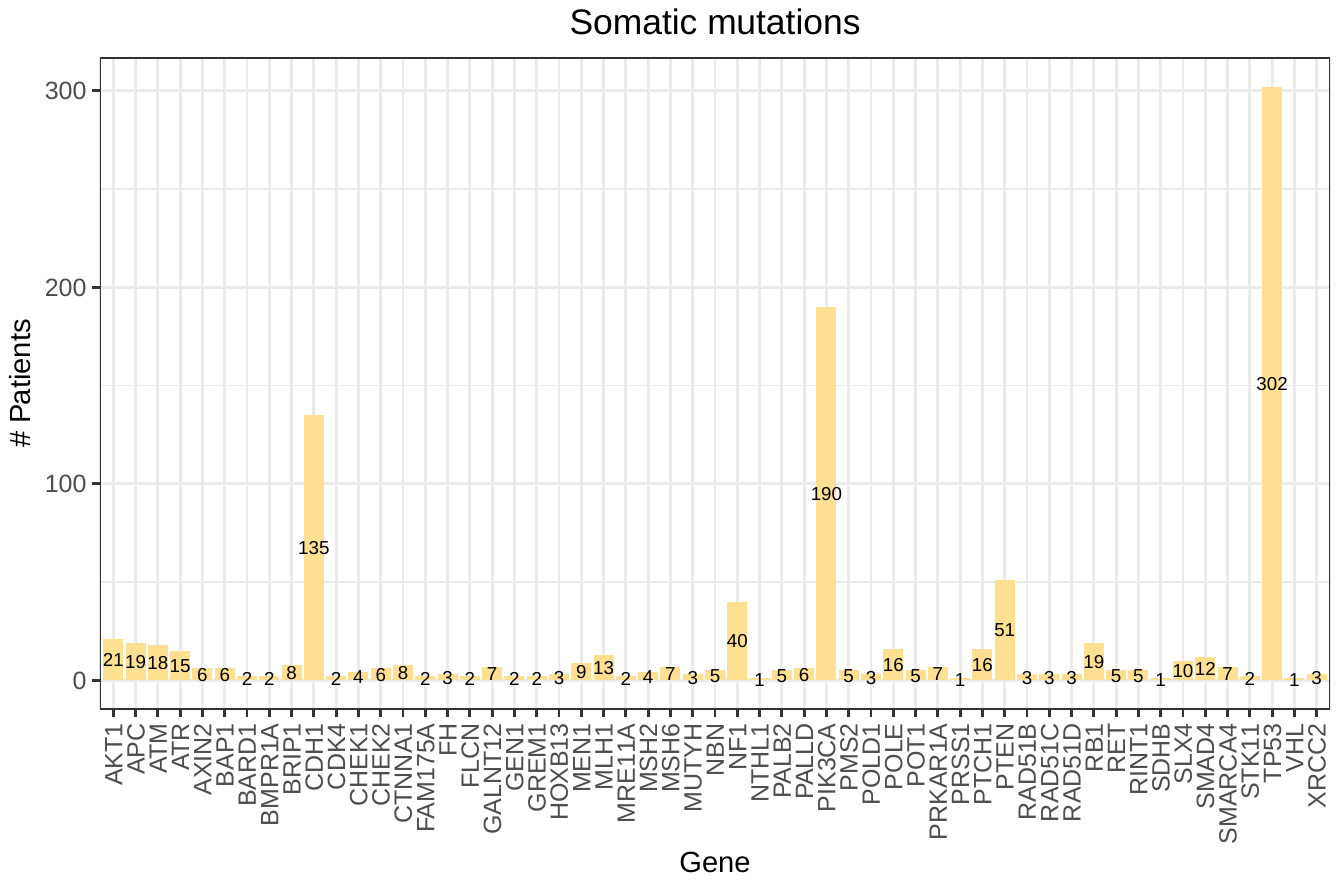
 **Figure S27: Number of patients with somatic mutations in non-BRCA cancer-predisposition genes.**  This graph omits genes in which we observed no somatic mutations.

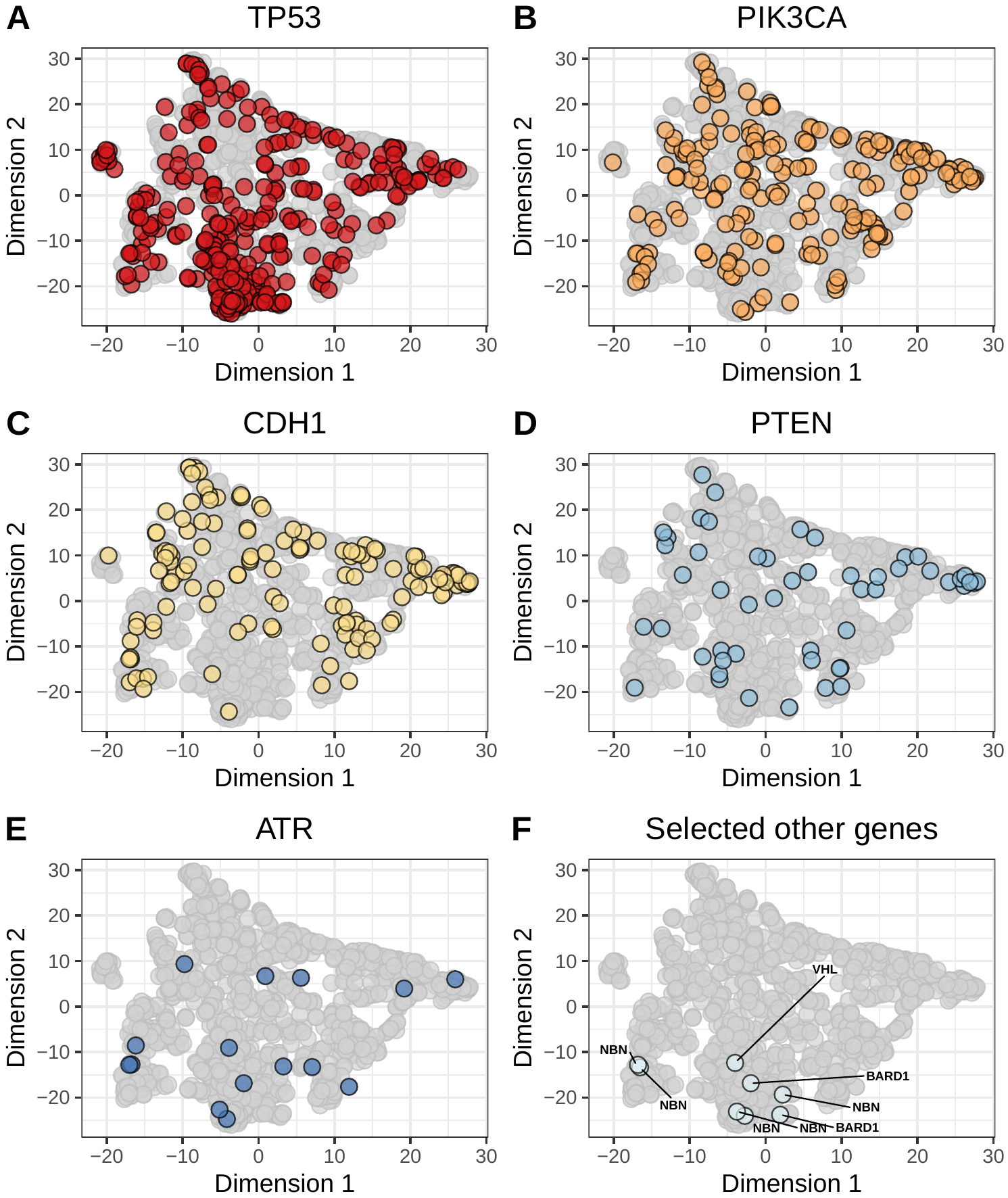
 **Figure S28: Non-BRCA somatic mutations on the somatic-mutation signature landscape using the *t*-SNE method.**  Using the same two-dimensional representation of mutational signatures shown in Figure S13, this plot indicates which patients had somatic mutations in non-BRCA cancer-predisposition genes. Diamond shapes indicate patients for whom *no* loss-of-heterozygosity was observed.

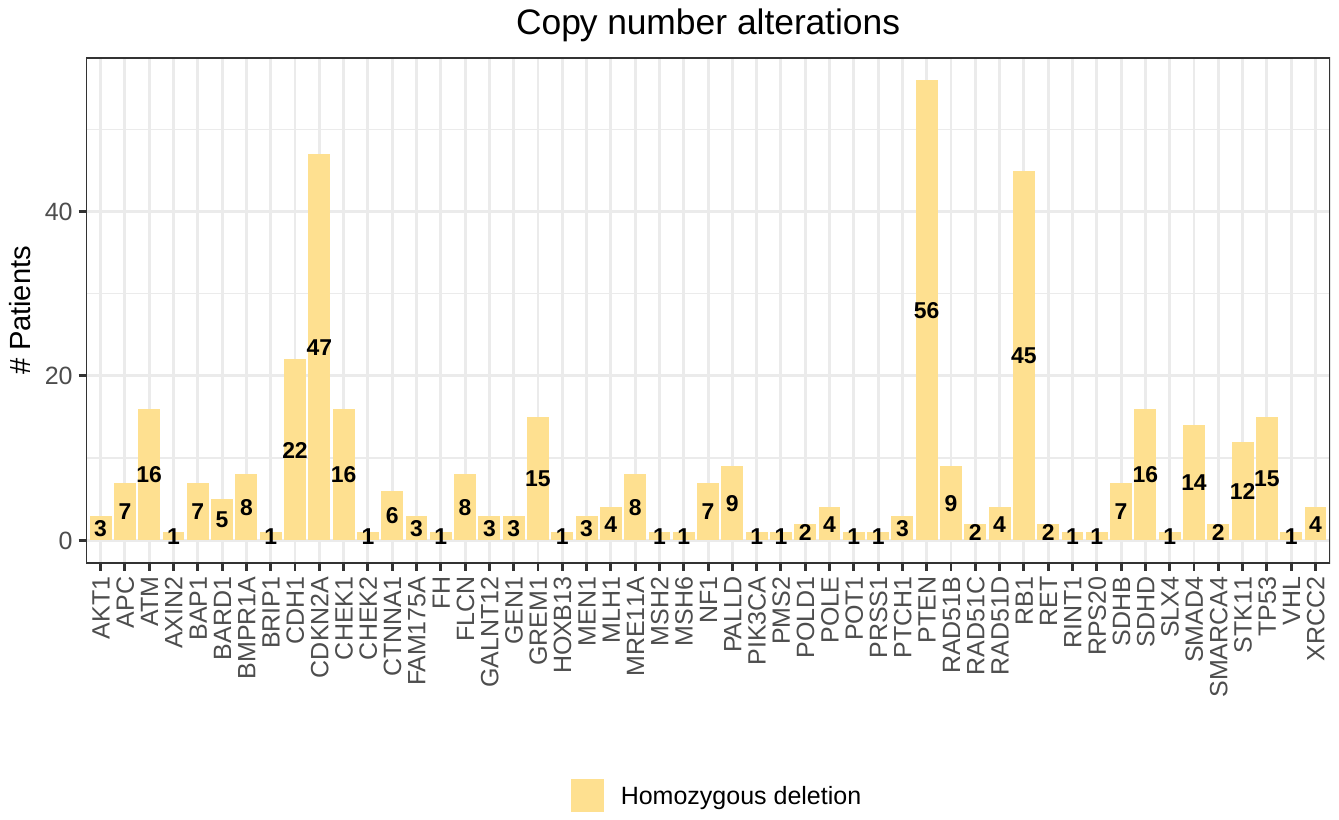
 **Figure S29: Number of patients with homozygous deletions in non-BRCA cancer-predisposition genes.**  This graph omits genes in which we observed no homozygous deletions.

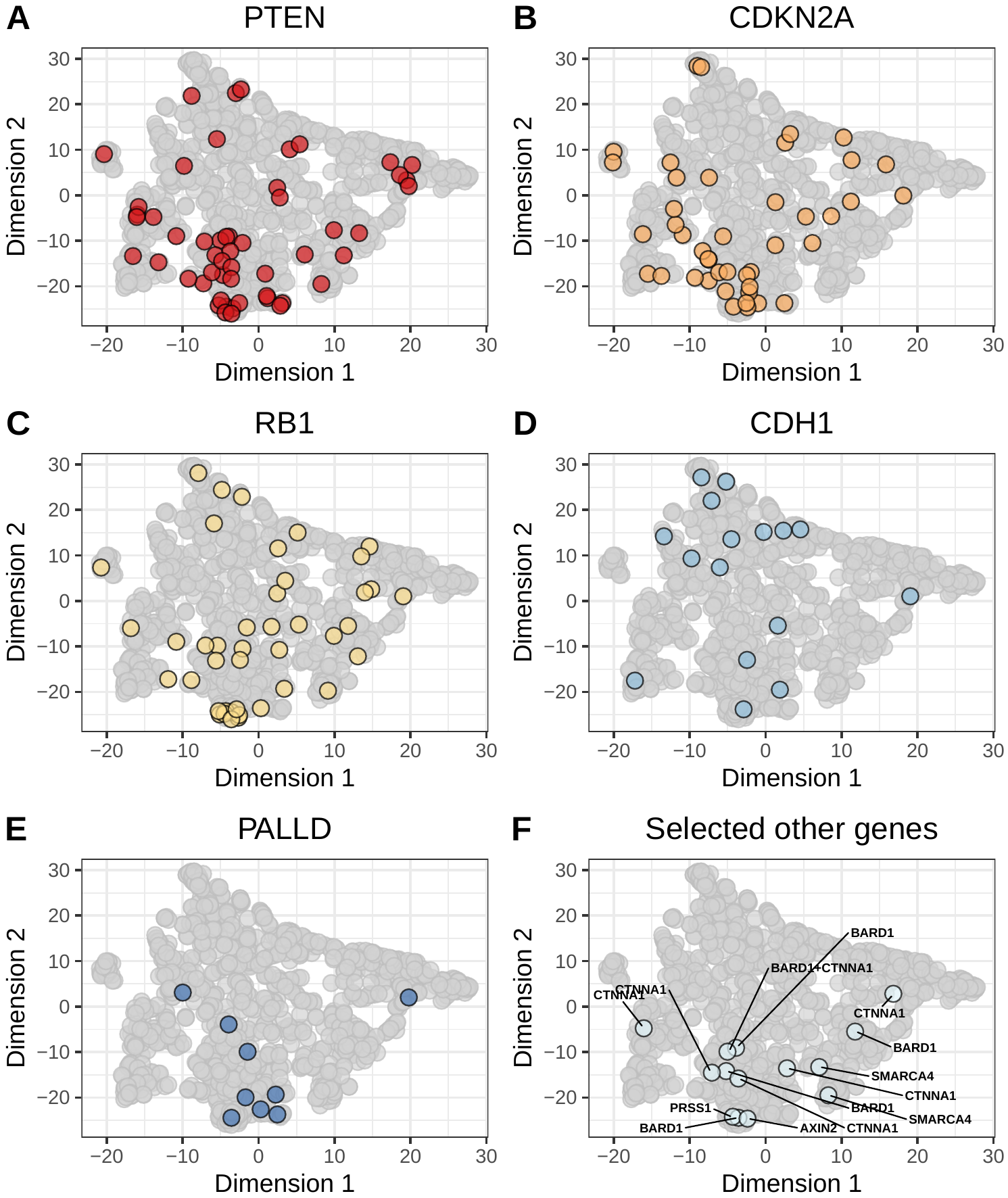
 **Figure S30: Non-BRCA homozygous deletions on the somatic-mutation signature landscape using the *t*-SNE method.**  Using the same two-dimensional representation of mutational signatures shown in Figure S13, this plot indicates which patients had homozygous deletions in non-BRCA cancer-predisposition genes. Diamond shapes indicate patients for whom *no* loss-of-heterozygosity was observed.

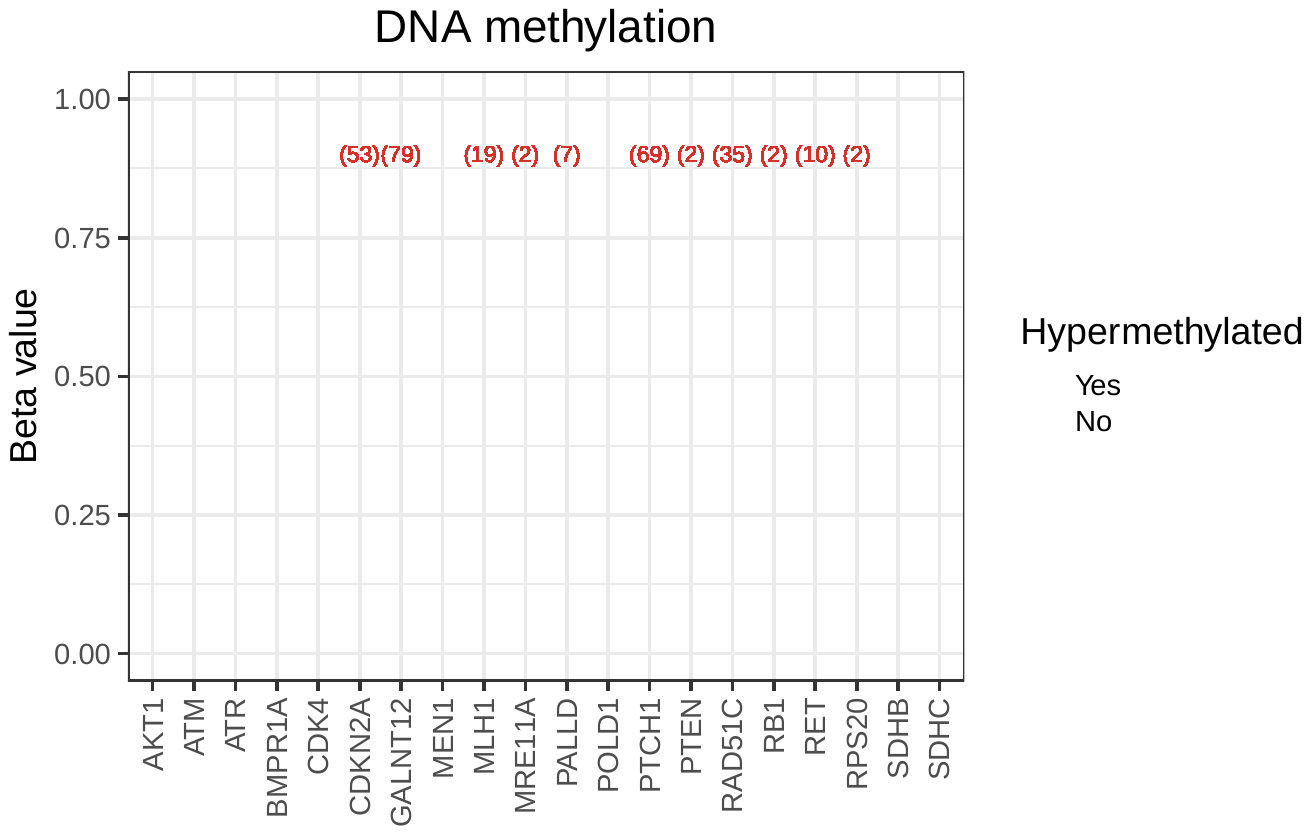
 **Figure S31: DNA methylation (beta) values for non-BRCA cancer-predisposition genes.**  Tumors that we classified as having hypermethylation events are highlighted as red points. This graph omits genes in which we observed no hypermethylation events.

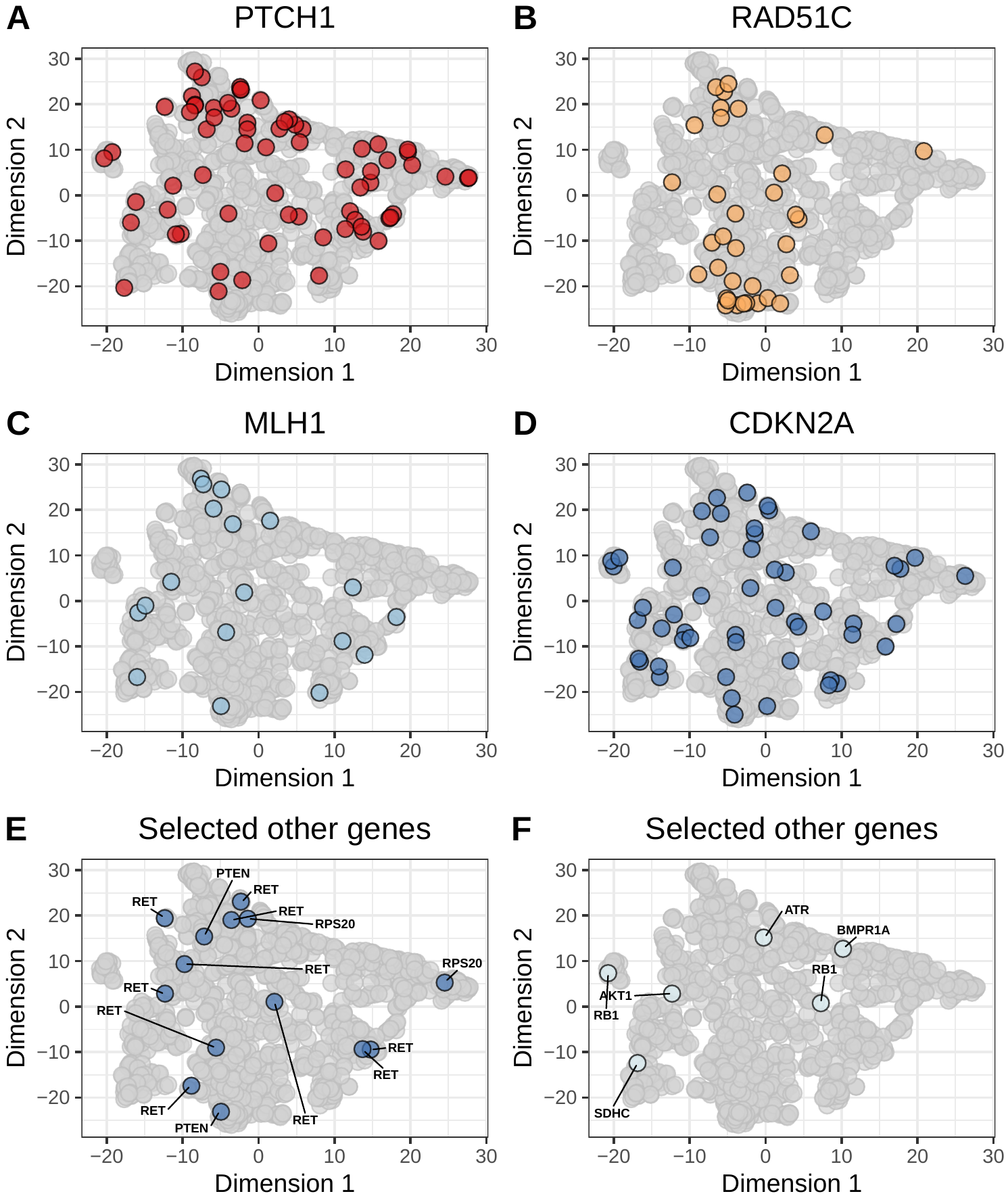
 **Figure S32: Non-BRCA hypermethylation events on the somatic-mutation signature landscape using the *t*-SNE method.**  Using the same two-dimensional representation of mutational signatures shown in Figure S13, this plot indicates which patients had hypermethylation events in non-BRCA cancer-predisposition genes. Diamond shapes indicate patients for whom *no* loss-of-heterozygosity was observed.

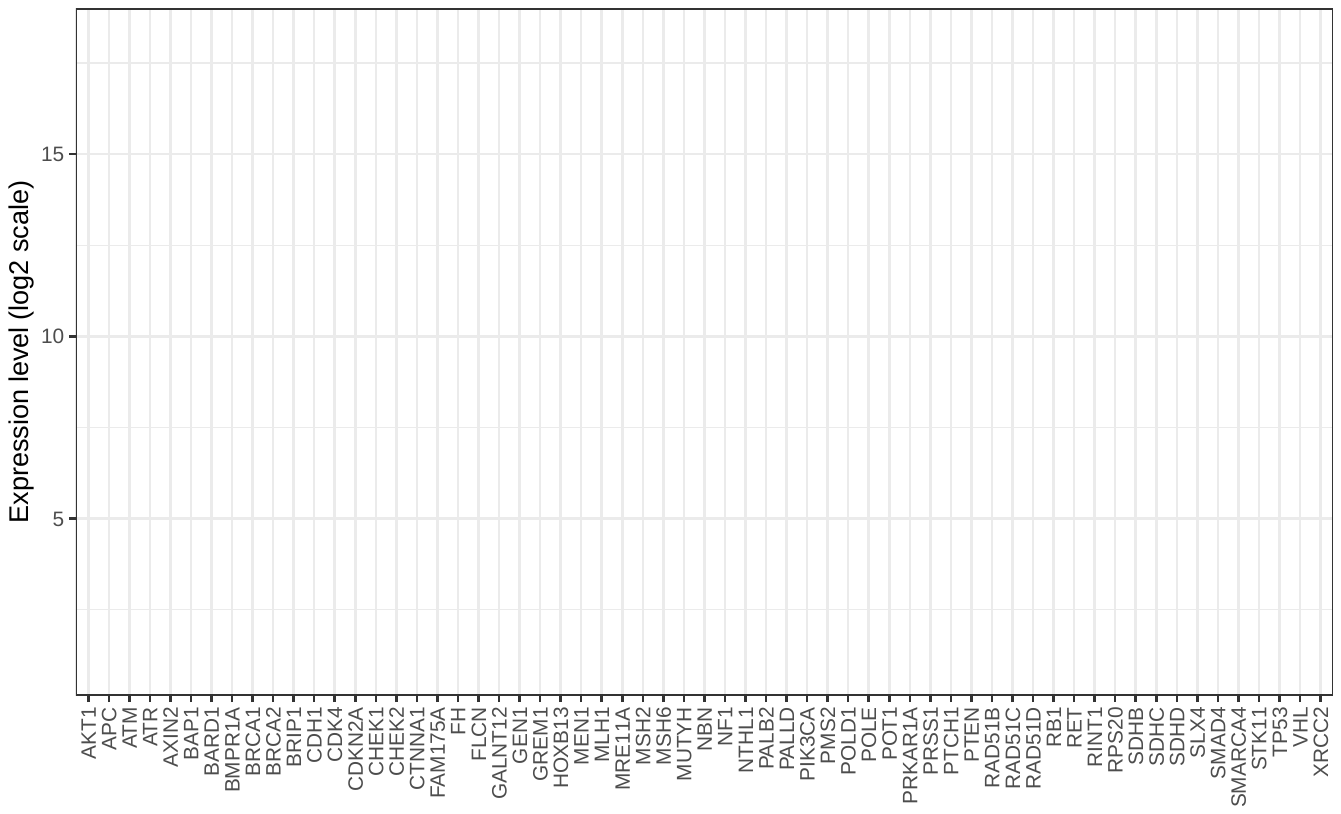
 **Figure S33: Gene-expression levels for all the genes we studied.**  For each gene, we identified tumors that expressed these genes at relatively low levels compared to other breast tumors; these low expressors are highlighted as red points.

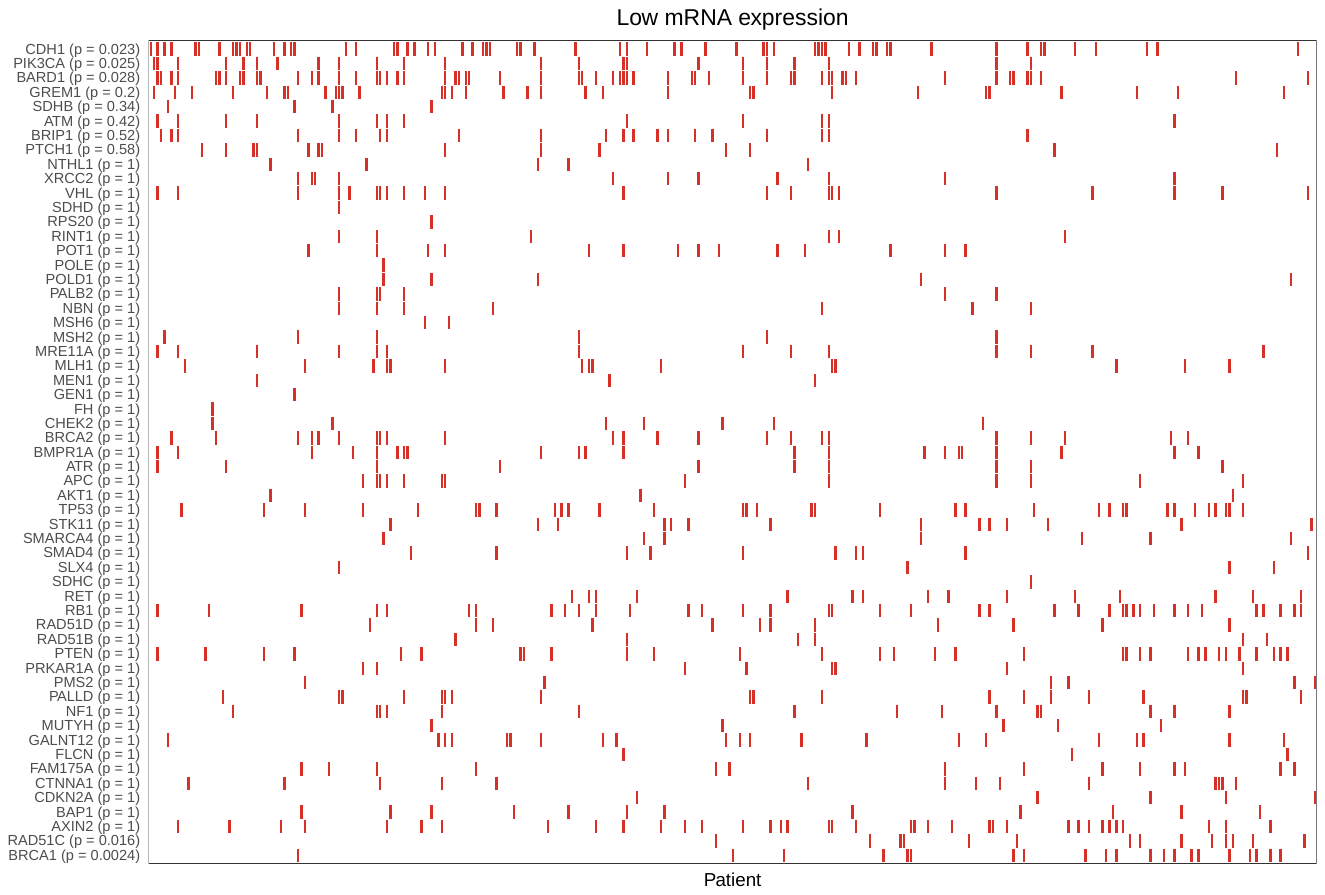
 **Figure S34: Relationship between BRCA aberration status and relatively low gene expression.**  We identified tumors with low expression for cancer-predisposition genes (see Figure S33) and evaluated whether the somatic-mutation signatures of these tumors were relatively similar or dissimilar to the BRCA reference group. Each red rectangle represents a patient sample that expressed a given gene at low levels. Low expression of *RAD51C* and *BRCA1* showed the strongest *positive* correlation between gene-expression status and the BRCAness reference group. Low expression of *BARD1* and *CDH1* showed the strongest *negative* correlation between gene-expression status and the BRCAness reference group. Genes for which no tumors exhibited low expression are omitted.

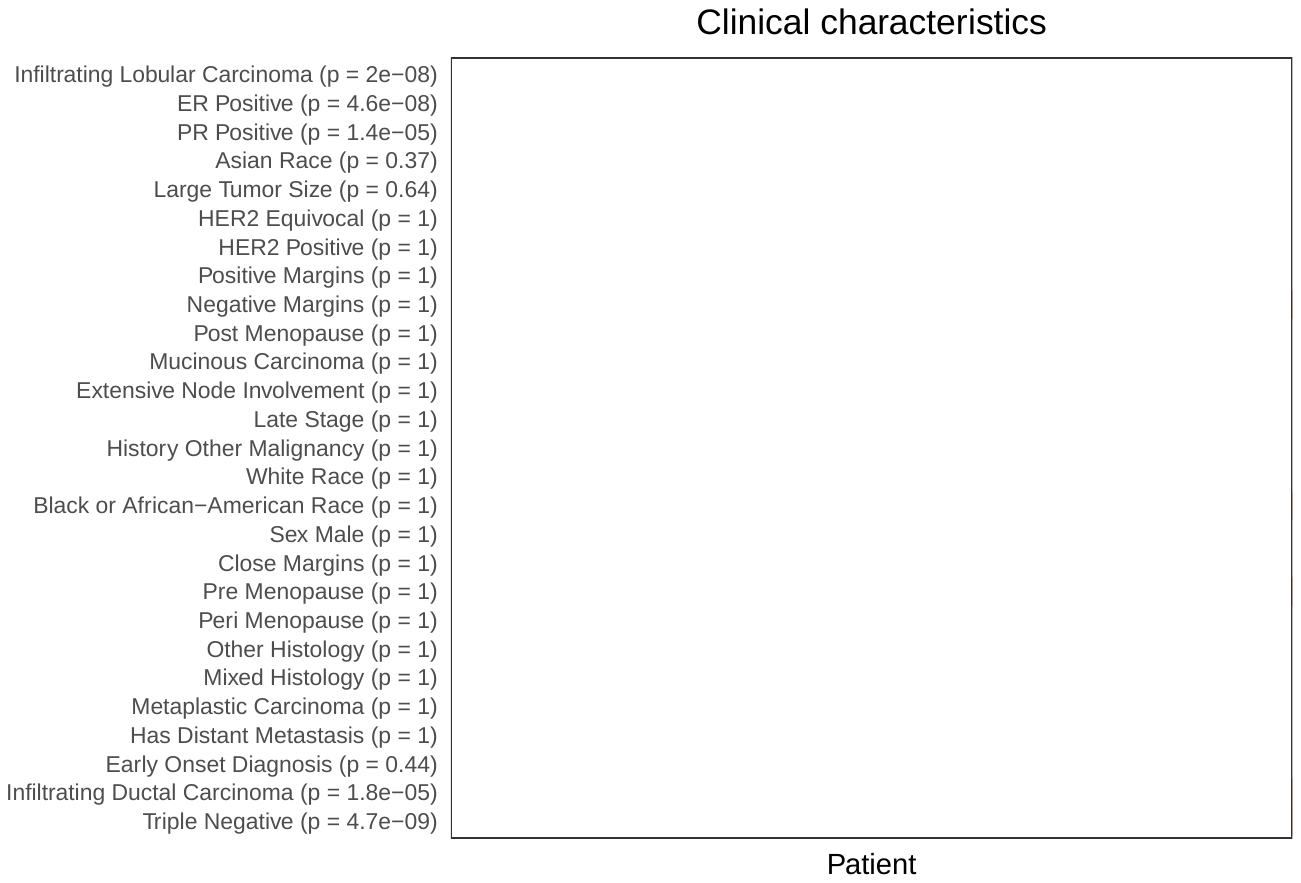
 **Figure S35: Relationship between BRCA aberration status and demographic, histopathological, and surgical observations in breast-cancer patients.**  Red rectangles indicate patients that were positive for each respective clinical characteristic. Tumors with triple-negative hormone receptors, infiltrating ductal carcinoma histologies, or close surgical margins overlapped most with BRCA-aberrant tumors based on somatic-mutation signatures.

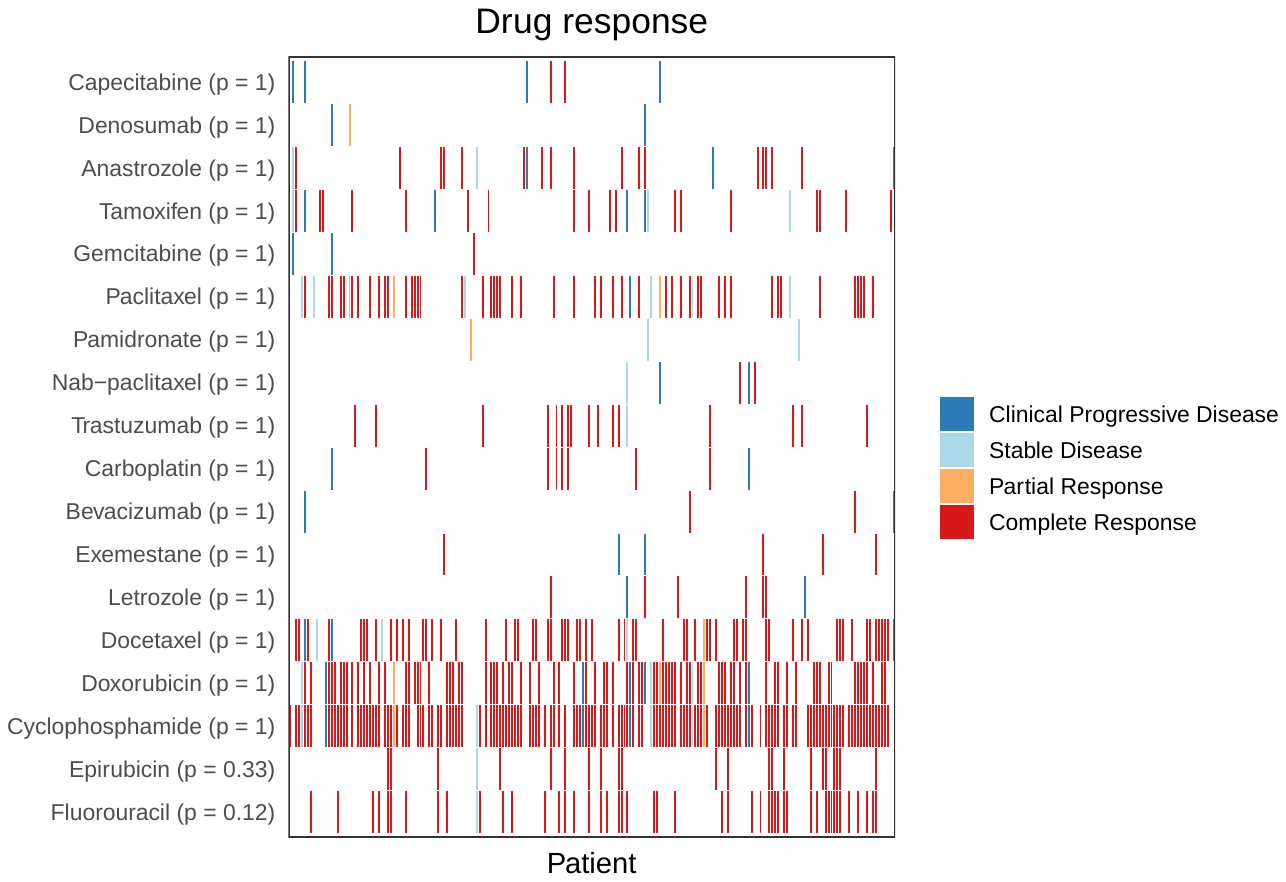
 **Figure S36: Relationship between BRCA aberration status and pharmacological responses in breast-cancer patients.**  We evaluated clinical treatment responses for 211 TCGA patients for whom drug-response data were available. Responses for none of the drugs were significantly correlated with BRCA aberration status based on somatic-mutation signatures.

### Supplementary Table

**Table S1: Summary of classification analysis for predicting a tumor’s aberration status.**  Via cross validation, we used gene-expression profiles and somatic-mutation signatures, respectively, to predict whether a given patient/tumor harbored particular types of aberrations. Sensitivity is equivalent to the true-positive rate. Specificity is equivalent to the true-negative rate. The area under the receiver operator characteristic curve (AUROC) quantifies the balance between sensitivity and specificity across a range of prediction thresholds.

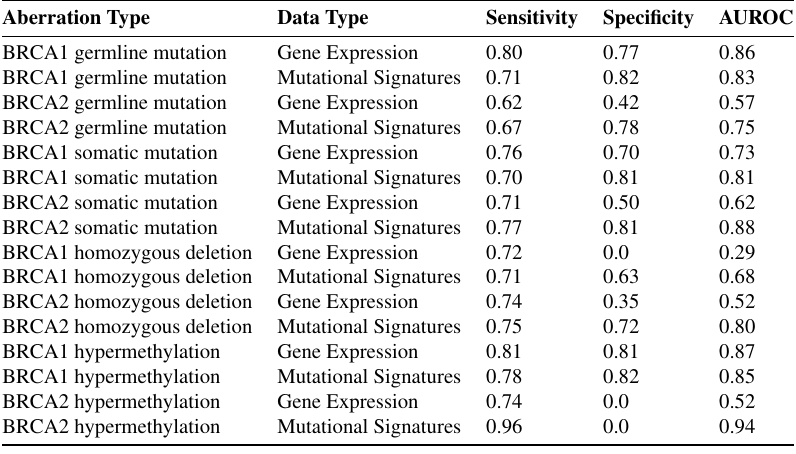
